# Alternative Splicing in TRPA1 Drives Sensory Adaptation to Electrophiles in Drosophilids

**DOI:** 10.1101/2025.05.09.653172

**Authors:** Hiromu C. Suzuki, Claire T. Saito, Srivarsha Rajshekar, Takaaki Sokabe, Diler Haji, Simon C. Groen, Julianne N. Peláez, Teruyuki Matsunaga, Ashleigh S. Takemoto, Kentaro M. Tanaka, Aya Takahashi, Makoto Tominaga, Shigeru Saito, Noah K. Whiteman

## Abstract

Behaviors are among the first traits to evolve as animals enter new niches, but their molecular bases are poorly understood. To address this gap, we used the mustard-feeding drosophilid fly *Scaptomyza flava*, which feeds on toxic Brassicales plants like wasabi that produce noxious, electrophilic isothiocyanates (ITCs or mustard oils). We found that *S. flava* exhibits dramatically reduced behavioral sensitivity to allyl isothiocyanate (AITC) compared to its microbe-feeding relatives *Scaptomyza pallida* and *Drosophila melanogaster*. We hypothesized that molecular evolution of the “wasabi receptor” TRPA1, known to detect ITCs in flies, could explain this loss of aversion. Our experiments revealed three interconnected evolutionary genetic changes consistent with this hypothesis. First, TRPA1 was expressed in labellar tissues of *S. flava* at the lowest levels among the three species, at a nearly four-fold lower level than in *D. melanogaster*. Second, *S. flava* expressed a higher proportion of TRPA1 splice variants previously reported to be less sensitive to chemical stimulus. Third, we identified amino acid substitutions in *S. flava* that could influence the structure of intracellular domains of TRPA1. To test the functional salience of these mechanisms *in vitro* and *in vivo*, we validated TRPA1 splicing isoforms using *Xenopus* oocyte electrophysiology and the *GAL4/UAS* system in *D. melanogaster*. Single TRPA1 isoform electrophysiology *in vitro* revealed evolution of the channel in the *S. flava* lineage towards reduced electrophile sensitivity. Ectopic expression of *S. flava* TRPA1 in *D. melanogaster* also consistently conferred weaker AITC sensitivity *in vivo* than expression of its orthologues, although this did not fully recapitulate differences in wild-type phenotypes between species, suggesting other molecular mechanisms were involved. To address this, we explored the consequences of isoform co-expression using oocyte electrophysiology. We found that enrichment of electrophile-insensitive TRPA1 splicing isoforms as observed in the salient *S. flava* sensory organs additively reduced cellular responses to AITC, which could further contribute to reduced electrophile aversion. Our findings illuminate how expression differences, protein structural changes, and especially alternative splicing, together can drive sensory evolution as animals behaviorally adapt to toxic new niches.

## INTRODUCTION

Animal behaviors are often encoded by complex genetic architectures. This mechanistic complexity has hampered efforts to link variation in genotype with variation in behavior (Bendesky and Bargmann 2011). An exception is progress in understanding how genetic changes in chemosensory genes expressed in the peripheral nervous systems of animals affect behavior. The relative simplicity and modularity of chemosensory functional units offers an appealing window for studying the molecular mechanisms underlying the evolution of behavior (Montell 2021; Valencia-Montoya et al. 2024).

Insects are useful models for addressing these questions because of their experimental tractability and an umwelt rich in volatiles and tastants. The vinegar fly family Drosophilidae, which includes the “fruit fly” *Drosophila melanogaster* and thousands of other species with diverse dietary niches, has played a central role in advancing our understanding of how animal behavior evolves through changes in chemosensation. Fueled by a wealth of genomic resources, genetic tools, and neuroanatomical insight, comparative analyses between *D. melanogaste*r and ecological specialists, such as noni-feeding *D. sechellia* (McBride 2007; McBride and Arguello 2007; Prieto-Godino et al. 2017; Auer et al. 2020; Álvarez-Ocaña et al. 2023), screw pine-feeding *D. erecta* (McBride and Arguello 2007), ripe fruit feeding *D. suzukii* (Karageorgi et al. 2017; Dweck et al. 2021; Wang et al. 2022), and cactus rot-feeding *D. mojavensis* (Diaz et al. 2018; Khallaf et al. 2020), have illuminated evolutionary genetic changes involved in the evolution of new chemosensory-driven behaviors. However, there remains a disconnect between our understanding of the specific molecular changes that give rise to these new behaviors, with notable exceptions (Prieto-Godino et al. 2017; Auer et al. 2020).

The fly genus *Scaptomyza*, an ecologically diverse group of 273 species nested within the Hawaiian *Drosophila* clade and sister to the subgenus *Drosophila* clade, provides another promising lineage to further investigate this problem. Of particular interest is the subgenus *Scaptomyza*, which includes obligate herbivorous species, such as *Scaptomyza flava* that attacks many mustard species in the Brassicaceae and Brassicales relatives (Whiteman et al. 2011). Unlike their generalist relatives in the other subgenera of *Scaptomyza* that feed on microbes within decaying plant tissues, *S. flava* larvae are obligate leafminers consuming internal cell layers (Aguilar et al. 2024) and adult females use leaf-cutting ovipositors to create feeding punctures in leaves and lay eggs (Whiteman et al. 2011; Peláez et al. 2022). Brassicales leaves are defended by both constitutive and inducible specialized metabolites, which impose a unique selective pressure on *S. flava* not present in microbial diets (Whiteman et al. 2011; Aguilar et al. 2024).

*S. flava* was reported as a leafminer of *Arabidopsis thaliana* in 1902 and can be maintained in the laboratory on this model plant, which allows for experimental manipulation of the host plant using an array of genetic tools (Whiteman et al. 2011). Herbivory evolved in the *Scaptomyza* lineage ca. 10-15 million years ago (Fig. 1A; Lapoint et al. 2013; Matsunaga et al. 2022; Peláez et al. 2022). This relatively recent dietary transition provides a useful context for identifying the evolutionary genetic changes accompanying this niche shift using a comparative approach. It is also an appealing model for studying the molecular basis of chemosensory evolution. Several host plant-related evolutionary changes in the chemosensory system of *S. flava* have been reported, including the lack of attraction to yeast odors and the gain of attraction to mustard-derived odorants, mediated by gene losses or duplications of respective olfactory receptor (*Or*) genes (Goldman-Huertas et al. 2015; Matsunaga et al. 2022).

**Figure 1.**
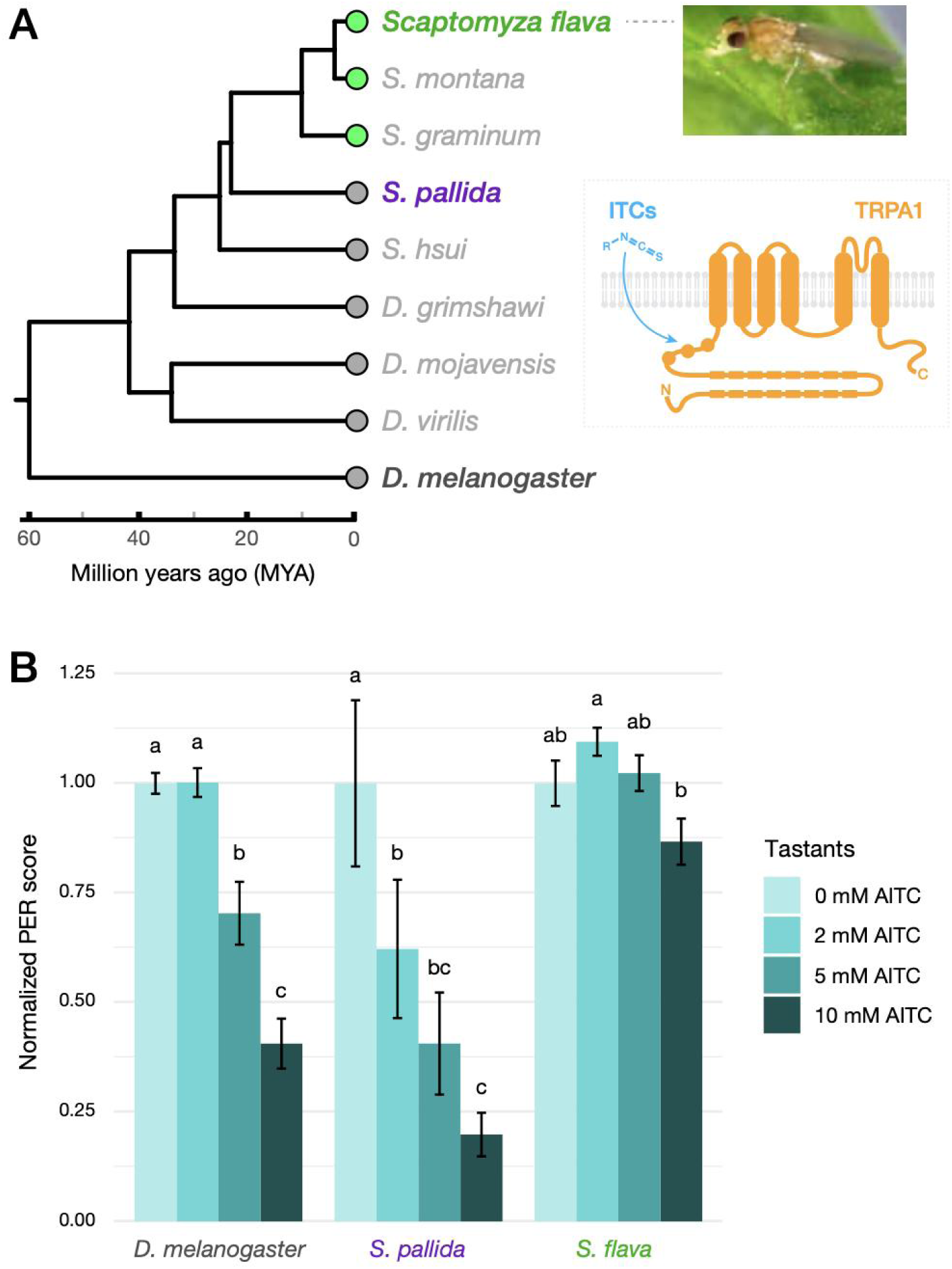
*Scaptomyza flava* exhibits attenuated feeding aversion against allyl isothiocyanate (AITC) (A) (Left) A time-calibrated species tree of drosophilids, including the genus *Scaptomyza*, based on 11 nuclear and mitochondrial genes (Matsunaga et al. 2022; Peláez et al. 2023). Colored circles at the tree tip represent the feeding ecology of the species (Grey: Microbivory, Green: Obligate herbivory). (Right) *Scaptomyza flava* female and a diagram of a TRPA1 channel targetted by isothiocyanates (ITCs). (B) Results of PER assay using adult females of the three focal species (*D. melanogaster*, *S. pallida*, and *S. flava*) at four different concentrations of AITC mixed in 300 mM sucrose solution. PER score of a single fly represents the feeding response to a given tastant over five trials, ranging between 0 to 5. Bar plots indicate individual PER scores for each treatment (n = 19-34 per treatment), normalized to the average score at 0 mM AITC within each species (see Method for details of data conversion). Letters indicate significant differences between tastants within species based on pairwise Wilcoxon rank-sum test with Benjamini-Hochberg correction (*p* < 0.05).

The major chemical defense system of mustard plants and their relatives in the Brassicales is known as the “mustard oil bomb.” During tissue damage, electrophilic, pain-causing toxins called isothiocyanates (ITCs), commonly known as mustard oils, can be formed through the hydrolysis of non-toxic β-thioglucoside N-hydroxysulfates called glucosinolates (GLSs). Specifically, when plant tissues are mechanically damaged by herbivores, GLSs are hydrolyzed by an enzyme called myrosinase and a subset of the GLS molecules are converted into ITCs, which are highly toxic owing to their electrophilicity and induce robust escaping behaviors in generalist insects, including *D. melanogaster* (Lichtenstein et al. 1964; Kang et al. 2010). Although other mustard-specialized insects like the cabbage white (*Pieris rapae*) can entirely suppress ITC formation by the action of lineage-specific proteins (Wittstock et al. 2004; Wheat et al. 2007; Okamura et al. 2022), *S. flava* allows ITCs to form during herbivory and mitigates mustard oil toxicity using the ancient mercapturic acid pathway (Gloss et al. 2014), which appears to have evolved adaptively under pressure from ITCs in *S. flava* (Gloss et al. 2019). Moreover, *S. flava* larvae develop more rapidly when reared on *A. thaliana* loss of function mutants that do not make GLSs required for the production of ITCs (Whiteman et al. 2012). Thus, *S. flava* encounters ITCs when attacking plants as adults and developing within them as immatures. Therefore, suppression of behavioral aversion against mustard oils is important for *S. flava* to successfully colonize the leaves of living mustard plants, yet complete ablation of ITC-sensing would also likely be maladaptive owing to the fact that development is slowed by the presence of dietary GLSs (Whiteman et al. 2012).

A candidate gene that may contribute to this behavioral change in *S. flava* is *TrpA1*, a member of the *Trp* (*Transient receptor potential*) channel gene family. TRPA1 is a polymodal cation channel, known as the “wasabi receptor”, activated by various sensory stimuli, including heat, UV radiation, reactive oxygen species, and algogens such as ITCs (Jordt et al. 2004; Talavera et al. 2020; Valencia-Montoya et al. 2024). Like most TRPA1 agonists, ITCs are electrophiles that form covalent bonds with several cysteine residues in the cytoplasmic domain of TRPA1, thereby changing the conformation of the entire channel (Hinman et al. 2006; Macpherson et al. 2007; Zhao et al. 2020). This mechanism-of-action is conserved from vertebrates to flies and planarians (Kang et al. 2010; Arenas et al. 2017). In adult *D. melanogaster*, TRPA1 channels serve a chemosensory function in bitter-sensing (*Gr66a*-expressing) gustatory neurons of the mouthparts (labellum and pharynx) and the legs (tarsi), triggering feeding aversion upon activation (Kang et al. 2010; Leung et al. 2020). Genetic knockout of *TrpA1* disrupts aversion to electrophiles, implying that the TRPA1 channel is the primary gustatory sensor for drosophilids to detect dietary electrophiles (Kang et al. 2010). Thus, the reduced gustatory aversion to ITCs in *Scaptomyza* flies could have evolved in part through molecular evolution of the TRPA1 channel.

Despite its deep conservation over evolutionary time, there are several cases wherein TRP channels are tied to dramatic sensory adaptations in other animals. A subspecies of the African mole-rat (*Cryptomys hottentotus pretoriae*) is insensitive to allyl isothiocyanate (AITC, the primary ITC in wasabi), hypothesized to be an adaptation that allows tolerance of venom components of ants found in their subterranean burrows (Eigenbrod et al. 2019). The lack of pain sensation is correlated with a reduced chemical sensitivity of TRPA1 due to the substitution of a key cysteine residue, providing a simple molecular mechanism for the behavioral change (Eigenbrod et al. 2019). Variation in heat activation thresholds of TRPA1 are also found among reptiles and amphibians in relation to species-specific thermal niches (Gracheva et al. 2010; Cordero-Morales et al. 2011; Akashi et al. 2018; Saito et al. 2022), suggesting that modification of the channel’s response properties might be a common strategy underlying ecological adaptation. A more complex example is found for another vertebrate heat sensor, TRPV1, in the vampire bat (*Desmodus rotundus*). This blood-feeder senses infrared signals from mammalian prey using pit organs. In a comparison with non-infrared-sensing bat species, TRPV1 of the vampire bat encoded two splicing isoforms with different threshold temperatures and this bat’s trigeminal neurons were highly enriched for a isoform with a truncated C-terminus that activates at a lower temperature than the a canonical isoform. The skewed expression ratio in favor of the splice isoform with higher heat sensitivity likely allows the vampire bat to sense the slightly elevated temperature in its prey that emit infrared (Gracheva et al. 2011).

We hypothesized that the *TrpA1* gene has also experienced adaptive evolution in the genus *Scaptomyza* owing to the novel chemical environment provided by the mustard host plants of *S. flava*, which produce ITCs upon tissue damage. Specifically, we addressed the working hypothesis that molecular evolution of TRPA1 facilitated specialization onto mustard host plants in the *S. flava* lineage by weakening gustatory aversion to ITCs. To this end, we conducted a behavioral screen followed by an in depth investigation of the characteristics of TRPA1 across three drosophilid species from different phylogenetic placements and with different diets: the mustard specialist herbivore *S. flava*, the closely related generalist microbivore *S. pallida*, and the rotting fruit and yeast-feeding *D. melanogaster*. Between the three focal species, we first compared gene-level and splicing isoform-level expression profiles of the *TrpA1* gene, and then investigated physiological and behavioral responses conferred by TRPA1 channel splicing isoforms against electrophiles in *Xenopus* oocyte and *D. melanogaster* rescue experiments, respectively. We found that reduced channel activity through alterations in gene expression, protein structure, and alternative splicing in TRPA1 are indeed key mechanisms correlated with adaptation to a specialized diet of mustard plants in *S. flava*. We conclude that incremental changes in multiple molecular mechanisms together can give rise to behavioral novelties.

## RESULTS

### Gustatory aversion of isothiocyanates is attenuated in *S. flava*

We first characterized feeding inhibition of flies in response to dietary AITC by monitoring PER (proboscis extension reflex) to tarsal/labellar stimulation (Shiraiwa and Carlson 2007; Kang et al. 2010). After a 24-hour starvation period, flies were challenged with a single concentration of AITC mixed in sucrose solution as test stimulus. Responses of individual flies were scored as a proxy of feeding inhibition by AITC and normalized to responses towards a control stimulus of 0 mM AITC, wherein a smaller score indicated stronger aversion (See Methods for details). This feeding assay showed that the two microbe-feeding species, *D. melanogaster* and *S. pallida*, clearly exhibited a stepwise reduction in feeding responses of female flies as the AITC concentration increased (Figure 1B). By contrast, females of the mustard specialist *S. flava* did not show significant feeding aversion towards AITC throughout the gradient, although we observed a slight reduction of PER at 10 mM AITC (approx. 13% lower than the 0 mM AITC; Figure 1B). This attenuated aversion towards AITC was also observed in male *S. flava* (Figure S1), despite the fact that they do not feed on plant exudates because they cannot pierce leaves, at least under laboratory conditions (Peláez et al. 2022). This suggests that while the behavioral aversion towards AITC in *S. flava* is not entirely lost, it is significantly weakened in both sexes of adults compared with its microbe-feeding relatives and consistent with our working hypothesis.

Given that food-associated odorants can promote feeding in *D. melanogaster* (Shiraiwa 2008; Oh et al. 2021), we also addressed whether and to what extent volatile ITCs from plants could enhance feeding in *S. flava* given that they are attracted to lower levels (Matsunaga et al. 2022), which would mitigate contact-mediated ITC aversion and might then explain the PER assay results. We repeated the PER assay in *S. flava* after surgically removing the antennae and maxillary palps to disrupt its olfactory inputs (Figure S1). We found that *S. flava* flies did not exhibit any significant increase or decrease of feeding rates based on AITC concentrations (Figure S1), in contrast to the slight aversion towards AITC observed in intact male flies (Figure S1). This pattern suggests that volatile ITCs – at least AITC – do not enhance feeding in *S. flava*.

**Supplementary Figure S1.**
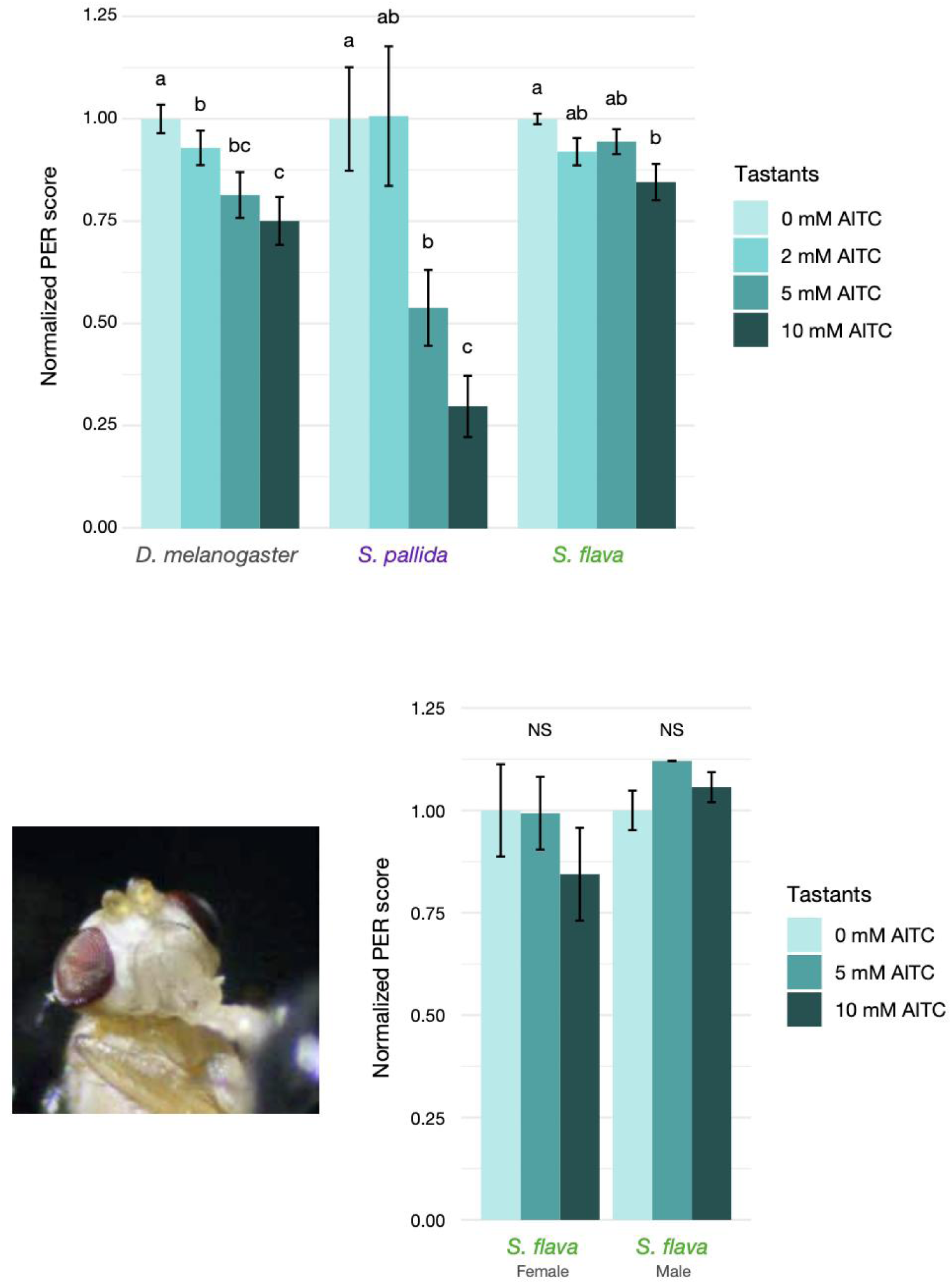
Additional data of PER assay. (Top) Results of PER assay using male flies from the three focal species (n = 20-32 per treatment). (Bottom left) A photograph of *S. flava* female with their antennal segments (3rd and 4th) and maxillary palps removed. (Bottom Right) Results of PER assay in antenna/palp-less *S. flava* (n = 9-14 per treatment). For both PER assays, individual PER scores were normalized to the average score at 0 mM AITC. Differences in PER scores between tastants within species or sexes were analyzed by pairwise Wilcoxon rank-sum test with Benjamini-Hochberg correction (*p* < 0.05).

Another hypothesis for these behavioral differences is that *S. flava* was hungrier than the other species after the starvation period prior to the assay, such that bitter sensation at the periphery was overridden by strong appetite (Inagaki et al. 2014; Yapici et al. 2016). To explore this possibility, we ran a simple hunger tolerance assay to monitor the survival rate of flies in foodless, humidified vials (Li et al. 2016). We found that hunger tolerance significantly varied between the species (Figure S2), and that the survival rates of *S. flava* were intermediate between those of *D. melanogaster* and *S. pallida* (Figure S2). In both sexes, *D. melanogaster* was most susceptible to the starvation treatment. *S. flava* and *S. pallida* were comparable in hunger tolerance up to 100-150 hours, but afterwards, *S. pallida* exceeded the tolerance of *S. flava* (Figure S2). The fact that the aversion towards AITC was clearly observed in the species at the two extremes of hunger tolerance (i.e., *D. melanogaster* and *S. pallida*), but not in *S. flava*, indicates that hunger does not explain the variation in AITC aversion across species. Taken together, we conclude that the weak aversion towards AITC in *S. flava* could be attributed to lineage-specific modifications of the sensory system, most likely in their gustatory neurons given that these neurons are responsible for mediating AITC detection through TRPA1.

**Supplementary Figure S2.**
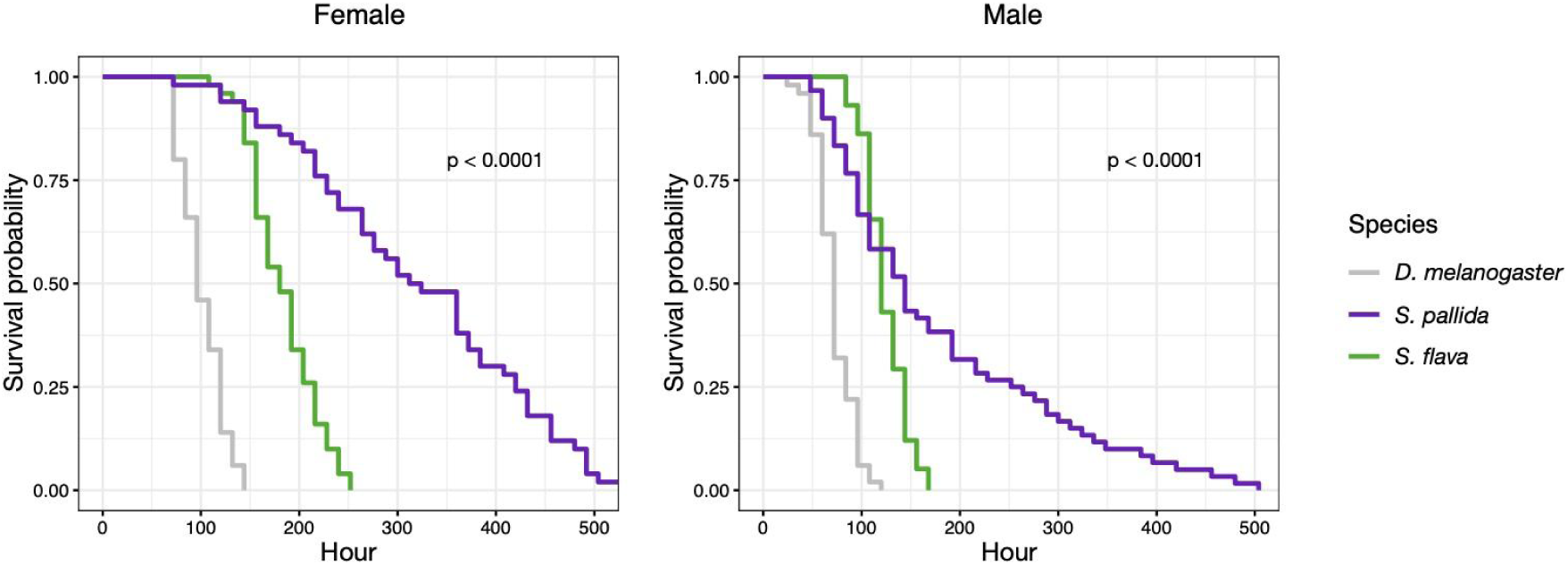
Hunger tolerance assay of adult flies. Survival plots of (Left) female and (Right) male flies, each of which consists of n = 5-6 independent groups of 8-10 flies. Differences between species were analyzed by log-rank test. Only group-wide *p*-values were shown.

### Cross-species variation in *TrpA1* gene expression

The *TrpA1* gene is intact in the *S. flava* genome and encodes an open reading frame. Bulk tissue RNA-sequencing confirmed robust expression of *TrpA1* in labella and tarsi of both *S. flava* sexes (Figures 2A-2F; Figure S3). Cross-species DEG analyses identified *TrpA1* as a differentially expressed gene in all comparisons but with varying degrees. For example, labellar *TrpA1* expression in *S. pallida* was only slightly higher than that in *S. flava* (Figures 2A and 2C), whereas there was almost four-fold higher expression in *D. melanogaster* compared to *S. flava* (Figures 2B and 2C). The ranks of *TrpA1* expression levels were reversed in tarsi, where *S. flava* showed the highest expression (Figures 2D-2F). This leg transcriptome analysis should be interpreted cautiously, however, because the *D. melanogaster* expression data were derived from both tarsi and tibia (Wang et al. 2022), while our *Scaptomyza* expression data were derived from tarsal segments alone. Nonetheless, these transcriptome patterns highlighted cross-species variation in *TrpA1* expression levels that could influence the differences in gustatory behaviors we observed.

**Figure 2.**
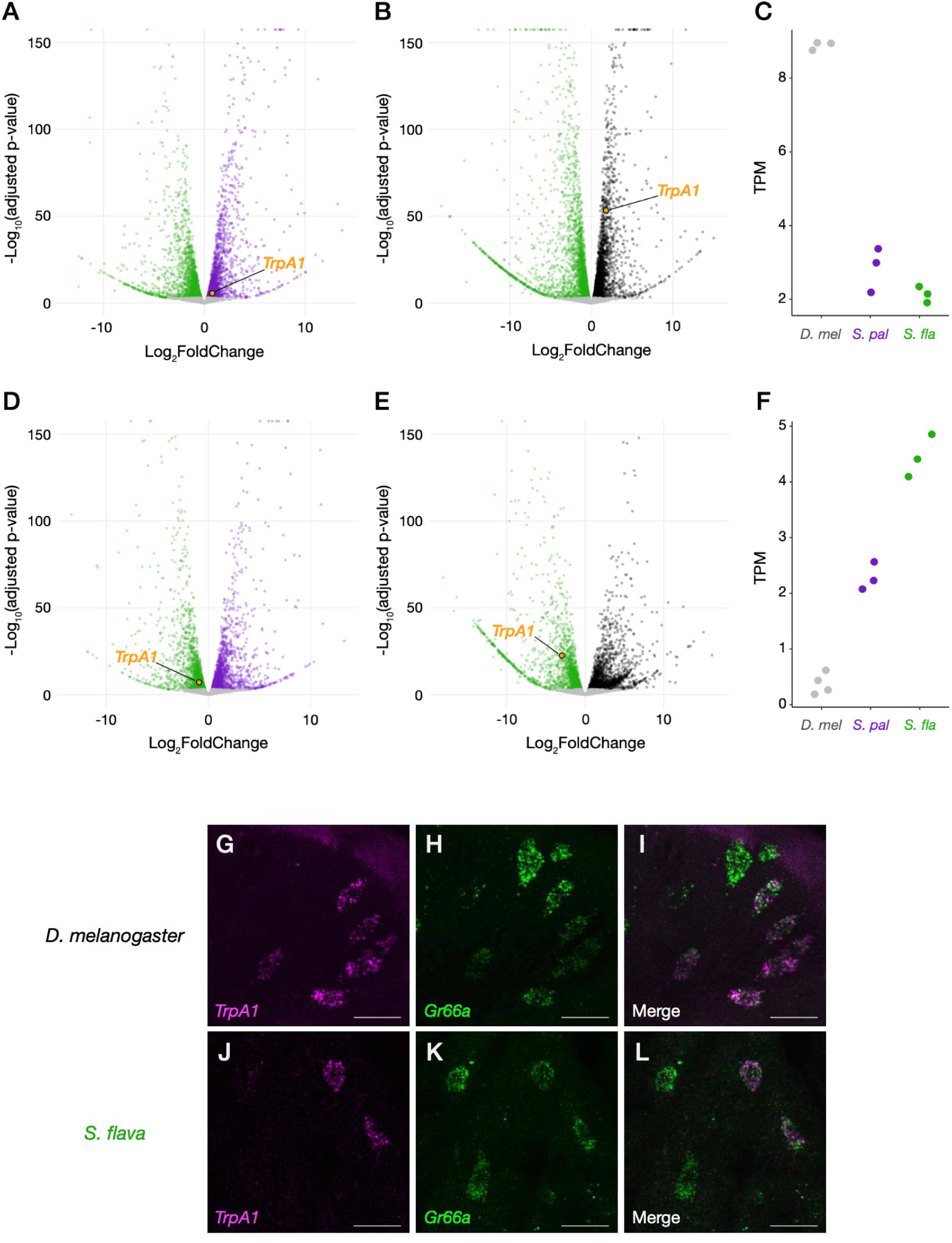
Gene expression of *TrpA1* is quantitatively variable but spatially conserved across drosophilids. (A, B) Volcano plots of cross-species DEG analysis using female labellar (mouth) transcriptome between (A) *S. flava* - *S. pallida*, and (B) *S. flava - D. melanogaster*. Colored dots represent differentially expressed genes between species (adjusted *p*-value < 0.001). (C) TPM (transcripts per million) values from labellar transcriptomes (n = 3 per species). (D, E) Volcano plots of cross-species DEG analysis using female tarsal (leg) transcriptome between (D) *S. flava* - *S. pallida*, and (E) *S. flava - D. melanogaster*. Colored dots represent differentially expressed genes between species (adjusted *p*-value < 0.001). (F) TPM values from tarsal transcriptome (n = 3-4). Note that *D. melanogaster* transcriptome data used here were obtained from (Dweck et al. 2021) and (Wang et al. 2022). (G-L) Confocal images of labella from (G-I) *D. melanogaster* female and (J-L) *S. flava* female, showing fluorescent signals from *TrpA1* (magenta; G, J), *Gr66a* (green; H, K), and merged mRNAs (I, L). Scale bars represent 10 µm.

Changes in chemosensory preferences could also be induced by receptor misexpression among sensory neurons. To characterize the types of neurons expressing TRPA1 channels, we visualized mRNA localization in the labellum of *S. flava* and *D. melanogaster* using HCR (hybridization chain reaction) RNA-FISH (Choi et al. 2018). We found in *S. flava* that signals of *TrpA1* mRNA formed neuron-like clusters and always co-localized with those of *Gr66a*, a marker of bitter-sensing gustatory neurons (Figure 2J-2L). This pattern was also consistent with our HCR images of *D. melanogaster* labella (Figure 2G-2I), as well as those from a previous study (Leung et al. 2020). These results showed that *TrpA1* expression in bitter-sensing gustatory neurons are conserved between the two lineages, indicating *S. flava* still perceives ITCs as deterrents despite their mustard-feeding ecology.

**Supplementary Figure S3.**
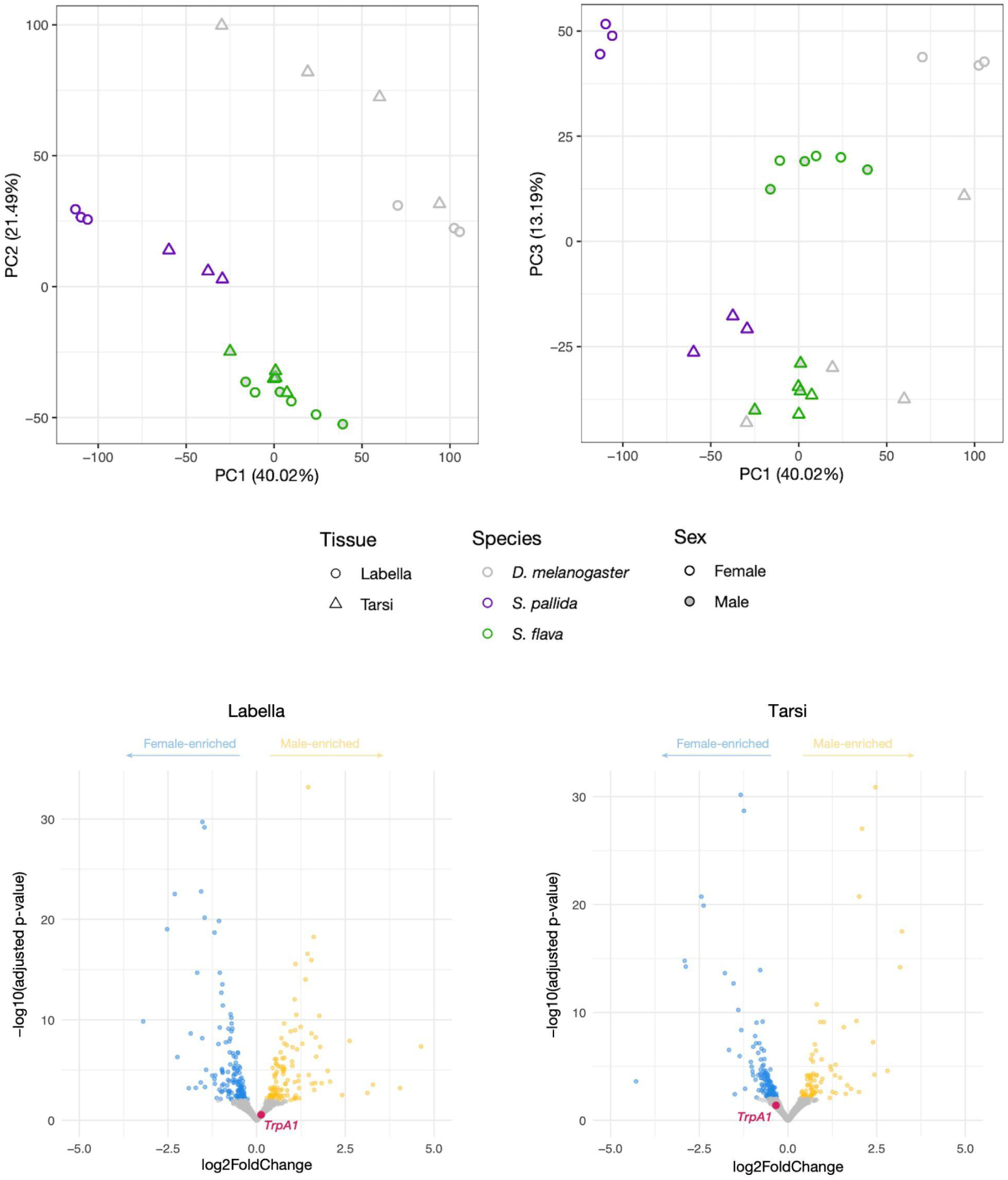
Labellar and tarsal transcriptome analyses. (Top) PCA plots for 25 transcriptome datasets based on read count data of 8,920 orthologous genes. (Bottom) Volcano plots of DEG analysis between female and male *S. flava*. *TrpA1* was not differentially expressed between sexes in both labella and tarsi.

### Compositions of *TrpA1* splicing isoforms vary predictably across species

The *D. melanogaster TrpA1* gene undergoes alternative splicing to produce five isoforms with variable expression and functional properties (Kang et al. 2012; Zhong et al. 2012; Gu et al. 2019; Leung et al. 2020; Gu et al. 2022; Sato et al. 2023; Du et al. 2024). These differences in TRPA1 splicing isoform function could potentially also be exploited during adaptation to dietary electrophiles as a source of sensory innovation in *S. flava* as in the case of the truncated TRPV1 splicing isoform that evolved in infrared-sensing vampire bats (Gracheva et al. 2011).

To this end, we investigated the structures and quantified the compositions of *TrpA1* splicing isoform mRNAs across *Drosophila/Scaptomyza* species and across the salient gustatory organs. Initial *de novo* assembly of RNA-sequencing reads revealed the presence of at least five *TrpA1* splicing isoforms expressed in the labella/tarsi of *S. flava* and *S. pallida* (Figure 3A). All *Scaptomyza TrpA1* splice variants contained Exons 1 and 2, showing similarities to the *TrpA1-C/D/E* splicing isoforms of *D. melanogaster* (Figure 3A). We found negligible expression of *TrpA1-A/B* analogues in *S. flava* labella and tarsi, and little expression of these in *S. pallida*, although the first exonic sequence (i.e., Exon 3) was intact in both genomes (Figure S4). Intriguingly, all *Scaptomyza* spp. *TrpA1s* acquired two additional exons upstream of Exon 1, named Exon L and Exon S (“Longer” or “Shorter” exons), each of which provided a putative start codon for different splicing isoform types (Figure 3A; Figures S4 and S5). Genomic comparisons revealed that Exon L appeared at least in the common ancestor of the subgenus *Drosophila* and retained some sequence similarity to the 5’ UTR of *D. melanogaster TrpA1*, implying a potentially ancient evolutionary origin of this motif (Figures S4 and S6). Exon S likely evolved at the base of the Hawaiian *Drosophila* and *Scaptomyza* clades, although we lacked transcriptional data for most species (Figures S4 and S6). These newly-discovered *Scaptomyza TrpA1* splicing isoforms were named *TrpA1-CL/CS/DL/DS/EL*, respectively, based on the choice of exons and their homology to *D. melanogaster TrpA1* splicing isoforms (Figure 3A; adopting the nomenclature from Zhong et al. 2012). The presence of these splicing isoforms was further validated by subcloning and Sanger sequencing of plasmids.

**Figure 3.**
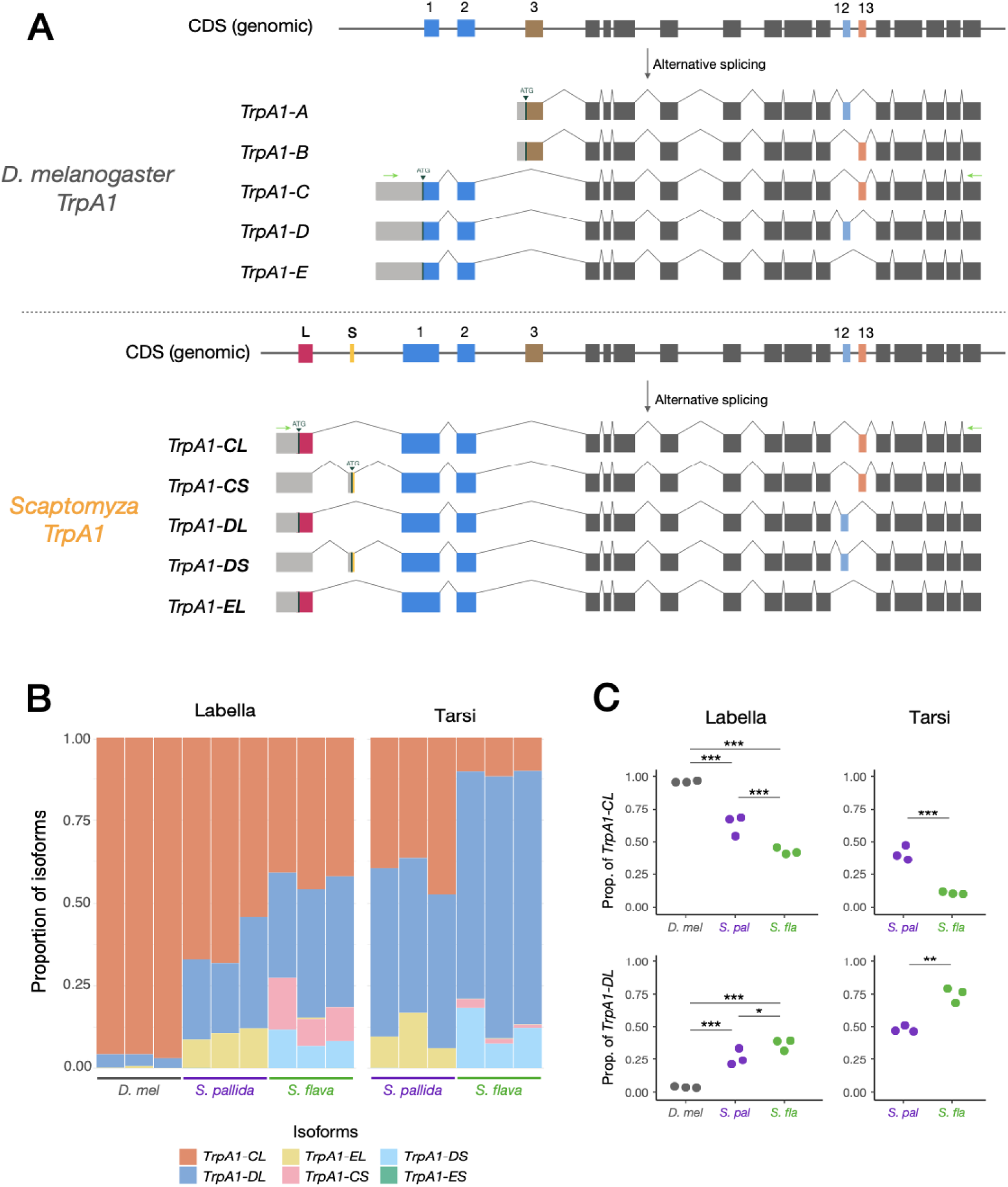
Compositions of *TrpA1* splicing isoforms vary among species. (A) Exon-intron structures of *TrpA1* splicing isoforms of *D. melanogaster* and *Scaptomyza* (i.e., those identified from *S. flava* and *S. pallida*). Small arrowheads indicate putative translation start sites, whose upstream are putative 5’UTR sequences (grey). Green horizontal arrows indicate the positions of PCR primers designed for amplicon sequencing (see Figure S7). (B) Compositions of *TrpA1* isoforms from female labelar and tarsal cDNA of the three focal species. Each vertical bar represents a single cDNA sample derived from an independent set of 40-50 flies. Note that *D. melanogaster TrpA1-C/D/E* splicing isoforms here were categorized as *TrpA1-CL/DL/EL*, respectively, because of the sequence similarity between its 5’UTR region and Exon L of *Scaptomyza TrpA1s* (see also Figures S4 and S6). *TrpA1-ES* was not detected in transcriptomes of any of the species but included as a potential isoform. (C) Relative abundances of *TrpA1-CL* and -*DL* in labellar and tarsal cDNA derived from individual sets of 40-50 flies (n = 3 per species). Variation in splicing isoform compositions among species was modeled by GLM with Gaussian distribution. Labellar compositions were further tested by Tukey’s post-hoc test with Benjamini-Hochberg correction. *: *p* < 0.05, **: *p* < 0.01, ***: *p* < 0.001.

Quantification of splicing isoform expression posed a challenge in the case of *Scaptomyza TrpA1*. Because exons that define splicing isoforms are distantly located on mRNAs (i.e., Exons L/S and 12/13 are approximately 2 kb apart; Figure 3A), we could not accurately parse out the expression of each splicing isoform using short-read RNA-sequencing or quantitative PCR. To address this issue, we carried out amplicon sequencing of *TrpA1* transcripts using a MinION long-read sequencer (Oxford Nanopore Technologies) that covers up to 4 Mb in a single read. Briefly, we PCR-amplified the entire repertoire of *TrpA1* transcripts from labellar or tarsal cDNA (except for *TrpA1-A/B*), sequenced them on the MinION, and sorted each sequence read into a closest isoform type using BLAST searching, assuming isoform compositions were maintained throughout the process (Clark et al. 2020; Figure S7). This approach revealed substantial variation in the composition of *TrpA1* splicing isoform expression across species and organs (Figure 3B), except for *D. melanogaster* tarsi, which failed the cDNA recovery due to low *TrpA1* expression levels (Figure 2F). Notably, while the *TrpA1-CL* (*TrpA1-C*) splicing isoform was predominant in *D. melanogaster* and enriched in *S. pallida* labella, *TrpA1-DL* was more represented in *S. flava* labella (Figures 3B and 3C). Differences between tissues were also striking, wherein *TrpA1*-*DL* tended to be more abundant in tarsi than in labella (Figure 3B and 3C). Isoforms containing Exon S, *TrpA1-CS* and -*DS*, were almost exclusively detected in *S. flava,* while the expression of *TrpA1-EL* was only visible in *S. pallida* (Figure 3B; Figure S8). These unique proportions of *TrpA1* splicing isoforms suggest that the differences in splicing isoform compositions might be linked to the behavioral variation across fly species.

**Supplementary Figure S4.**
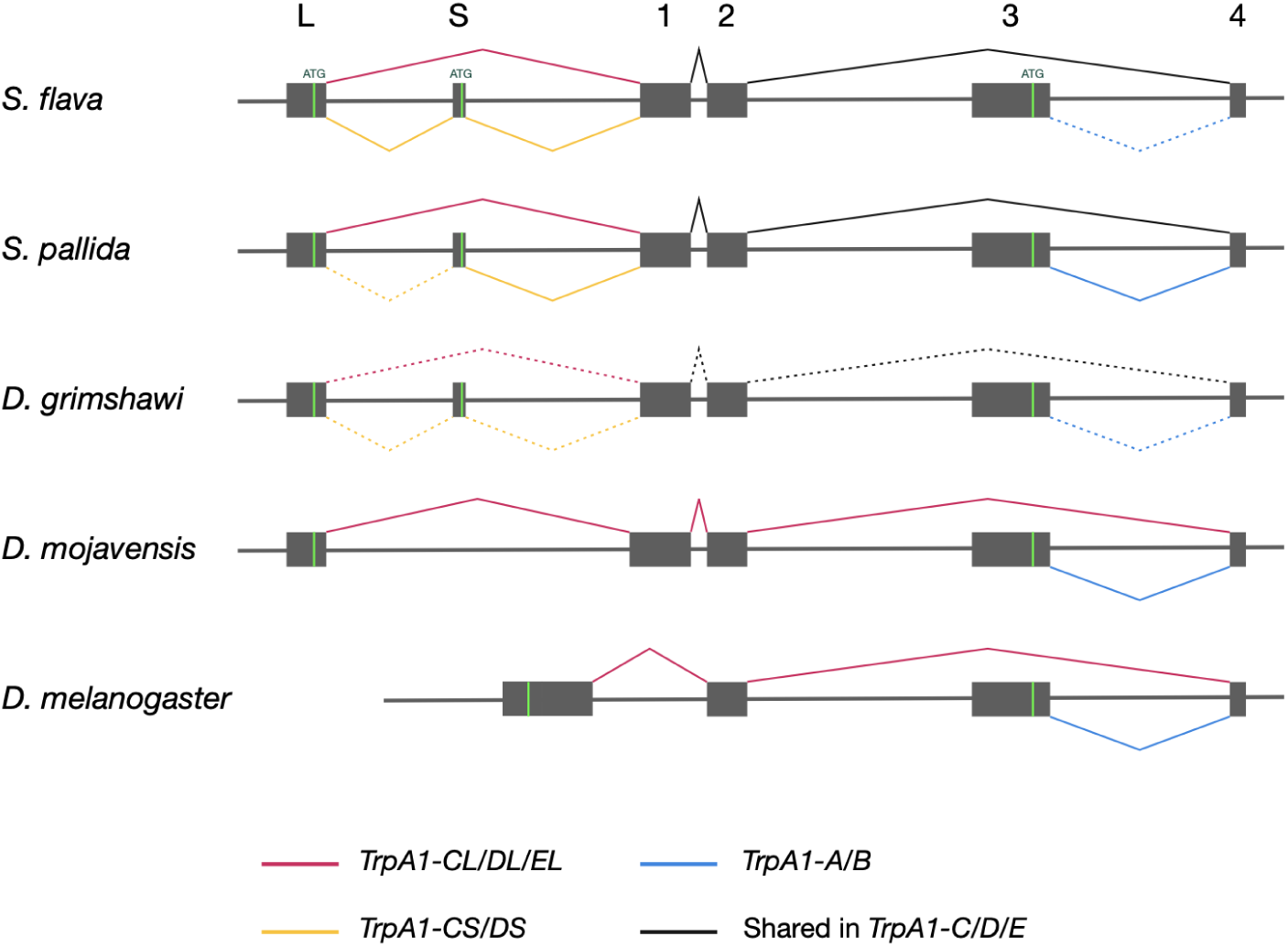
Evolution of N-terminal exons of *TrpA1* across drosophilids. Colored lines indicate exon-intron boundaries specific to each isoform type. Dashed lines show the lack of transcriptional evidence (inferred only from genomic/CDS-level information). Green vertical lines indicate putative (i.e., the most upstream) translation start sites for each isoform.

**Supplementary Figure S5.**
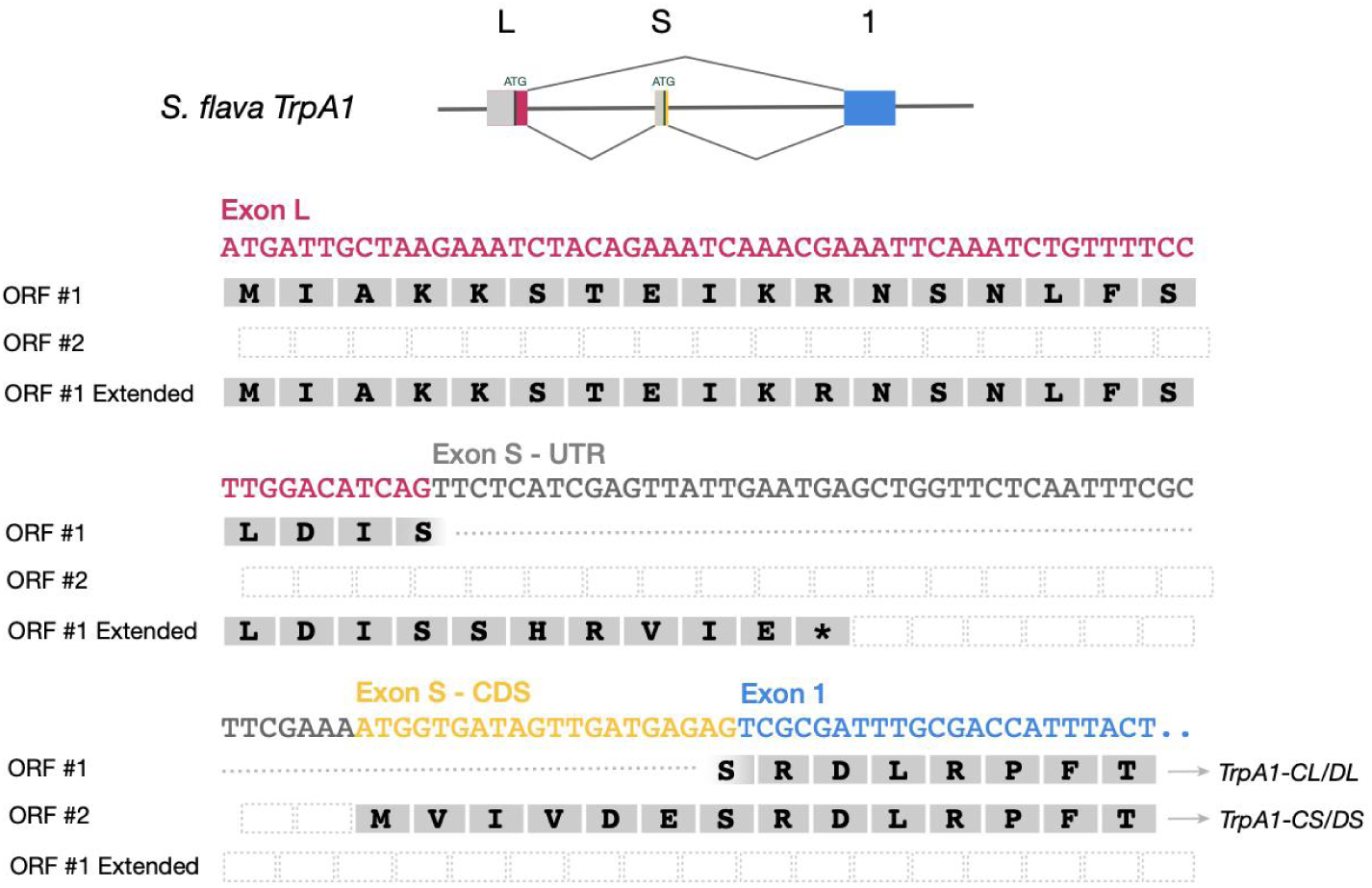
Open reading frames (ORFs) of *S. flava TrpA1* splicing isoforms. N-terminal exons and potential ORFs are shown. ORF#1 is likely a splice variant of *TrpA1-CL/DL* splicing isoforms where Exon L directly connects to Exon 1. When Exon S is incorporated into a spliced mRNA, however, ORF#1 would soon encounter a premature stop codon. The longest products would be translated from ORF#2 for *TrpA1-CS/DS* isoforms, which would result in shorter products despite the longer mRNAs. The entire Exon L may be a 5’UTR for the translation of *TrpA1-CS/DS* isoforms.

**Supplementary Figure S6.**
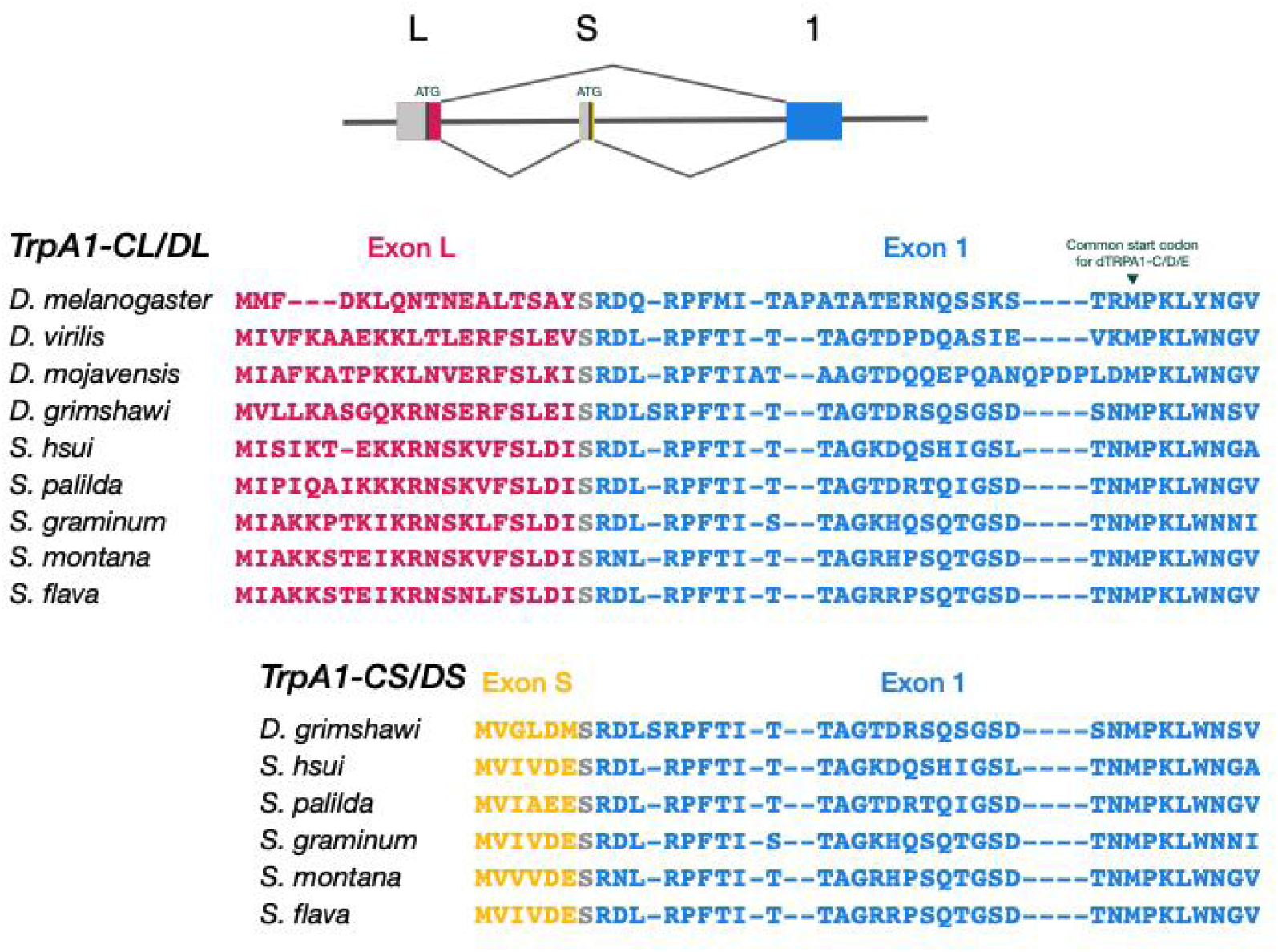
N-terminal protein alignments of *TrpA1* splicing isoforms across drosophilids. Note that Exon L and Exon 1 of *D. melanogaster TrpA1* together form a single exon (see Figure S4), the majority of which is usually treated as 5’UTR.

**Supplementary Figure S7.**
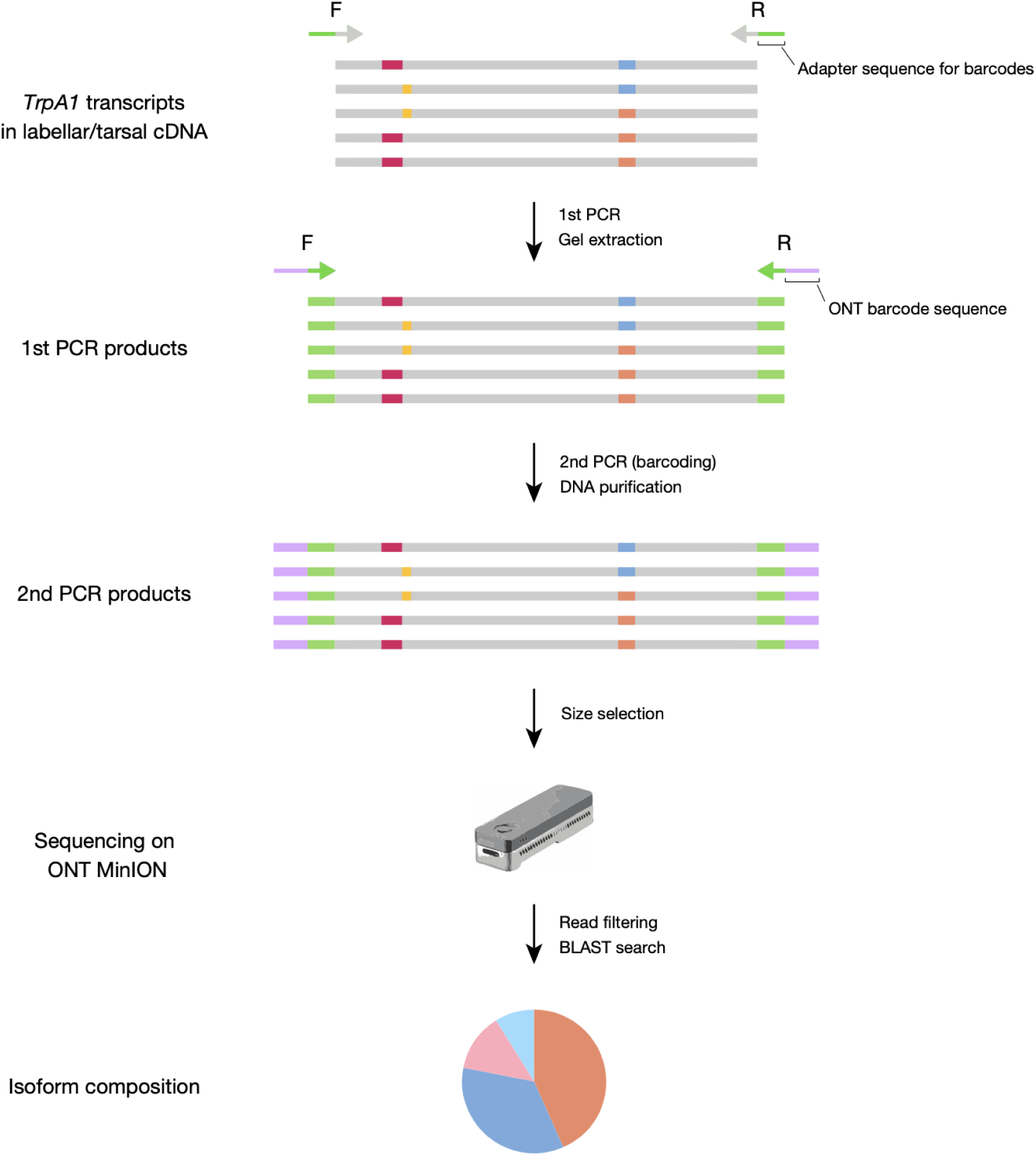
A schematic diagram of *TrpA1* amplicon sequencing. Colored regions indicate alternatively spliced exons. An original image for ONT MinION sequencer was obtained from: https://store.nanoporetech.com/us/minion-mk1b-basic-starter-pack.html.

**Supplementary Figure S8.**
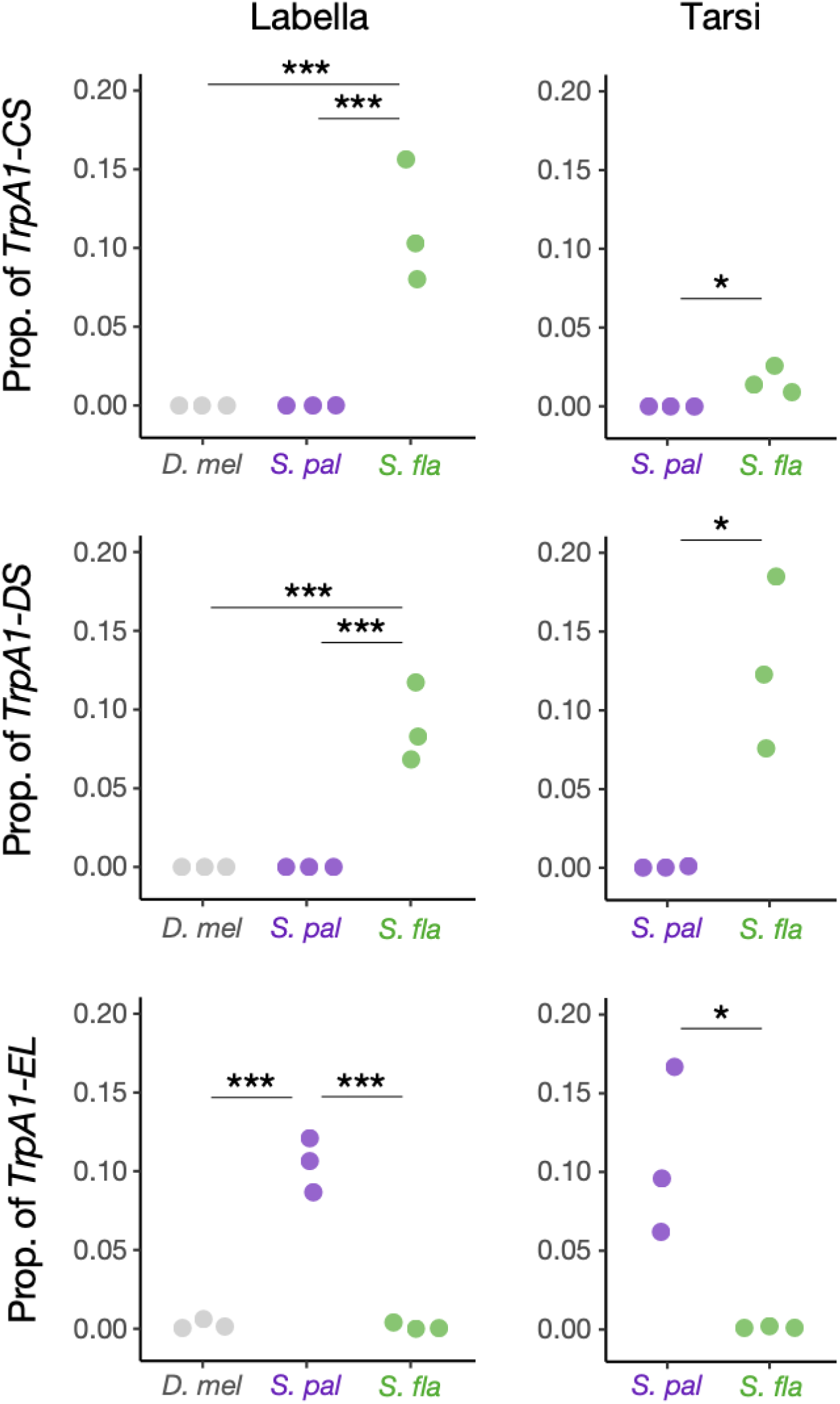
Relative abundances of *TrpA1-CL/DL/EL* splicing isoforms. Variation among species (*D. melanogaster* or *D. mel*, *S. pallida* or *S. pal*, and *S. flava* or *S. fla*) was modeled by GLM with Gaussian distribution. Labellar comparisons were further tested by Tukey’s post-hoc test with Benjamini-Hochberg correction. *: *p* < 0.05, **: *p* < 0.01, ***: *p* < 0.001.

### Herbivore-specific substitutions may modify structure of TRPA1

We next determined the extent of TRPA1 amino acid divergence between species to acertain whether it could be useful to compare physiological and *in vivo* studies downstream. Unlike the TRPA1 in the pain-insensitive African mole-rat, electrophile-detecting cysteine residues required for channel activation (Hinman et al. 2006; Kang et al. 2010) were conserved in all *Drosophila* and *Scaptomyza* TRPA1s investigated (Figure S9). We therefore ruled out the loss of cysteines as a hypothesis for reduced feeding aversion to electrophiles.

We then characterized the substitutions among species that have the potential to influence the overall structures and physiology of TRPA1 channels. Multiple sequence alignment from six *Scaptomyza* and *Drosophila* species revealed high conservation of TRPA1 proteins and identified 13 residues (out of >1,200 total AAs) that were uniquely substituted in the herbivorous or mustard-feeding lineages (i.e., *S. flava* and *S. montana*), while conserved in other fly species (Figure S10 and S11). These substitutions included four residues in ankyrin repeat domains (ARDs), four in transmembrane (TM) domains, and three in C-terminal domains (Figure S10). Among these, only two substitutions found in ARDs, V244A and N501S, and one located between the first and second TM domains, S864N, were predicted to affect local structures of the channel based of the prediction by AlphaFold Server (Abramson et al. 2024). For example, A244 in *S. flava* TRPA1 allowed the formation of a hydrogen bond between R248 and E280, while this interaction was not formed with ancestral V244 (Figure S10). Since both alanine and valine have non-polar side chains that do not form hydrogen bonds themselves, this change may be attributed to the smaller side chain of alanine. S501 was predicted to form hydrogen bonds with three adjacent residues (E497, Q498, C502), whereas N501 did not interact with any of these (Figure S10). ARDs were previously implicated with interspecific variation in thermal and chemical response of the TRPA1 (Cordero-Morales et al. 2011). These structural predictions suggest that observed herbivore-specific and mustard-feeding specific substitutions could potentially change the secondary structures of TRPA1 and influence its physiology. These findings suggest physiological comparisons of TRPA1 function would be fruitful. We also observed that evolutionary changes at the protein level were perhaps more subtle than those we found in gene expression profiles, highlighting the need for functional studies across both aspects of TRPA1 channel function.

**Supplementary Figure S9.**
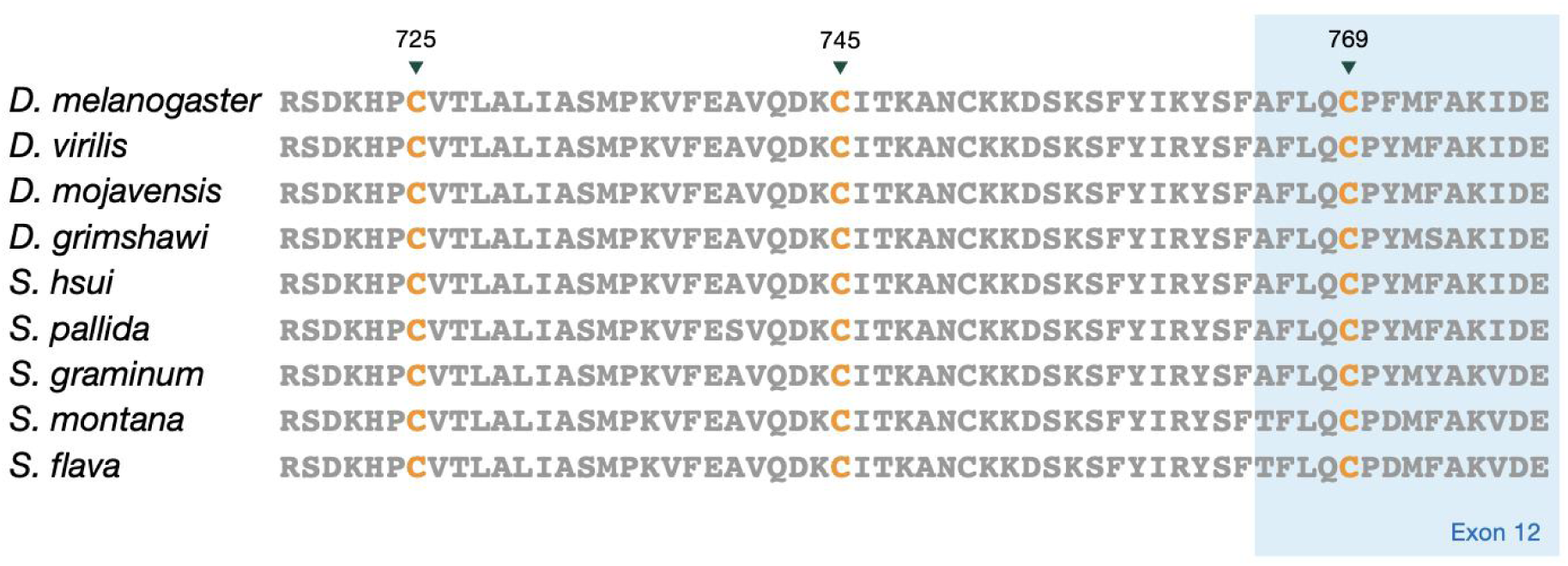
Key cysteine residues of TRPA1 are conserved across drosophilids. ITCs and many other TRPA1-activating chemicals are known to form covalent bonds with a few intracellular cysteine residues to trigger gating of the channel. Three important cysteines implicated in human TRPA1 (Hinman et al. 2006) are highlighted. Positions of amino acid residues are based on the order of *S. flava* TRPA1. Note that the cysteine at 769th residue falls on Exon 12, which is an alternatively spliced exon and only present in homologues of TRPA1-A/D.

**Supplementary Figure S10.**
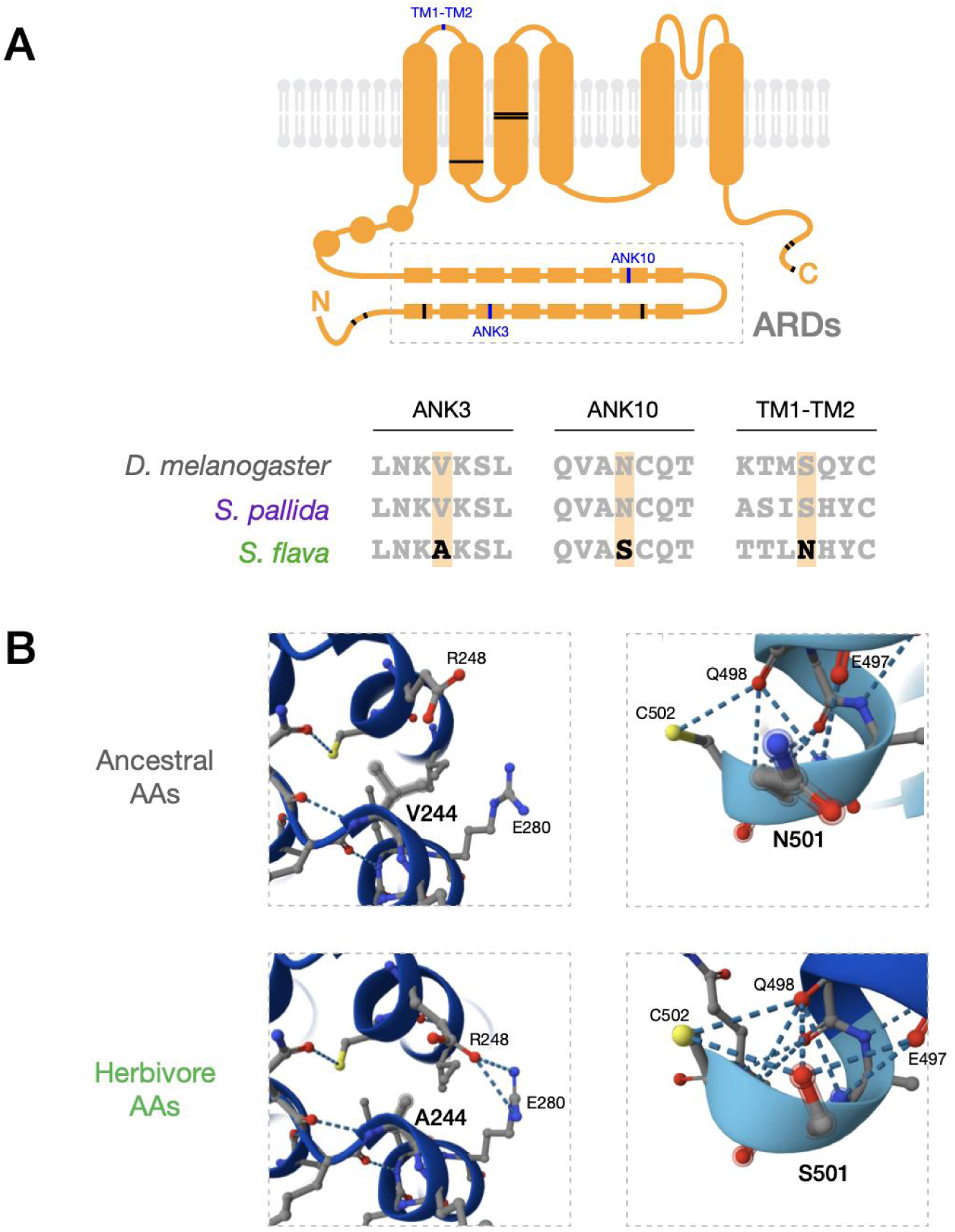
Herbivore-specific amino acid substitutions potentially modify the secondary structure of TRPA1. (A) Bars in the TRPA1 protein cartoon highlights the positions of herbivore-specific amino acid substitutions. Blue bars indicate substitutions that could potentially influence local structures of the TRPA1 channel. (B) AlphaFold-led prediction of the secondary structure and hydrogen bonds at 244th (Left, 3rd ANK domain) and 501st (Right, 10th ANK domain) residues of TRPA1 ankyrin repeat domain. Dashed lines indicate predicted hydrogen bonds between adjacent residues. The top row shows predictions with ancestral amino acids (V244, N501), while the bottom row shows those with derived herbivore-specific amino acids (A244, S501).

**Supplementary Figure S11.**
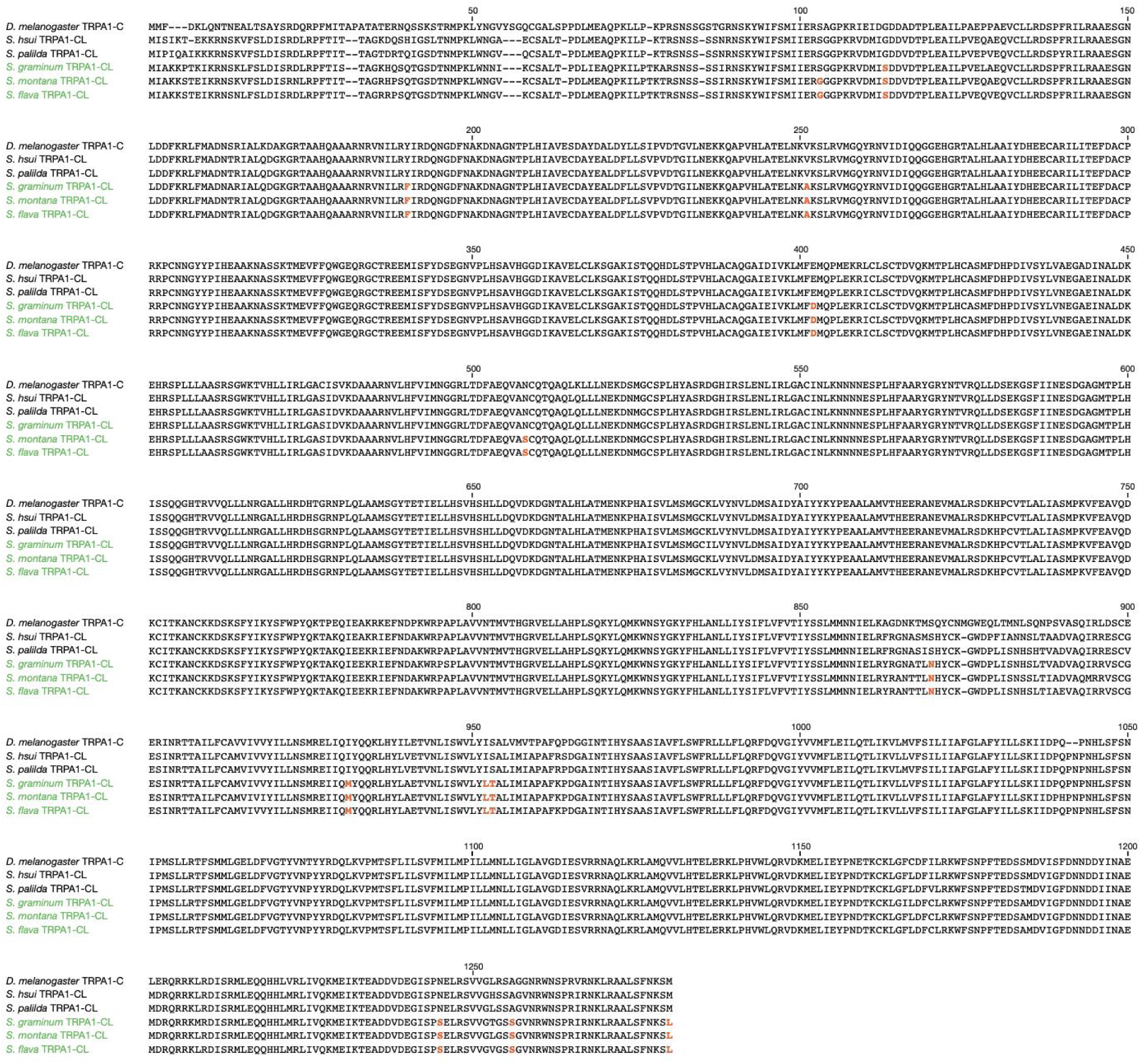
A multiple sequence alignment of TRPA1 from six *Drosophila* and *Scaptomyza* species. Green color indicates herbivorous species. Amino acid substitutions specific to herbivores or the mustard-feeding lineage (i.e., *S. montana* and *S. flava*) that are otherwise fixed are indicated in orange.

### TRPA1 channels show species-specific responses to AITC *in vitro*

We characterized *in vitro* differences in functionalities of TRPA1 channels (including the canonical splicing isoforms within and among species) detected from the three drosophilids using two-electrode voltage clamp (TEVC) recordings of *Xenopus laevis* oocytes. As reported previously (Kang et al. 2010), we observed that drosophilid TRPA1 channels stimulated by AITC did not immediately return to the baseline current (Figures 4A-4G), likely due to their unique mode of channel activation (Hinman et al. 2006; Macpherson et al. 2007). This property prevented us from applying a standard current normalization method in which a single cell is consecutively stimulated by multiple dosages such that raw current sizes are corrected by a maximum current amplitude. Instead, we used a strategy in which TRPA1-expressing oocytes were initially stimulated by heat (>40°C) and subsequently by a single application of AITC, such that an AITC-evoked peak current amplitude (*I_AITC_*) was normalized by a heat-evoked current (*I_Heat_*) at each recording (Figure 4A). This approach assumed that the sizes of heat-evoked currents reflected the degree of TRPA1 expression in oocytes, which was supported by the positive correlations between the two raw current sizes (Figure S12), and succeeded in reducing variances in the raw current data. We repeated measurements across concentration gradients of AITC and eventually obtained a dose-response relationship based on normalized current values (*I_AITC_/I_Heat_*) for each isoform to analyze the physiological property of a channel (Figure 4A). Here, our main interests were the maximum normalized current, representing the strength of channel activation, and the half-maximal effective concentration (EC_50_), representing the sensitivity of the channel to a chemical stimulus.

**Figure 4.**
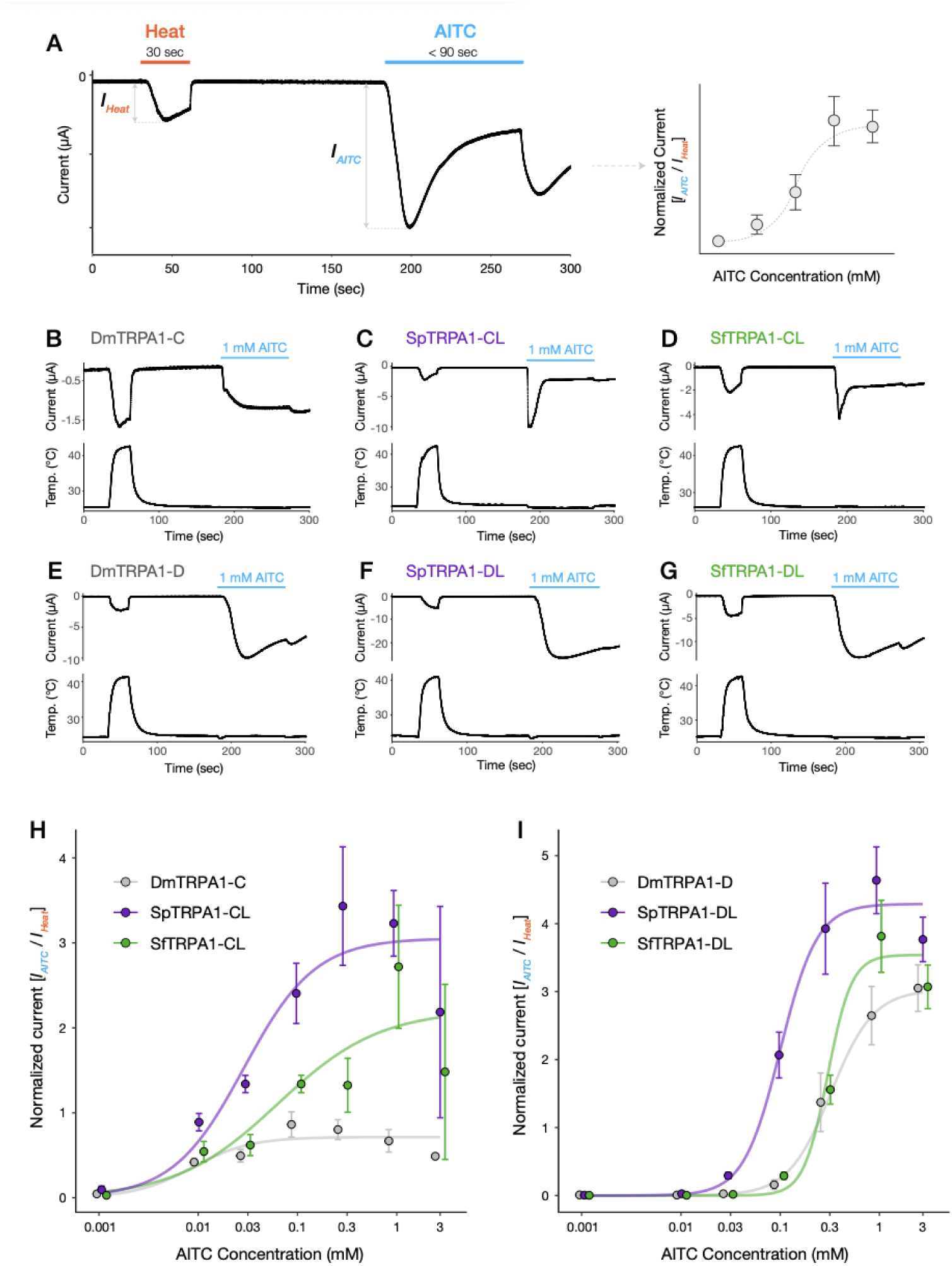
AITC-evoked responses of the TRPA1 channel vary among species *in vitro*. (A) Schematics of electrophysiological recordings from *Xenopus* oocytes and subsequent analysis. A single oocyte was stimulated first by heat (>40 °C, 30 seconds) and later by AITC (up to 90 seconds). The peak amplitude of a raw AITC-evoked current (*IAITC*) was normalized by a heat-evoked current (*IHeat*), such that a single normalized current (*IAITC*/*IHeat*) was obtained for each recording. Data collection was repeated along a dosage series of AITC to visualize the channel’s response as a dose-response curve. (B-G) Example current traces of six TRPA1 splicing isoforms analyzed against heat and AITC (1 mM) stimulations. (H) Dose-response curves of three TRPA1-CL splicing isoforms (i.e., DmTRPA1-C, SpTRPA1-CL, and SfTRPA1-CL). Circles represent mean normalized current values at each concentration of AITC (n = 4-10 per group). (I) Dose-response curves of three TRPA1-DL splicing isoforms (i.e., DmTRPA1-D, SpTRPA1-DL, and SfTRPA1-DL). Circles represent mean normalized current values at each concentration of AITC (n = 3-9 per group).

Under this framework, we conducted cross-species comparisons of AITC dose-responses for TRPA1-CL and TRPA1-DL, the two “canonical” TRPA1 splicing isoforms of *Scaptomyza* (Figure 3B and Figure S6; again, we considered *D. melanogaster* TRPA1-C/D to be homologous to *Scaptomyza* TRPA1-CL/DL because of their sequence similarity). We first found that basic properties of each TRPA1 splicing isoform type were highly conserved across species. TRPA1-CL was characterized by activation at lower AITC concentrations and rapid responses at higher concentrations (Figures 4B-4D; Figure S13). By contrast, TRPA1-DL was largely insensitive to AITC at lower doses, while its activation was greater at higher doses with a slower peaking time than TRPA1-CL (Figures 4E-4G; Figure S13). The cross-isoform differences in chemical sensitivity were also consistent with those previously reported in *D. melanogaster* and *Anopheles gambiae* TRPA1s (Du et al. 2024), suggesting that the functional differentiation among TRPA1 splicing isoforms may be widely conserved throughout the Diptera. Moreover, all canonical splicing isoforms responded to heat stimulation in oocytes, consistent with earlier recordings from *D. melanogaster* cell lines (Gu et al. 2019).

Despite some degree of functional conservation, however, we also observed substantial interspecific variation in the activity and the sensitivity of the TRPA1 channels to AITC. Among the three focal species we studied, *S. pallida* TRPA1 (SpTRPA1) generated the largest normalized currents for both splicing isoform types (Figures 4H and 4I; Table 1). Surprisingly, the weakest activity (i.e., maximum current amplitude) was found for both isoforms of *D. melanogaster* TRPA1 (DmTRPA1; Figures 4H and 4I; Table 1), despite the strong behavioral aversion to AITC in this species (Figure 1B). *S. flava* TRPA1 (SfTRPA1) splicing isoforms were as weakly activated as DmTRPA1 at lower doses, but they eventually generated large currents at higher AITC concentrations (Figures 4H and 4I; Table 1). Median effective concentrations (EC_50_) were overall lower in TRPA1-CL isoforms than in TRPA1-DL isoforms (Table 1), consistent with visual observations of current traces (Figure 4B-4G). In both isoforms, SfTRPA1 showed larger EC_50_ values against AITC than SpTRPA1 (Table 1). DmTRPA1-C was by far the most sensitive channel among TRPA1-CL isoforms (Table 1).

**Table 1.** Summary of electrophysiological recordings from six canonical TRPA1 splicing isoforms. Maximum normalized currents and EC50 were predicted values from dose-response curve estimation by drc package (Ritz et al. 2015). Values following the (±) symbol represent standard errors for corresponding parameters.

|  | TRPA1-CL |  |  | TRPA1-DL |  |  |
| --- | --- | --- | --- | --- | --- | --- |
|  | DmTRPA1-C | SpTRPA1-CL | SfTRPA1-CL | DmTRPA1-D | SpTRPA1-DL | SfTRPA1-DL |
| Maximum normalized current [$I_{AITC}/I_{Heat}$] | 0.714±0.059 | 3.05±0.34 | 2.21±0.47 | 3.03±0.37 | 4.29±0.29 | 3.54±0.22 |
| EC<sub>50</sub> (mM) | 0.0086±0.0043 | 0.029±0.015 | 0.065±0.055 | 0.341±0.092 | 0.103±0.015 | 0.316±0.031 |
| Mean $I_{Heat}$ (μA) | 2.60 | 1.39 | 1.82 | 3.52 | 4.08 | 3.32 |
| Number of recordings | 35 | 42 | 40 | 31 | 34 | 35 |

Because our current normalization relied on the sizes of heat-evoked currents, it was possible that a channel with a weaker heat activation (*I_Heat_*) was represented with larger normalized currents (*I_AITC_*/*I_Heat_*). This relationship was partially observed for TRPA1-CL isoforms, where SpTRPA1-CL exhibited significantly weaker heat currents than the other two homologous channels (Figure S14; Table 1). However, these differences in heat-evoked currents did not fully explain their variation in electrophile sensitivity. For instance, the difference of the maximum normalized currents between DmTRPA1-C and SpTRPA1-CL (4.28-fold difference; Figure 4H; Table 1) was much greater than that of heat currents (1.87-fold difference; Figure S14; Table 1), which alone did not resolve the cross-species variation in normalized currents. We did not observe significant differences in heat currents among TRPA1-DL isoforms (Figure 4K). We concluded that the observed cross-species differences in the activity of TRPA1 isoforms still hold true even after taking into account their variation in heat activation.

Taken together, SfTRPA1 indeed showed lower sensitivity to AITC and a smaller current amplitude in response compared with SpTRPA1, which suggests that the physiology of the TRPA1 channel evolved concurrently with the transition to herbivory in the *S. flava* lineage and shaped the weak electrophile avoidance. However, these physiological differences in TRPA1 function between species did not entirely align with their differences in behavioral aversion towards AITC (Figure 1B), due to the robust activity of SfTRPA1 at the higher AITC concentrations and the observation that the weakest current amplitude was of DmTRPA1. This suggests that alternative mechanisms, such as differential expression of differentially ITC-sensitive splicing isoforms may underlie the differences in feeding aversion towards AITC among species.

**Supplementary Figure S12.**
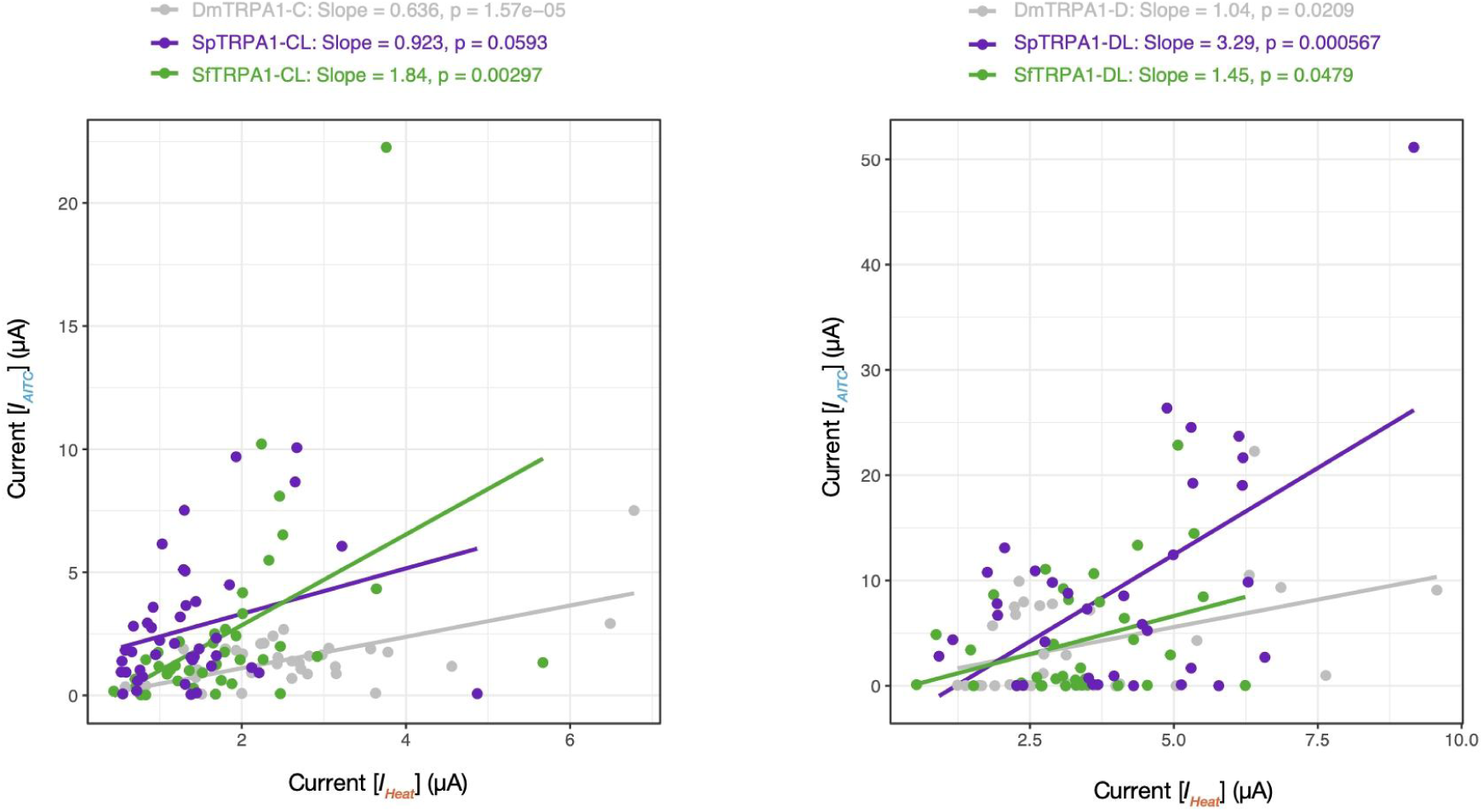
Correlations between raw heat-evoked and AITC-evoked currents. Each dot represents a single oocyte. (Left) TRPA1-CL and (Right) TRPA1-DL splicing isoforms. Data from all AITC concentrations were pooled for each splicing isoform.

**Supplementary Figure S13.**
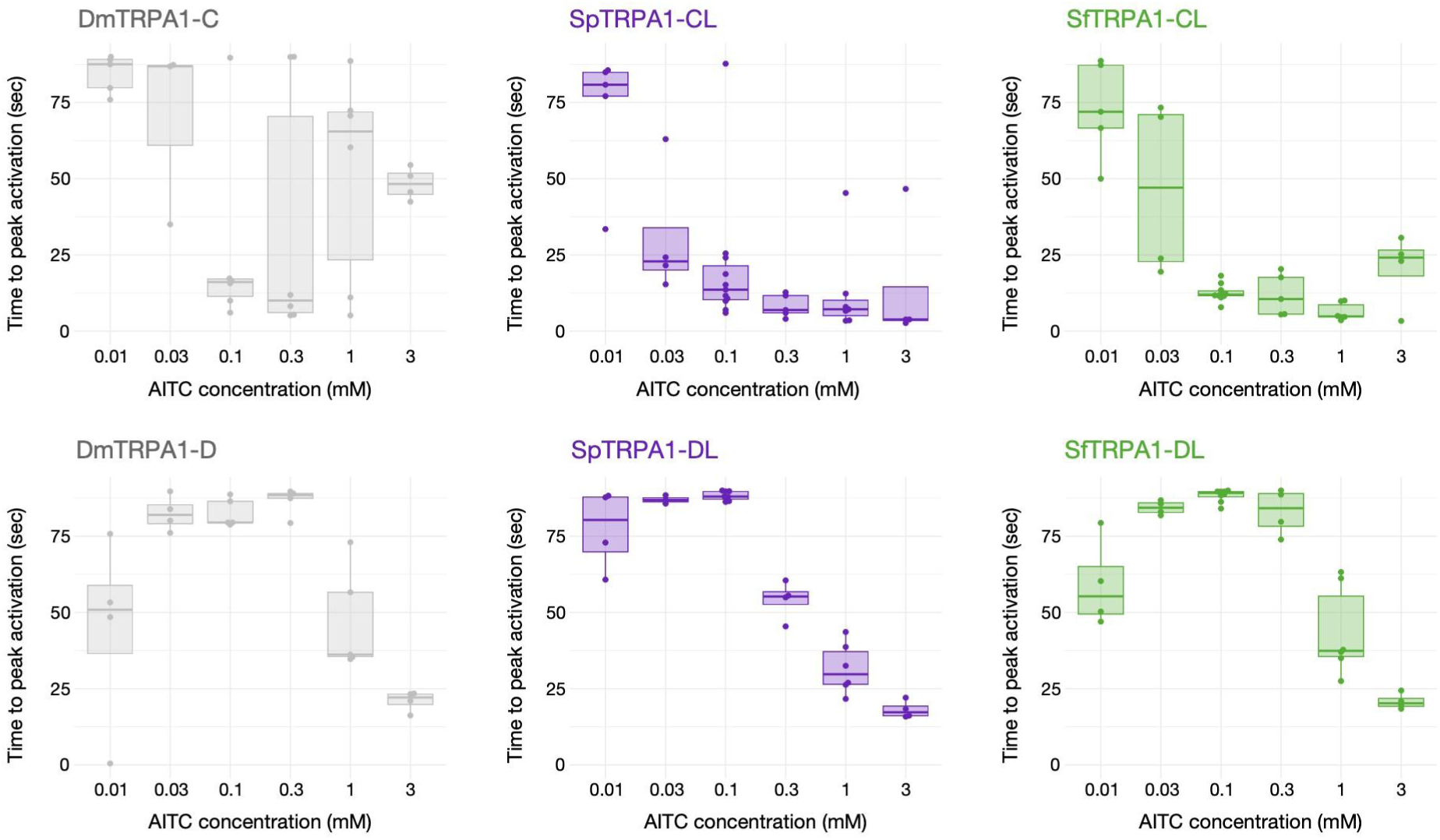
Times to reach peak activation in six canonical TRPA1 splicing isoforms. Recordings were carried out within a 90-second AITC stimulation window, across the concentration series from 0.001 mM to 3 mM AITC. Note that the smallest concentration (0.001 mM AITC) was omitted from this analysis due to the lack of observable chemical-invoked currents. Overall, TRPA1-CL(C) isoforms tended to peak faster than TRPA1-DL(D) splicing isoforms.

**Supplementary Figure S14.**
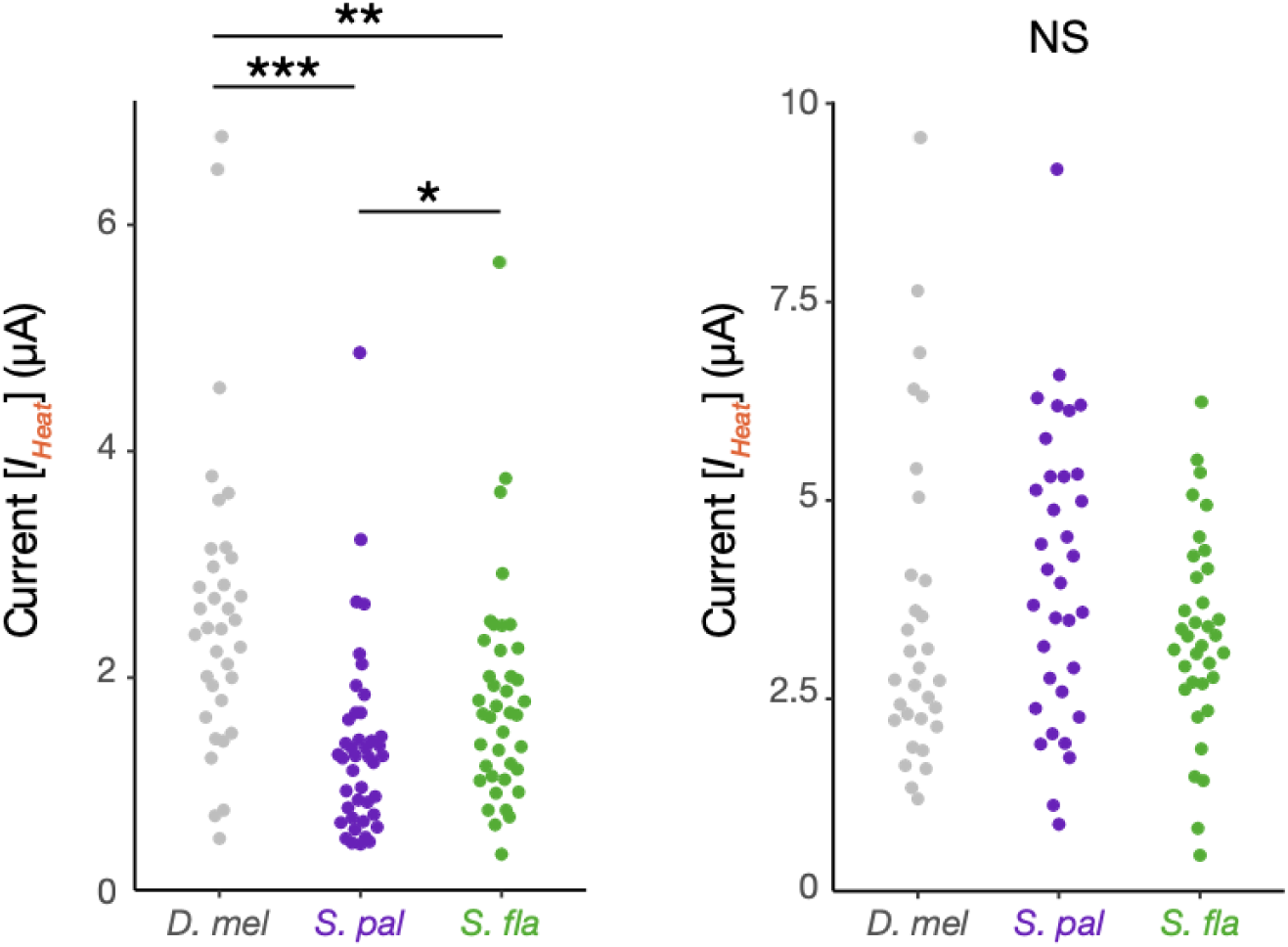
Comparisons of raw heat-evoked currents (*IHeat*) (Left) TRPA1-CL and (Right) TRPA1-DL splicing isoforms, tested by Kruskal-Wallis test and post-hoc Conover’s test with Benjamini-Hochberg correction. *: *p* < 0.05, **: *p* < 0.01, ***: *p* < 0.001.

### Ectopic expression of TRPA1-CL splicing isoforms partially recapitulates wild-type behaviors

Given the interspecific differences in the physiological responses of the TRPA1 channels to AITC, we next addressed whether and to what extent those changes could influence feeding aversion behavior and its evolution. To this end, we designed an *in vivo* genetic rescue experiment using the *GAL4/UAS* system in *D. melanogaster* to test whether *in vivo* expression of TRPA1 isoforms from *Scaptomyza* spp. and *D. melanogaster* can recapitulate the differences in gustatory behaviors between wild-type flies of these species. Using PhiC31-mediated insertion, we created three homozygous *UAS-TrpA1* lines each carrying a full-length copy of *SfTrpA1-CL*, *SpTrpA1-CL* or *DmTrpA1-C*, respectively, and drove their expression in bitter-sensing gustatory neurons under the *DmGr66a* promoter (Lee et al. 2009; Weiss et al. 2011) (Figure 5A). *TrpA1-CL* was selected as the focal splicing isoform since it is the only canonical isoform abundantly expressed in all three species (Figure 3B). All *Gr66a-GAL4* and *UAS-TrpA1* lines were initially set to *TrpA1* null (*TrpA1^1^*) mutant background (Figure S15) and crossed for experiments (Figure 5A). Test genotypes (i.e., *Gr66a>TrpA1*) and control flies (i.e., *UAS-TrpA1/+*) were then phenotyped using the PER assay to evaluate the contributions of ectopic *TrpA1* expression on their electrophile sensitivities. Flies were challenged with a non-volatile electrophile NMM (N-methylmaleimide) as a deterrent instead of an ITC, because NMM invokes a similar gustatory response as AITC but is far less volatile and would not simultaneously risk activating olfactory pathways, which are responsive to volatile ITCs (Matsunaga et al. 2025).

**Figure 5.**
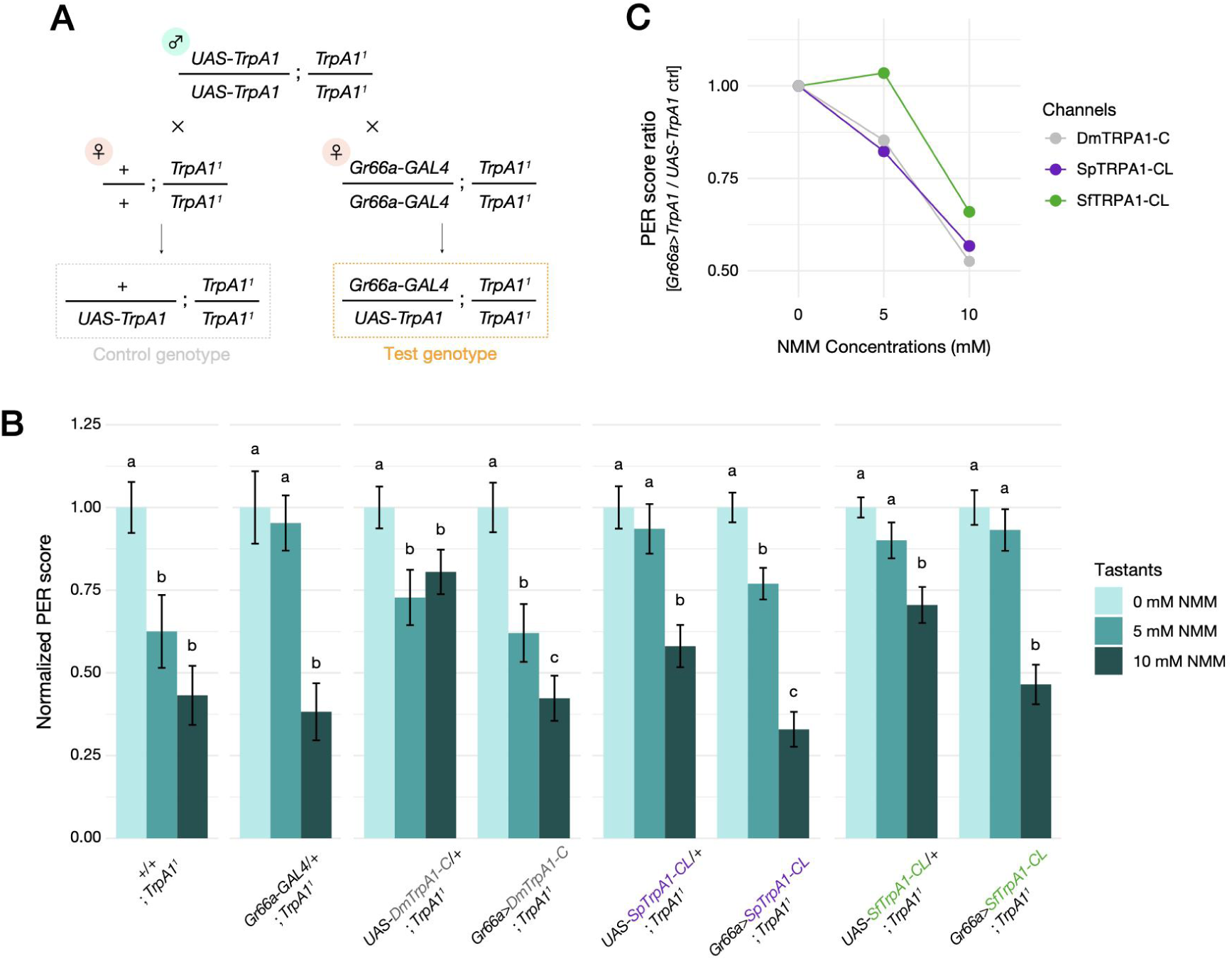
Ectopic expression of *Drosophila/Scaptomyza* TRPA1s partially rescues wild-type behavior. (A) Crossing schemes of *Gr66a>TrpA1* (test group) and *UAS-TrpA1* (control group) flies. Note that all flies were homozygotes of *TrpA1^1^* allele that lacked endogenous *dTrpA1* expression. (B) Results of PER assay in eight genotypes across the concentration series of NMM mixed in 100 mM sucrose solution. PER score of a single female fly represents the feeding response to a given tastant over five trials. Bar plots indicate individual PER scores for each treatment (n = 15-42) normalized to the average score at 0 mM NMM within each genotype (see Method for details of data conversion). Letters indicate significant differences between NMM concentrations within each genotype, tested by pairwise Wilcoxon rank-sum test with Benjamini-Hochberg correction (*p* < 0.05). (C) Ratios of average PER scores between *Gr66a>TrpA1* line and *UAS-TrpA1/+* line for each TRPA1 channel expressed, plotted as line graphs over the three concentrations of NMM. Error bars are not applicable because the data is n = 1 for each genotype and NMM concentration.

Although all fly mutants had a *TrpA1*-null background, 10 mM NMM was consistently aversive to all control genotypes (Figure 5B). 5 mM NMM showed mixed results: flies of the *TrpA1^1^* and *UAS-DmTrpA1-C* lines avoided NMM, while flies of other control genotypes did not show clear signs of aversion compared with the 0 mM NMM treatment (Figure 5B). These patterns suggest that the aversion towards NMM was retained in our transgenic lines via unknown TRPA1-independent mechanisms that were also likely influenced by different genetic backgrounds. Nevertheless, we still found that ectopic expression of SpTRPA1-CL clearly and consistently increased the degree of aversion at 5 mM NMM compared with its control strain, while SfTRPA1-CL did not significantly confer a feeding aversion behavior (Figure 5B), a pattern consistent with the physiological responses we observed in oocyte electrophysiology (Figure 4H).

To further isolate the contribution of ectopic TRPA1 expression in *D. melanogaster*, we took the ratios of average PER scores between test (*Gr66a>TrpA1*) and control (*UAS-TrpA1/+*) genotypes for each TRPA1 channel and plotted these as dose-responses (Figure 5C). SfTRPA1-CL conferred weaker electrophile aversion at 5 mM and 10 mM NMM compared with SpTPRA1-CL or DmTRPA1-C (Figure 5C), consistent with a model of phenotypic evolution wherein TRPA1 channel property underlies the attenuated electrophile aversion of *S. flava*. However, the differences in feeding aversion were much more subtle between *D. melanogaster* flies ectopically expressing SfTRPA1 or SpTRPA1 (Table 2; Figure 5B) than between wild-type *S. flava* and *S. pallida* flies (Table 2; Figure 1B). This mismatch indicates that TRPA1 physiology alone cannot fully recapitulate interspecific feeding preferences, potentially due to other mechanisms including the involvement of other nociceptive molecular sensors (e.g., Tracey et al. 2003; Al-Anzi et al. 2006) or regulation of alternative splicing. Overall, our genetic rescue experiments revealed the potential importance of TRPA1 channel physiology for the evolution of gustatory behaviors, but also highlighted the presence of additional molecular mechanisms that might contribute to the near-absence of electrophile aversion in *S. flava*.

**Table 2.**
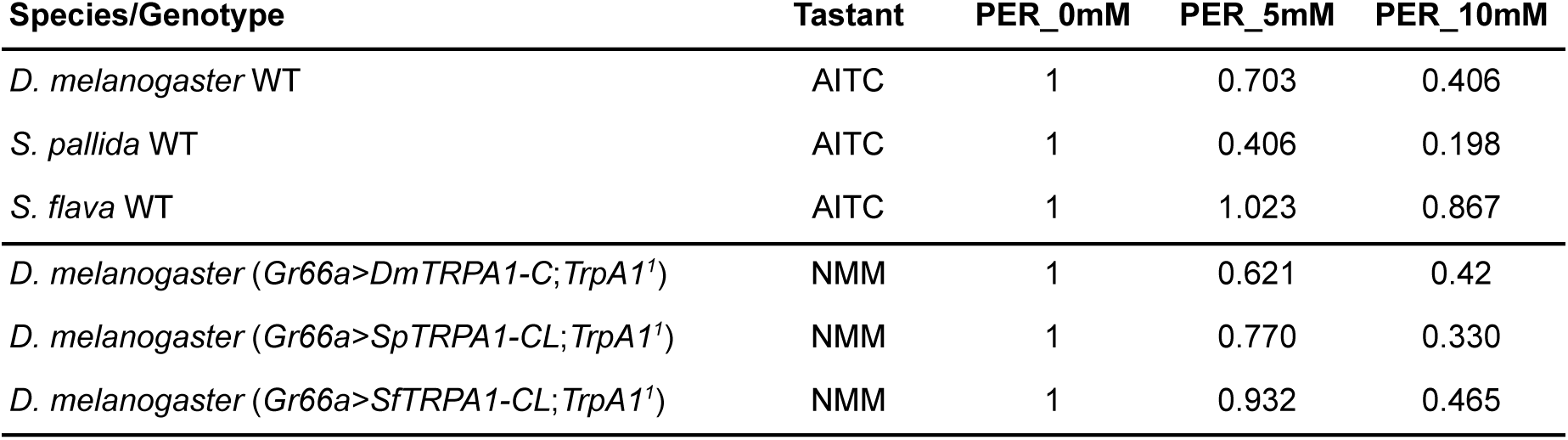
Summary of wild-type and transgenic PER assays. Average PER scores normalized to 0 mM AITC or NMM were retrieved from Figure 1B (wild-type) and three rescue genotypes in Figure 5B (transgenic)

**Supplementary Figure S15.**
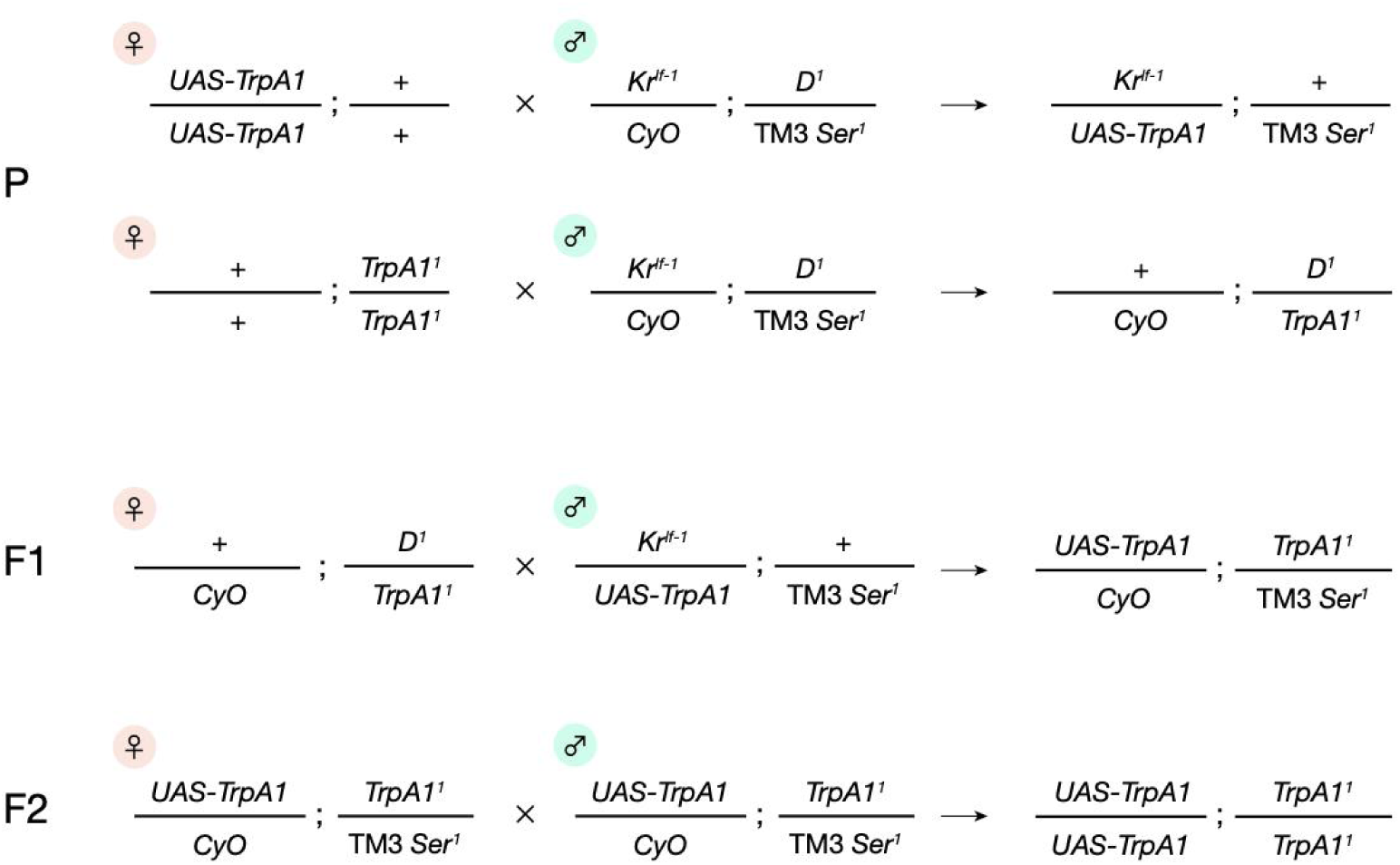
Crossing schemes to obtain double homozygotes of *UAS-TrpA1* and *TrpA1^1^* alleles. A double balancer line *Kr^lf-1^/CyO;D^1^*/TM3 *Ser^1^*) was utilized to merge two alleles. Only second and third chromosomes are shown. *Gr66a-GAL4*;*TrpA1^1^*homozygotes were also obtained in the same procedure.

### Co-expression of canonical TRPA1 splicing isoforms modulates AITC-evoked currents *in vitro*

To search for alternative mechanisms behind sensory adaptation in *S. flava* beyond gene expression and TRPA1 sequence differences, we turned our attention to the regulation of alternative splicing of the *TrpA1* gene, inspired by the example from the vampire bat in which a truncated TRPV1 splice variant underlies prey finding behavior (Gracheva et al. 2011). Amplicon sequencing highlighted the difference in expression proportions of the two canonical TRPA1 isoforms, TRPA1-CL and TRPA1-DL, between *S. flava* and *S. pallida* (Figure 3B and 3C). Since these two isoforms exhibited differential sensitivity towards AITC (Table 1; Figure 4H and 4I), we hypothesized that variation in co-expression of canonical TRPA1 isoforms influences the chemical sensitivity of gustatory sensory neurons. To this end, we explored the functional consequences of isoform co-expression, again using *Xenopus* oocyte electrophysiology. By taking advantage of direct RNA injection into oocytes, we were able to precisely manipulate the compositions of TRPA1 isoforms expressed in a cell by pre-mixing multiple mRNAs at a particular ratio. For initial experiments, we prepared canonical *TrpA1* mRNA mixtures in integer ratios that best represented our observations from amplicon sequencing (Figure 3B; i.e., *S. flava* labella CL:DL=7:6, *S. pallida* labella CL:DL=5:2, *S. flava* tarsi CL:DL=1:7, *S. pallida* tarsi CL:DL=6:7).

We found that cells responded more actively at lower AITC concentrations as the proportion of TRPA1-CL, an isoform with a higher chemical sensitivity, increased in both species (Figure 6A and 6B). By contrast, enrichment of the TRPA1-DL isoform decreased the current amplitudes at low AITC concentrations (Figure 6A and 6B). These patterns were consistent with our working hypothesis and showed that co-expressed TRPA1 isoforms additively modulated the current response of cells and potentially of gustatory neurons.

**Figure 6.**
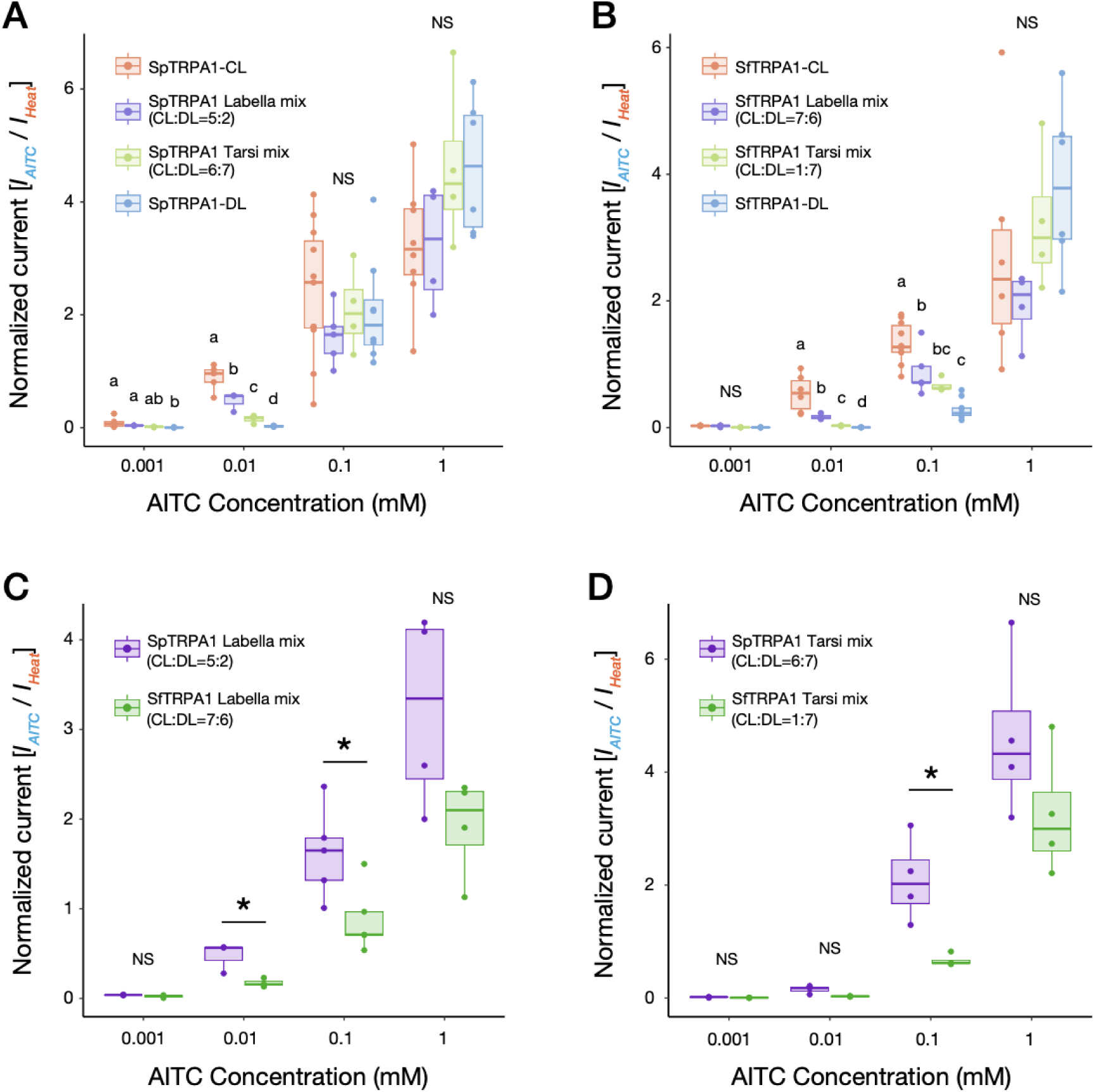
Co-expression of canonical TRPA1 isoforms *in vitro* additively modulates electrophile-evoked responses. (A, B) Heat-normalized AITC-evoked currents of (A) *S. pallida* TRPA1 mixtures (n = 3-11) and (B) *S. flava* TRPA1 mixtures (n = 3-10) at the four concentrations of AITC in recordings of *Xenopus laevis* oocytes. Differences between isoforms were tested by one-way ANOVA followed by post hoc Tukey’s post hoc test (0.1 and 1 mM AITC) or Kruskal-Wallis test followed by post hoc Conover’s test (0.001 and 0.01 mM AITC) with Benjamini-Hochberg correction. Letters indicate the significant differences of the mean between groups at each concentration (*p* < 0.05). (C, D) Heat-normalized AITC-evoked currents of RNA mixtures mimicking (C) labellar *TrpA1* expression (n = 3-5 per group) and (D) tarsal *TrpA1* expression (n = 3-4 per group), using the same datasets from (A) and (B). Differences between species at each concentration were tested by Student’s t-test or Welch’s t-test. *: *p* < 0.05.

Using the same dataset, we further compared the normalized responses of labella- and tarsi-mimicking mixtures between *S. flava* and *S. pallida*. We found that *S. flava* TRPA1 mixtures yielded significantly weaker responses than *S. pallida* at 0.1 mM AITC in both mixtures (Figures 6C and 6D), consistent with the weaker current amplitudes of SfTRPA1 than of SpTRPA1 (Figures 4H and 4I). Intriguingly, the degree of interspecific difference varied at 0.01 mM AITC: responses of oocytes were significantly greater for labellar mixtures of *S. pallida* than of *S. flava* (Figure 6C), while those for tarsal mixtures did not differ between species (Figure 6D). The relative inactivity of *S. flava* labellar mixture at 0.01 mM was likely achieved through the higher proportion of TRPA1-DL in this mixture compared with its *S. pallida* counterpart (Figure 3B and 3C). This pattern indicates that species-specific or tissue-specific regulation of alternative splicing could indeed drive sensory differentiation among drosophilids with divergent dietary needs. Overall, our results with mRNA mixtures that phenocopied the native isoform ratios in labella and tarsi of flies from different species provide a model for howthe physiological response of the sensory neurons can be modulated through differential regulation of TRPA1 isoforms at least *in vitro*.

**Supplementary Figure S16.**
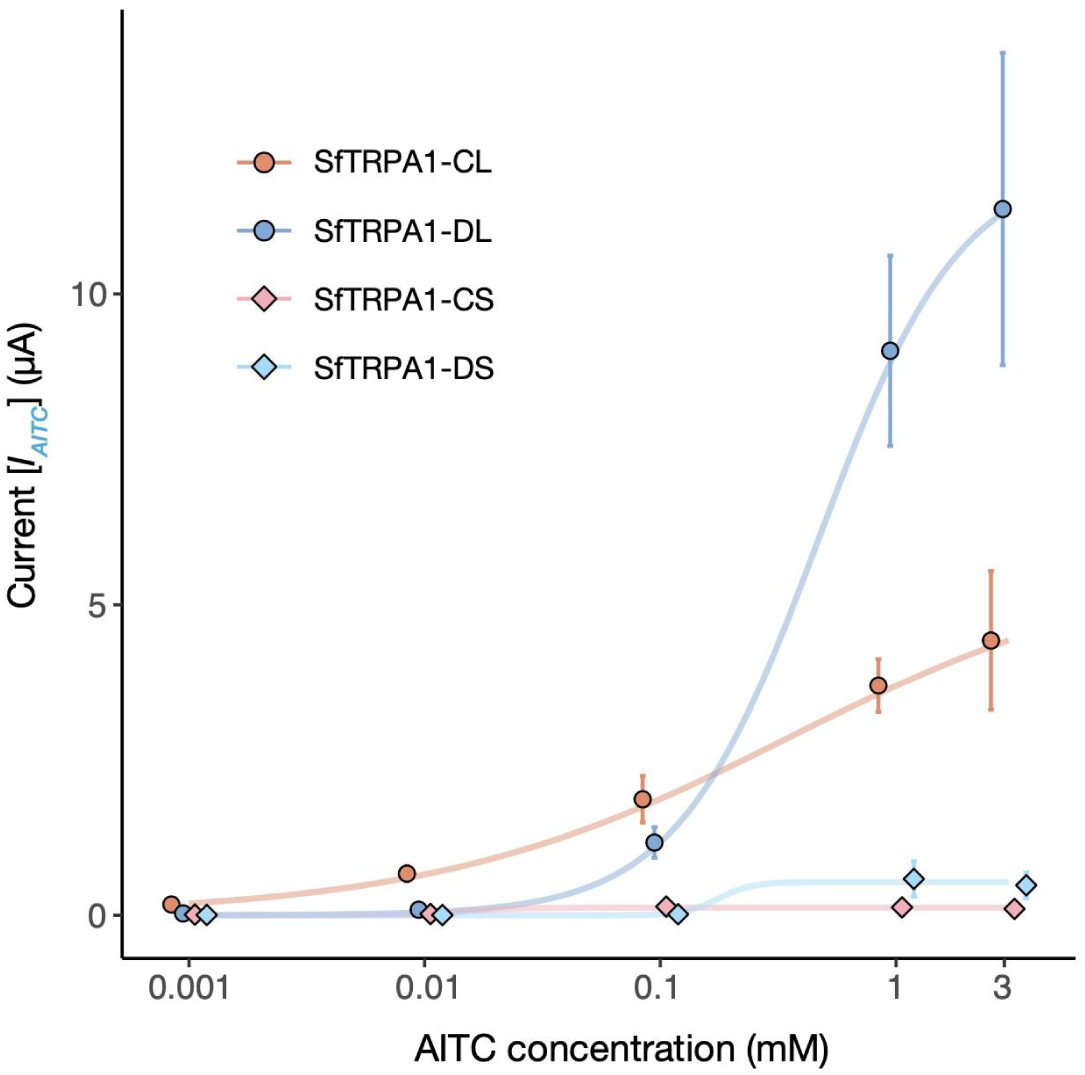
Dose-response curves of the four SfTRPA1 isoforms. Note that the prediction of dose-response was performed on raw current values (*IAITC*), instead of normalized values (*IAITC*/*IHeat*), because weak heat activation of the inactive isoforms prevented proper normalization. n = 3-12 per group.

### Co-expression of inactive TRPA1 isoforms attenuates AITC-evoked currents

Our amplicon sequencing also revealed two underrepresented TRPA1 isoforms from *S. flava*, SfTRPA1-CS and SfTRPA1-DS (Figures 3B and 7A). We next investigated the characteristics of these minor TRPA1 channel variants and found that their responses to AITC or heat stimulations were substantially weaker than those of canonical isoforms, even at the highest AITC concentrations (Figure 7C; Figure S16). Likewise, a minor isoform in *S. pallida*, SpTRPA1-EL, barely responded to both stimuli (Figure 7B), consistent with its homology to the DmTRPA1-E isoform with a similar non-responsive phenotype (Gu et al. 2019).

**Figure 7.**
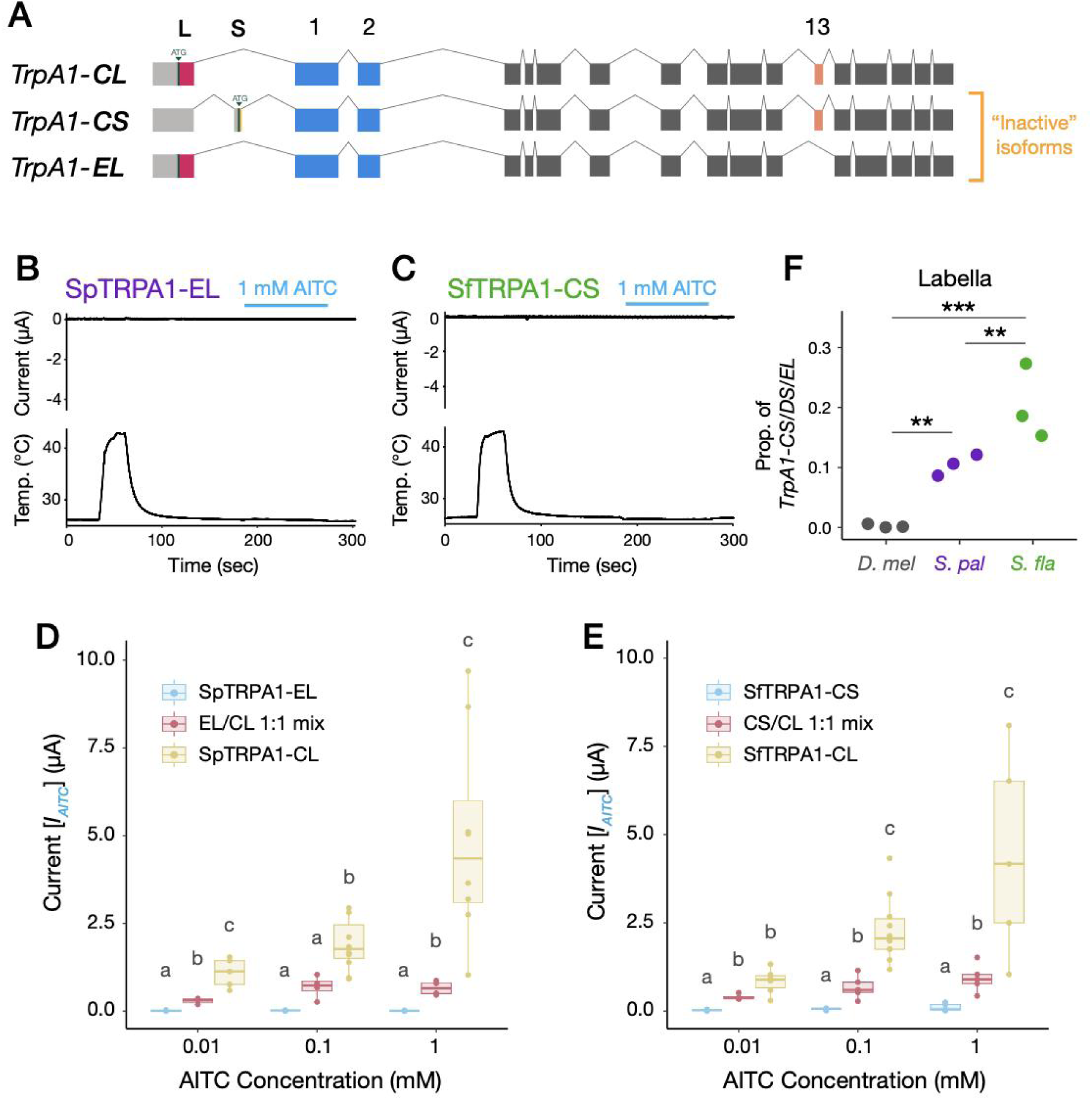
Co-expression of “inactive” TRPA1 isoforms reduces electrophile-evoked responses *in vitro*. (A) Exonic structures of *Scaptomyza TrpA1-CL* and two “inactive” isoforms, *TrpA1-CS* and *TrpA1-EL*. Green small arrowheads indicate putative translation start sites, whose upstream are putative 5’UTRs (pale colored). (B, C) Example current traces (from *Xenopus* oocytes) of (B) SpTRPA1-EL and (C) SfTRPA1-CS stimulated by heat and 1 mM AITC. (D, E) Comparisons of raw AITC-evoked current amplitudes between TRPA1-CL, an inactive TRPA1 isoform (either TRPA1-EL or -CS), and a 1:1 mixture of the two isoforms from (D) *S. pallida* (n = 3-11) and (E) *S. flava* (n = 4-10). Letters indicate the statistical differences in the mean current amplitudes at each AITC concentration using Kruskal-Wallis test followed by post-hoc Conover’s test with Benjamini-Hochberg correction (*p* < 0.05). (F) Relative abundances of all inactive isoforms together (i.e., *TrpA1-CS*+*TrpA1-DS*+*TrpA1-EL*) in labellar cDNA, based on the data from Fig. 3B (n = 3 per species). Variation in splicing isoform compositions among species was modeled by GLM with Gaussian distribution, and further tested by Tukey’s post-hoc test with Benjamini-Hochberg correction. **: *p* < 0.01, ***: *p* < 0.001.

We further investigated how these “inactive” TRPA1 isoforms may affect the physiology of sensory neurons based on oocyte electrophysiology. To evaluate the contributions of the inactive TRPA1s, we created 1:1 mRNA mixtures of both an inactive and a canonical TRPA1 isoform (i.e., TRPA1-CL) for *S. flava* and *S. pallida*, and compared their responses to single-isoform recordings. The activity of the 1:1 mixtures fell in between those from the inactive and the canonical isoforms in both species (Figures 7D and 7E), indicating that the presence of inactive TRPA1 isoforms may simply lower the neuronal response to AITC in an additive manner. Since the proportion of inactive splicing isoforms in labella was greater in *S. flava* than in *S. pallida* or in *D. melanogaster* (Figure 7F), it is possible that these inactive TRPA1 channels play an additional role in causing the evolutionary divergence of gustatory behavior among these species.

Taken together with the data from canonical isoform mixtures, these *in vitro* experimental manipulations revealed that the enriched expression of a particular TRPA1 splicing isoform could shift the cellular response towards the phenotype conferred by this isoform, whether that is towards higher or lower sensitivity to AITC stimulation. We hypothesize that such a mechanism may account in part for the observed differences between species (and organs) in mediating gustatory avoidance behaviors of flies towards AITC.

## DISCUSSION

Niche shifts in animals are often coupled with the evolution of behavioral traits (Duckworth 2009), but the genetic basis and molecular mechanisms underlying such events remain elusive. Chemosensory traits are good candidates for exploring this problem because of their relative simplicity and the modularity of their functional units. Here, we investigated the evolutionary, genetic, and molecular basis of differences in gustatory behavior between a relatively recently-derived specialist herbivore, *Scaptomyza flava*, and its microbe-feeding relatives, *S. pallida* and *Drosophila melanogaster*. Importantly, the *Scaptomyza* genus is nested phylogenetically within the paraphyletic subgenus *Drosophila*, which includes the Hawaiian *Drosophila* as well as *D. mojavensis* and *D. virilis*. Thus, species of *Scaptomyza* are of comparable phylogenetic distance from the genetic model species *D. melanogaster* as these congeners. By focusing on the TRPA1 channel, a primary transducer of electrophilic compounds like mustard-derived ITCs, which are so important in the niche shift of *S. flava* to GLS-bearing plants, we aimed to decipher the genetic and molecular mechanisms associated with behavioral evolution in this mustard-plant specialist. Specifically, we hypothesized that *S. flava* has evolved a weaker gustatory aversion against ITCs and that some combination of changes in the expression, protein sequence, and regulation of alternatively spliced transcripts (encoding the splicing isoforms) of the *TrpA1* gene contributed to this reduction in avoidance to ITCs.

Our behavioral assay revealed significantly attenuated feeding aversion against allyl isothiocyanate (AITC) in *S. flava*, likely mediated by changes in the gustatory circuit (Figure 1B). However, the *S. flava* genome maintains an intact *TrpA1* gene and the *S. flava* TRPA1 (SfTRPA1) channel is functionally conserved as an electrophile-gated ion channel like its orthologues in *D. melanogaster* and *S. pallida* (Figure 4H and 4I). We previously reported the entire loss of behavioral attraction to yeast odor in *S. flava* and the associated losses of genes encoding receptors tuned to these volatiles, along with their dietary transition from microbe feeders to mustard specialists (Goldman-Huertas et al. 2015). Such extreme chemosensory loss was not the case for gustatory aversion towards ITCs, likely because fly TRPA1 also mediates detection of a wide array of nociceptive stimuli and its loss would be highly deleterious (Himmel and Cox 2020).

Another element of functional conservation of TRPA1 across species is the physiological difference between splicing isoforms. Functional diversity among *D. melanogaster* TRPA1 (DmTRPA1) splicing isoforms has been documented elsewhere (Kang et al. 2012; Zhong et al. 2012; Gu et al. 2019; Leung et al. 2020; Gu et al. 2022; Du et al. 2024). Isoform DmTRPA1-C and its homologs are consistently described as channels more sensitive to chemicals compared with isoform DmTRPA1-D and its homologs (Leung et al. 2020; Du et al. 2024) We found that such differences in dose dependency were also conserved between two “canonical” TRPA1 splicing isoforms of *S. flava*, TRPA1-CL and -DL (Figure 4H and 4I; Figure S13). Although canonical *Scaptomyza TrpA1s* recruited a new exon at the N-terminus (Exon L; Figure 3A), this alone did not seem to confer a novel response property to the channel.

Despite a high degree of amino acid sequence conservation, we also discovered some salient physiological differences between *S. flava* TRPA1 (SfTRPA1) and *S. pallida* TRPA1 (SpTPRA1) that could be associated with adaptation to mustard plants. Notably, SfTRPA1 exhibited a lower amplitude of ionic currents and a lower electrophile sensitivity in both canonical isoforms compared with SpTRPA1 (Figure 4H and 4I; Table 1). Such physiological shifts in the TRPA1 channel would decrease firing rates of peripheral gustatory neurons against ITCs and thus reduce behavioral aversion, consistent with *S. flava*’s mustard-feeding ecology. Genetic rescue experiments in *D. melanogaster* further recapitulated the behavioral variation across species, wherein SpTRPA1-CL conferred a larger degree of electrophile aversion compared with SfTRPA1-CL (Figure 5B and 5C). These results are consistent with our working hypothesis that changes in TRPA1 channel physiology can drive behavioral divergence. Kang and colleagues (Kang et al. 2010) compared the chemical responses of the TRPA1-D isoform (called “TRPA1(A)” in their paper) of *D. mojavensis* and *D. virilis* to that of *D. melanogaster*. Despite some differences in the recording scheme, they reported an EC_50_ value for DmTRPA1-D similar to ours (0.278±0.024 mM (Kang et al. 2010); 0.341±0.092 mM (This study, Table 1)), while EC_50_ values for the other two species indicated higher sensitivity to chemicals, comparable to our observation for SpTRPA1-DL (i.e., DmojTRPA1-D: 0.121±0.013 mM, DvirTRPA1-D: 0.108±0.012 mM (Kang et al. 2010); SpTRPA1-DL: 0.103±0.015 mM (This study, Table 1)). A parsimonious interpretation of these patterns is that the sensitivity of TRPA1-DL(D) isoform was already high at the most recent common ancestor of the subgenus *Drosophila* (which includes *Scaptomyza*) and later it decreased only in the *S. flava* lineage. We hypothesize that the physiological evolution of TRPA1 coincided with, and potentially facilitated, the dietary transition to mustards and other Brassicales plants in *Scaptomyza*.

Altogether, results from our *in vitro* and *in vivo* experiments support the hypothesis that physiological changes to the sensitivity of the TRPA1 channel to electrophiles like AITC in part underlie gustatory evolution in *S. flava* (Figure 8). Although causal amino acid substitutions behind the physiological evolution of SfTRPA1 remain unidentified, the conservation of cysteine residues that trigger channel gating, which underwent substitutions in TRPA1 from the pain-insensitive African mole-rat (Eigenbrod et al. 2019), indicates that natural selection targeted other domains of the channel. Herbivore-specific and mustard-feeding specific substitutions in ankyrin repeat domains (ARDs) could potentially modify regulation of TRPA1 channel gating via structural changes of ARDs (Cordero-Morales et al. 2011). Functional consequences of substitutions in ARDs warrant further investigation.

**Figure 8.**
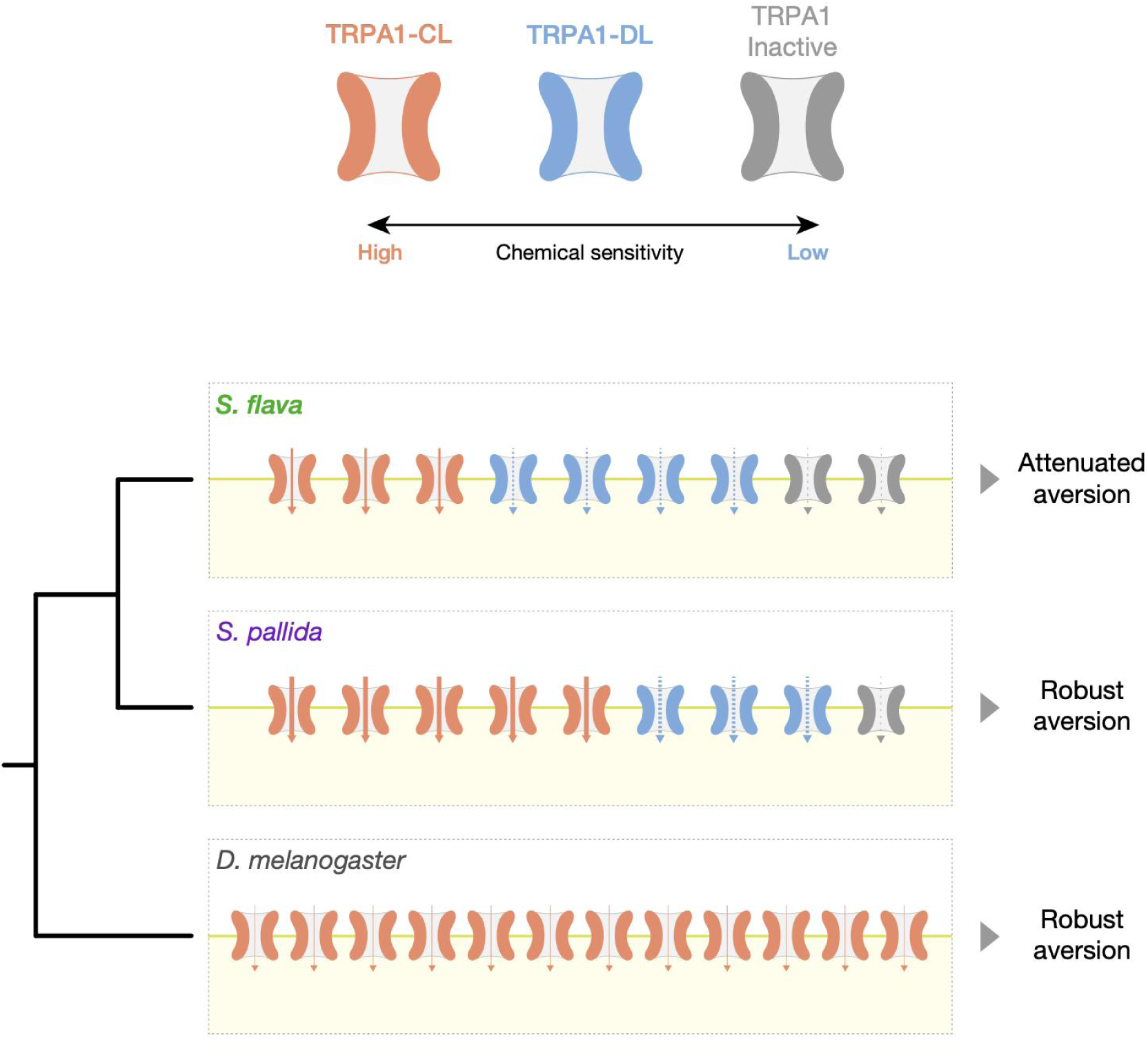
A model of sensory evolution through the combinatorial molecular changes of TRPA1 based on the results of our study. The diagram illustrates the expression patterns of TRPA1 channels in (labellar) gustatory neurons. Thickness and density of dashed arrows represent the activity of each TRPA1 isoform (i.e., degree of positively charged currents conducted by the channels) at lower ITC concentrations. Numbers of ion channels indicate the absolute expression level of *TrpA1* and the relative expression of splicing isoforms. Our analyses suggest that the weak electrophile aversion of *S. flava* is primarily attributed to the desensitization to chemical stimulus and the enrichment of less sensitive splicing isoforms of the TRPA1 channel, compared with the microbe-feeding relative *S. pallida*. In microbe-feeding *D. melanogaster*, which has the highest overall *TRPA1* gene expression levels and highest enrichment of sensitive splicing isoforms, likely compensate for the weakest activity of its TRPA1 channel and maintains its robust aversion against ITCs.

However, we also noticed that the degree of behavioral difference between the transgenic *D. melanogaster* strains expressing *TrpA1* isoforms of each species was smaller than what was observed among wild-type flies (Figure 1B). Note that our wild-type and transgenic PER assay had a few other differences that could also influence the result, including test chemicals (AITC or NMM) and the use of the binary expression system (i.e., *GAL4/UAS*) to boost the *TrpA1* gene expression in transgenic flies (Figure 5A). We also observed the behavioral response of *TrpA1*-null control flies against NMM (Figure 5B), which might suggest the presence of TRPA1-independent sensory pathways (e.g., Tracey et al. 2003; Al-Anzi et al. 2006). Nevertheless, consistently greater electrophile aversion in TRPA1-expressing lines compared with their controls (Figure 5C) supports the strong phenotypic contribution of TRPA1 under our experimental design. Therefore, we concluded from our behavioral data that the divergence of TRPA1 sequence and physiology alone does not fully explain the cross-species behavioral differences, which suggests the presence of alternative mechanisms for the lack of electrophile aversion in *S. flava*.

A more salient phenotypic difference could arise from the interspecific variation we found in the composition of *TrpA1* splicing isoforms (Figure 3) and its modulation of cellular responses to sensory stimuli *in vitro* (Figure 6 and 7). *TrpA1* isoforms are co-expressed in nociceptive C4da neurons in larval *D. melanogaster* (Gu et al. 2019). Because labellar expression of *S. flava TrpA1* was restricted to a small number of bitter-sensing neurons (Figure 2J-2L), it is reasonable to infer that these *Scaptomyza TrpA1* isoforms are co-expressed in gustatory neurons. Our oocyte electrophysiological recordings after introducing mixtures of TRPA1 channel isoforms with different response properties showed that physiological responses of a cell tended to be closer to those of predominantly expressed isoforms (Figure 6 and 7). The isoform compositions that mimicked labellar *TrpA1* expression patterns of *S. flava* and *S. pallida* produced different responses to a lower AITC concentration (Figure 6C), while the tarsal-mimicking *TrpA1* mixtures did not show a difference (Figure 6D). This suggests that isoform composition of TRPA1 channels allows fine-tuning of the dose-dependency of gustatory neurons and that the regulation of these isoforms could be subject to adaptive evolution depending on the species-specific or tissue-specific selective pressures.

We propose that proportionally higher expression of TRPA1-DL, a less electrophile-sensitive isoform, could be another mechanism facilitating the observed evolutionary shift in diet within the *S. flava* lineage. This is because elevated TRPA-DL levels could suppress feeding aversion towards Brassicales plants through the reduced activity of ITC-sensing neurons at low chemical concentrations (Figure 8). Proportionally higher expression of TRPA1-CL, on the other hand, would render ITC-sensing neurons more sensitive and could have been favored in non-mustard-feeding species like *S. pallida* to prevent them from consuming electrophilic toxins (Figure 8). This model is analogous to the enrichment of a heat-sensitive TRPV1 isoform in the vampire bat, which likely enables the sensation of infrared emitted by its mammalian prey (Gracheva et al. 2011). Expression of “inactive” TRPA1 isoforms, such as SfTRPA1-CS or SpTRPA1-EL, could further reduce sensitivity of ITC-sensing neurons, as their presence in isoform mixtures proportionately reduced current amplitudes against stimulation (Figure 7). A higher proportion of these inactive isoforms in *S. flava* than in *S. pallida* (Figure 7F) could also have contributed to the attenuation of bitter-sensing neurons against ITCs in the *S. flava* lineage. Taken together, our results suggest that the evolution of *TrpA1* isoform composition is another molecular mechanism associated with the ecological shift to Brassicales host plants in *S. flava*, likely occurring in parallel with the evolution of TRPA1 channel physiology.

One surprising finding was that DmTRPA1s produced the lowest current amplitude among the three species (Figure 4H and 4I) despite the strong feeding aversion for ITCs in *D. melanogaster* (Figure 1B). This apparent contradiction could be explained by taking into account both gene expression and physiology. First, the labellar expression level of *D. melanogaster TrpA1* was by far the highest among the three species (Figure 2A-2C). Second, *D. melanogaster* labella almost entirely lacked the expression of inactive *TrpA1* isoforms (Figure 3B). Lastly, the EC_50_ of DmTRPA1-C, which is the dominant isoform in *D. melanogaster* labella, was much lower than homologues in the other two species (Table 1). These findings together suggest that *D. melanogaster* achieves robust electrophile aversion by enriching its bitter-sensing neurons with a higher number of sensitive TRPA1 channels to compensate for the weak activity of each single channel (Figure 8). This is yet another example wherein the combination of channel physiology and gene expression of TRPA1 may explain behavioral variation. Our dataset also highlights the perils of using *D. melanogaster* as a representative, “default”, or baseline species and the benefits of using taxa representing a phylogenetic gradient and focusing on character evolution rather than living taxa as stand-ins for ancestors *per se* (Jenner 2006).

HCR RNA-FISH showed that *TrpA1* expression was restricted to bitter-sensing neurons in the *S. flava* labellum, indicating that ITCs serve as gustatory deterrents even for this mustard feeder (Figures 2G-L). This pattern reflects the dual aspects of ITCs for specialized herbivores: although volatile ITCs serve as host-associated cues (token stimuli) that allow specialist herbivores to find their food and oviposition sources (Finch 1978; Wallbank and Wheatley 1979; Matsunaga et al. 2022), they are still toxins that can delay development and will result in lethality at high doses (Lichtenstein et al. 1964; Whiteman et al. 2011). Rather than switching the valence of ITC perception to full acceptance, the mustard specialist *S. flava* may have fine-tuned the strength of bitter sensing-neuron activity so that it does not trigger aversion unnecessarily. On the other hand, the high *TrpA1* expression in *S. flava* tarsi was unexpected. Leg gustatory neurons of the flies serve as a gateway to assess the quality of a potential food source, in which plant secondary metabolites are usually perceived as aversive stimuli (Scott 2018). However, some butterflies perceive specific host plant-derived chemicals through their chemosensory neurons in the legs as attractive oviposition-stimulating cues, such as glucosinolates for the cabbage white (*Pieris rapae*) (Huang et al. 1993a; Huang et al. 1993b; Yang et al. 2021) and synephrine for the Asian swallowtail (*Papilio xuthus*) (Nishida et al. 1987; Ozaki et al. 2011; Ryuda et al. 2013). Robust *TrpA1* expression in the tarsi could potentially allow for more precise evaluation of host plant quality for *S. flava*, although this is speculative.

In summary, our findings demonstrate how evolution of *TrpA1* shapes behavioral variation across drosophilid flies through a combination of differences in overall expression levels, protein evolution, and compositional modulation of functionally diverse splicing isoforms. The latter two mechanisms are prominent contributors to the attenuated aversion of *S. flava* towards dietary electrophiles (Figure 8). To our knowledge, this is one of the first chemosensory examples in which alternative splicing of sensory channel genes may play a role in the evolution of novel behavior in animals.

While changes in receptor sensitivity are commonly observed in association with sensory evolution (Gracheva et al. 2010; McBride et al. 2014; Prieto-Godino et al. 2017; Akashi et al. 2018; Eigenbrod et al. 2019; Auer et al. 2020; Brand et al. 2020; Saito et al. 2022; Capek et al. 2025) alongside gains and losses of receptor genes (McBride 2007; McBride and Arguello 2007; Goldman-Huertas et al. 2015; Matsunaga et al. 2022; Peláez et al. 2023), alternative splicing has only rarely been associated with sensory novelty except for the case of TRPV1 in the infrared-sensing vampire bat (Gracheva et al. 2011).

This contrast can be interpreted through the lens of evolutionary constraint: large-effect mutations in functionally modular gene families, like most insect chemoreceptors, can more easily pass through the selective sieve and remain in a population, while such mutations in conserved and more pleiotropic genes are more likely to be maladaptive and therefore rarely observed as segregating or as substitutions. When evolutionary novelty arises through changes in highly conserved genes, one solution is gene duplication, which allows for retention of a functional copy of a gene to mitigate fitness costs from new mutations in the other copy, as seen in the parallel evolution of neurotoxin resistance via duplication of the insect GABA_A_ receptor gene *Rdl* (Guo et al. 2023). Similarly, modulation of alternative splicing may offer flexibility as the population explores a novel fitness landscape while maintaining core gene functions. A prediction flowing from this model is that alternative splicing provides avenues for the evolution of novel protein sequences for genes that are otherwise under strong constraint. Long-read sequencing technology and the tools of model organisms for detailed functional experiments are now allowing for discovery of whether and to what extent this mechanism gives rise to adaptive phenotypic evolution. Our findings suggest that a focus on alternative splicing (and its regulation) as a molecular mechanism for adaptation is likely to be fruitful.

## KEY RESOURCES TABLE

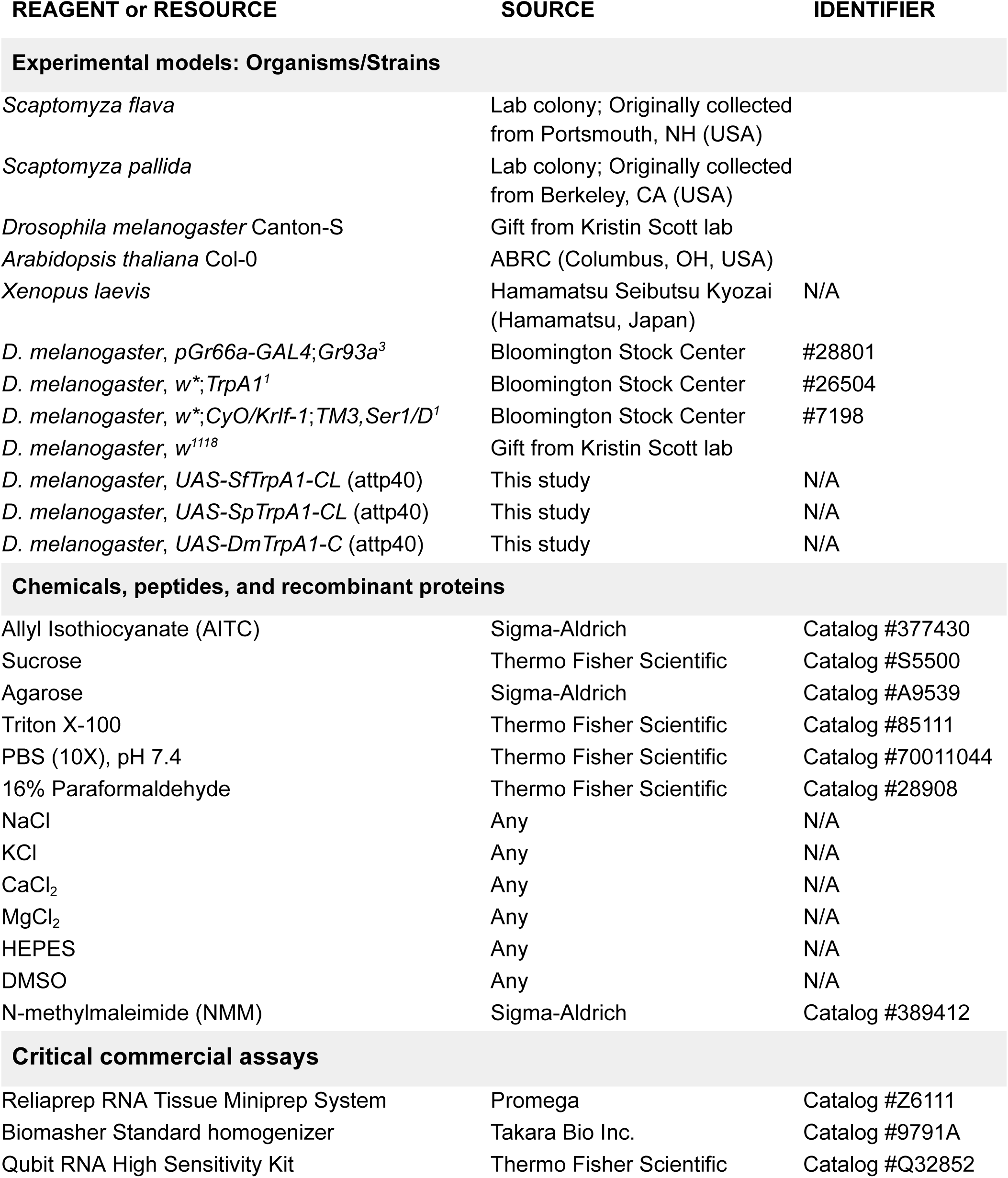
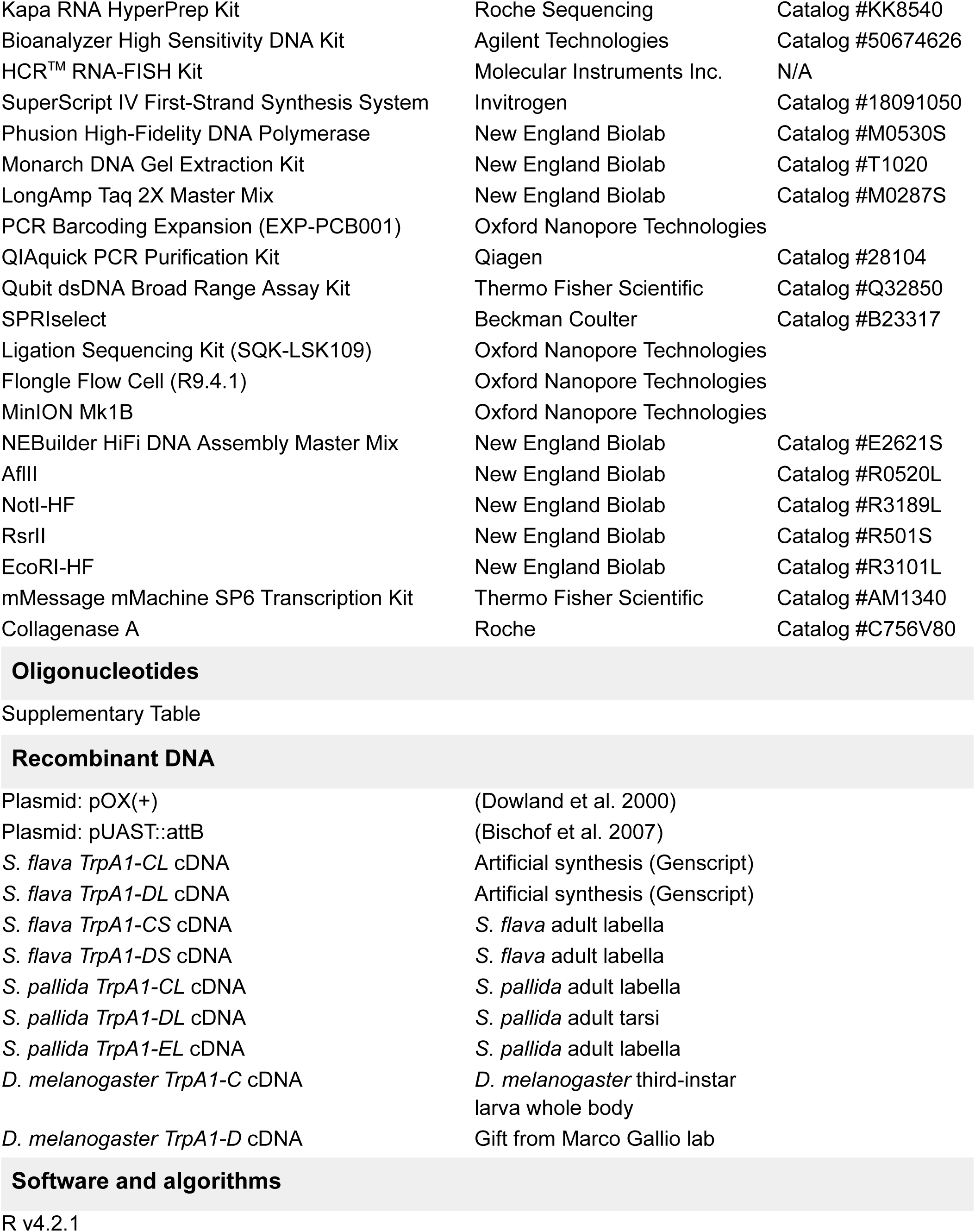
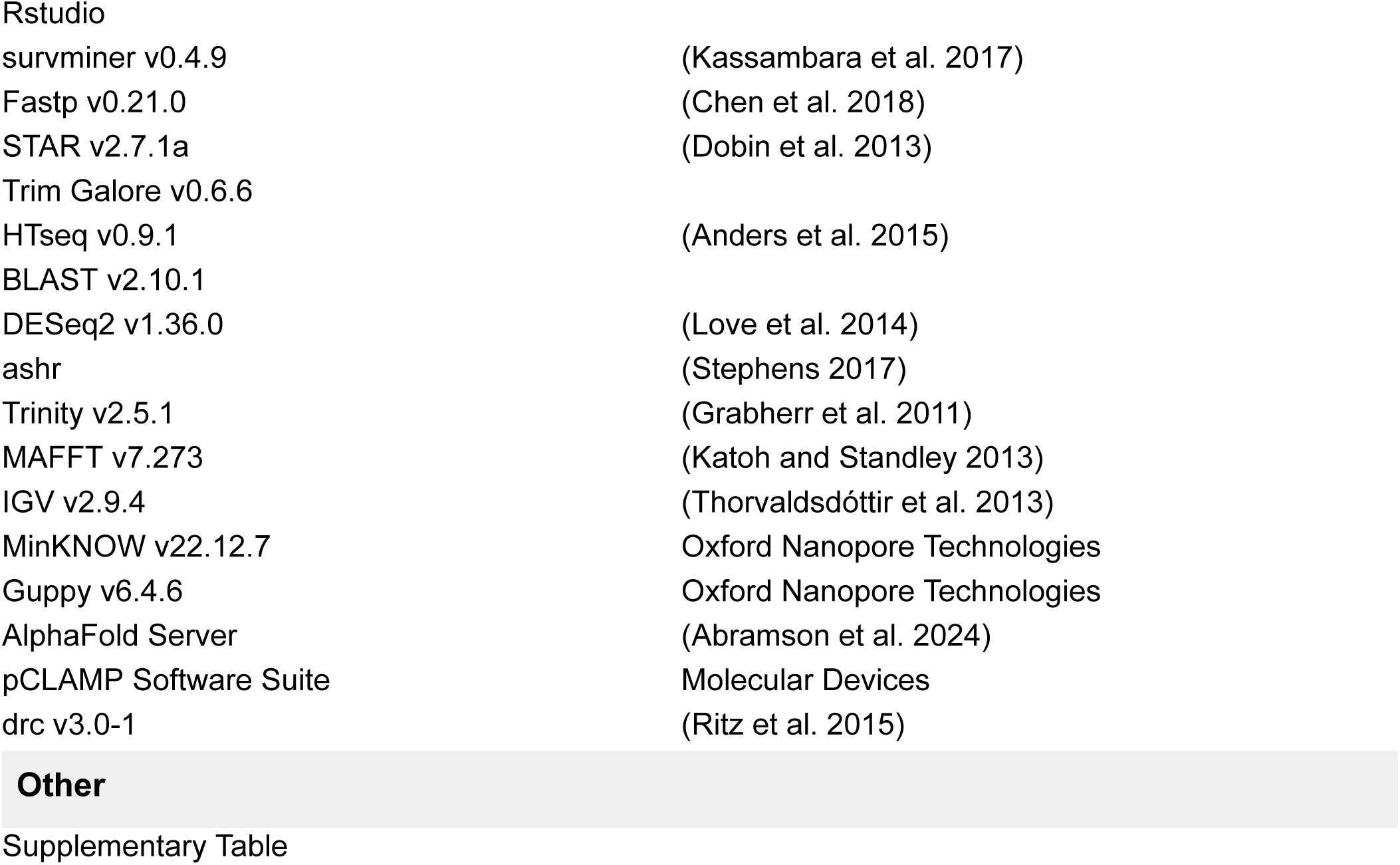

### METHOD DETAILS

#### Fly strains

The *Scaptomyza flava* laboratory colony was originally established from >150 wild larvae collected as leafminers from wild mustard plants in Portsmouth, NH (USA) in 2008. The colony was maintained in cages containing *Arabidopsis thaliana* Col-0 strain as larval and adult food source at 20-21°C and 60% relative humidity (RH) in a walk-in environmental room on a 12L:12D cycle. The *Scaptomyza pallida* lab colony was an isofemale line established from a wild female fly collected at the University of California, Berkeley campus (CA, USA) in 2017. The colony was reared on organic spinach (purchased at Trader Joe’s, CA, USA) placed on top of molasses-cornmeal media purchased from UC-Berkeley Fly Food Facility. Other rearing conditions were identical to those of *S. flava* colony. Wild-type and transgenic *Drosophila melanogaster* lines were obtained from Bloomington Drosophila Stock Center or Kristin Scott lab (University of California, Berkeley), except for the ones newly created in this study. *D. melanogaster* strains were maintained at two different locations with different settings. At UC Berkeley, flies were maintained at 24°C on a 12L:12D cycle and reared on molasses-cornmeal media from UC Berkeley Fly Food Facility. At National Institute of Physiological Sciences (NIPS), flies were maintained at 25°C on a 12L:12D cycle and reared on in-house glucose-cornmeal media (Sato et al. 2023). Flies at NIPS were used for behavioral assay with ectopic TRPA1 expression (Figure 5). All other experiments using *D. melanogaster* were performed with the strains at UC Berkeley.

#### Behavioral assay (Wild-type)

PER is a common behavior assay in *Drosophila* to measure feeding preference to a given chemical by monitoring the frequency of proboscis extension after the presentation of test chemicals (Shiraiwa and Carlson 2007). We first conducted PER assay to characterize species-specific preference towards allyl isothiocyanates (AITC). Our protocol for PER assay was based on (Kang et al. 2010) with several modifications.

Adult flies were collected from their larval media soon after the emergence and transferred to plastic vials containing a cotton ball with 10% honey water (pure organic honey purchased at Trader Joe’s, CA) to account for larval food differences. We did not separate male and female flies during this period, thus we assume that all female flies were already mated at the time of experiments. 2-9 day old flies were starved for 24 hours in a foodless humidified vial prior to the assay. Flies were then anesthetized with CO_2_, mounted on a glass slide dorsal side down using nail polish, and allowed to recover in a humidified chamber for at least 2 hours. Flies that did not show active leg movement after the 2-hour recovery period were removed from the assay. During the PER assay, all flies were first satiated with water, and each fly was offered a single type of tastant (i.e., 300 mM sucrose solution or its mixture with a given concentration of AITC) by touching their forelegs with test chemicals as a small droplet from a P200 pipette tip. We picked AITC as a representative ITC for this and subsequent experiments, because it is the most commonly used ITC in previous laboratory-based studies using *Drosophila*. Each presentation continued up to 10 seconds, and the response of a fly to the tastant within the 10-second window was scored based on our criteria: 1 pt - extended proboscis and maintained drinking the droplet >2 seconds; 0.5 pt - extended proboscis but failed to maintain contact to the droplet, or only showed brief contact; 0 pt - did not show proboscis extension and failed to contact the droplet. Feeding droplet was immediately withdrawn when a fly continued PER for 3 seconds to minimize satiety. Presentation was repeated five times per fly (with a 5-10 minute interval between presentations), such that each fly accumulated a total PER score in a range between 0 to 5, wherein the larger score indicated the stronger feeding response. We performed PER assay at four different AITC concentrations (i.e., 0, 2, 5, 10 mM AITC in a 300 mM sucrose solution), assuming the biological ITC concentrations for the Brassicales plants were somewhere between 1 mM to 5 mM. We tested n = 19-34 individual flies at each combination of species, sex, and AITC concentration (see Supplementary Table). All experiments were performed at zeitgeber time (ZT) 7-10.

To visually highlight the degree of feeding inhibition, individual PER scores at each treatment was normalized to the average PER score at 0 mM AITC within species, in which the PER score closer to 0 indicated the stronger feeding inhibition by AITC (Figure 1B and S1). The difference in feeding response across the AITC dosage series was tested within each species/sex combination using pairwise Wilcoxon rank-sum test with Benjamini-Hochberg correction using normalized individual PER scores. See Supplementary Tables for the raw data and the associations between original and normalized PER scores.

#### Generation of transgenic flies

We performed genetic rescue experiments in *TrpA1* null-mutant *D. melanogaster* lines to test whether the ectopic expression of *Scaptomyza* TRPA1 sufficiently recapitulates the behavioral variation of wild-type species. We focused our efforts on TRPA1-CL(C) isoform because it is the only predominantly expressed isoform across the three species (Figure 3B). We first digested the fly transgenesis vector pUAST::attB (Bischof et al. 2007) using NotI-HF and EcoRI-HF restriction enzymes (New England Biolab). Three *TrpA1* isoforms (*SfTrpA1-CL*, *SpTrpA1-CL*, and *DmTrpA1-C*) were subcloned into pUAST::attB using NEBuilder HiFi DNA Assembly Master Mix (New England Biolab). These *UAS-TrpA1* cassettes were inserted into *D. melanogaster* genome targeting at attP40 site on chromosome 2 using PhiC31 integrase. Transgenesis of *D. melanogaster* embryos and the initial transformant screening were performed at Genetivision (Huston, TX, USA), which successfully produced independent *UAS-TrpA1* lines for each *TrpA1* isoform. Other mutant strains were obtained from Bloomington *Drosophila* Stock Center (#28801: *pGr66a-GAL4*;*Gr93a^3^*, #26504: *w\**;*TrpA1^1^*, #7198: *w\**;*CyO/KrIf-1*;*TM3,Ser1/D^1^*). All mutant lines were initially backcrossed to *w^1118^* for five generations to homogenize their genetic background. *pGr66a-GAL4* (after omitting the *Gr93a^3^* allele) and all *UAS-TrpA1* lines were then crossed to *TrpA1^1^* (= *TrpA1* null-mutant) lines so all flies became null mutants of the endogenous *TrpA1* allele (Figure S15). Each *UAS-TrpA1*;*TrpA1^1^* line was then crossed either to *pGr66a-GAL4*;*TrpA1^1^* line (test genotype) or to *+/+*;*TrpA1^1^* line (control genotype) to obtain the genotypes we used in experiments.

#### Behavioral assay (Transgenic lines)

Procedures of the PER assay for transgenic flies were identical to those for wild-type flies except for these modifications: (1) Adult flies were collected soon after the emergence and kept on fresh media until the 24-hour starvation period. (2) 3-6 day old female flies were used for the assay (assuming all flies were mated by this period). (3) After the starvation period, individual flies were transferred into P200 plastic tips (without anesthesia) such that only the head and forelegs were exposed and allowed to access tastants. As test substances, three concentrations (0, 5, 10 mM) of N-methylmaleimide (NMM) were prepared in 100 mM sucrose solution containing 0.5% DMSO. We switched from AITC that could interfere with the endogenous olfactory pathway of flies (Matsunaga et al. 2025), and instead used another TRPA1 activator, NMM, such that we can evaluate the effect of ectopic TRPA1 expression on gustatory behavior. (4) Each fly was tested with a single tastant five times, in which each presentation lasted up to 5 seconds. All assays were performed at ZT 6.5-9.5.

PER scores were calculated in the same criteria as those in wild-type PER, which ranged from 0 to 5 over the five trials. These raw PER scores were eventually normalized to the average PER score at 0 mM NMM within each genotype. Differences in the distribution of total PER scores were statistically tested within genotypes using pairwise Wilcoxon rank-sum test with Benjamini-Hochberg correction based on normalized individual PER scores. To highlight the contribution of ectopic TRPA1 expression, we further took the ratio of average PER scores between a test genotype (*Gr66a>TrpA1*;*TrpA1^1^*) and a control genotype (*UAS-TrpA1/+*;*TrpA1^1^*) for each TRPA1 channel expressed, and plotted the dose-responses as line graphs (Figure 5C). Error bars were not applicable for the ratio because there was only one value for each genotype and NMM concentration. See Supplementary Tables for the raw data and the associations between original and normalized PER scores.

#### Starvation tolerance assay

Adult flies were collected upon emergence and reared on 10% honey water. 8-10 flies (of 2-5 days old) of each species/sex were grouped and transferred to a plastic vial containing 1% agarose, and incubated under a laboratory condition (20-21 °C, 60% humidity, 12:12 L:D cycle). We then monitored the number of dead/live flies every 12 hours until all individuals were dead. 5-6 replicates for each species/sex group were tested. Survival rates differences between species were analyzed for each sex by pairwise log-rank test using survminer package v0.4.9 (Kassambara et al. 2017) in R v4.2.1.

#### RNA extraction and RNA-sequencing

Adult females of *S. flava* and *S. pallida* were collected immediately after eclosion from the original vials and transferred to humidified vials containing 10% honey water for up to 3 days. Labella and tarsi were hand-dissected using razors and forceps while the flies were anesthetized by CO_2_. Dissected tissues were then collected in LB+TGA lysis buffer from Reliaprep RNA Tissue Miniprep System (Promega, USA) and disrupted using Biomasher Standard homogenizer (Takara Bio Inc., USA) in a dry ice ethanol bath. Approximate numbers of individuals pooled for a single replicate were 100-120 for labella and 70-75 for tarsi (forelegs, midlegs, and hindlegs were pooled). We prepared three biological replicates for each combination of species and tissue. Sample lysates were stored in −80 °C freezer until subsequent RNA extraction.

Total RNA were extracted from the lysates using ReliaPrep RNA Tissue Miniprep System (Promega) according to the manufacturer’s protocol, and quantified using Qubit RNA High Sensitivity kit (Thermo Fisher Scientific). Extracted RNA samples were subsequently processed into cDNA libraries using KAPA RNA HyperPrep Kit (Roche Sequencing) according to the manufacturer’s protocol with a few modifications, such as the use of in-house indexing adapters developed by the Functional Genomics Laboratory (FGL), a QB3-Berkeley Core Research Facility at UC Berkeley. Approximately 300 ng total RNA for labellar samples and 100 ng for tarsal samples were used as a starting material per library. Fragment lengths of cDNA libraries were quantified with Bioanalyzer High sensitivity DNA kit (Agilent Technologies, USA) and sequenced on NovaSeq 6000 platforms (Illumina, USA) for 150 bp paired-end reads at Genome Sequencing Laboratory (GSL) at UC Berkeley. Raw RNA-seq reads were deposited to NCBI SRA database under BioProject accession number PRJNA779814.

#### Differential gene expression analysis

We conducted differential gene expression (DGE) analyses to compare and characterize the expression levels of *TrpA1* across species. We filtered raw RNA-seq reads using Fastp v0.21.0 (Chen et al. 2018) and mapped the clean reads on reference genomes for *S. flava* (Peláez et al. 2023) and *S. pallida* (Kim et al. 2021) using STAR v2.7.1a (Dobin et al. 2013) to generate BAM files. We also downloaded labellar and tarsal RNA-sequencing datasets of *D. melanogaster* female (Dweck et al. 2021; Wang et al. 2022) and mapped them onto *D. melanogaster* reference genome (BDGP6.46) in the same manner. Since *D. melanogaster* short reads were 75 bp long, we created another set of BAM files for *S. flava* by trimming the reads into 75 bp using Trim Galore v0.6.6 for the expression analysis between *D. melanogaster* and *S. flava*, to minimize potential biases due to read length differences (Chhangawala et al. 2015). BAM files were subsequently converted into count data using HTseq v0.9.1 (Anders et al. 2015).

We next performed reciprocal BLASTN searches using the whole set of protein coding sequences of *S. flava* and *S. pallida* to identify 1-to-1 orthologous relationships between species. We screened gene pairs by bitscores and removed redundant genes using manual Python scripts, resulting in 10,439 orthologous gene pairs between *S. flava* and *S. pallida*. Similar procedures yielded 9,563 orthologous gene pairs between *S. flava* and *D. melanogaster*. Principal component analysis (PCA) was conducted on 8,920 orthologous genes to visualize similarities among samples using prcomp package in R v4.2.1. Cross-species DGE analysis were performed using DESeq2 v1.36.0 (Love et al. 2014) with ashr for Log Fold Shrinkage (Stephens 2017). Because of the high false-positive rates intrinsic to cross-species analyses, we applied a strict cutoff (adjusted p-value < 0.001) to identify differentially expressed genes between species. We calculated TPM (transcripts per million reads) of the *TrpA1* gene based on the count data and gene lengths for each species.

#### Fluorescent *in situ* hybridization

We conducted RNA FISH to examine the types of gustatory neurons expressing *TrpA1* in the fly labellum. HCR (hybridization chain reaction) RNA FISH kit was ordered from Molecular Instruments (CA, USA) (Choi et al. 2018). Custom mRNA probes for *D. melanogaster* and *S. flava TrpA1,* as well as *Gr66a* as a marker for bitter-sensing neurons, were designed and produced at Molecular Instruments (CA, USA). *TrpA1* probes targeted Exons 5-11 so they bind to all splicing isoforms equally. Female *D. melanogaster* and *S. flava* between 3-10 days old were anesthetized with CO_2_. Their mouthparts were hand-dissected with fine forceps and fixed in a 2 mL fixative solution (4% paraformaldehyde in 1X PBS, with 0.1% Triton X-100) at 4 °C on a nutator for 24 hours. Following the fixation, samples were washed twice in 2 mL 1X PBS + 3% Triton X-100, once in 2 mL 1X PBT (i.e., PBS + 0.1% Triton X-100), and four times in 1 mL 1X PBT. Hybridization and amplification steps were mostly identical to the manufacturer’s protocol for *D. melanogaster* embryos, except for the higher probe/hairpin concentrations suggested in (Bontonou et al. 2024): probe solutions were created by adding 5 μL each of experimental (*TrpA1*) and control (*Gr66a*) probes to 300 μL probe hybridization buffer. Likewise, hairpin solutions were prepared by adding 10 μL of each 3 μM hairpin stock to 300 μL amplification solution.

Samples were imaged with a Zeiss LSM 880 microscope using an Airyscan detector and a 63x/1.4NA oil immersion objective. *Gr66a* and *TrpA1* mRNAs were visualized using the 488 nm (3% power) and 561 nm (2% power) excitation lasers, respectively. Z-stacks were acquired with 944 x 944 pixel image resolution, with a pixel size of 0.07 µm and 0.2 µm step size. After acquisition in Airyscan Fast mode, raw images were deconvolved using the Zen Black software. Airyscan-processed images were subsequently analyzed using Fiji (ImageJ).

#### Detection of *TrpA1* splicing variants and orthologues

To predict mRNA structures of *Scaptomyza TrpA1* splice variants, we first conducted *de novo* assembly of aforementioned RNA-seq short reads using Trinity v2.5.1 (Grabherr et al. 2011), and subsequently ran TBLASTN searches (v2.10.1) against the assembled contigs of *S. flava* and *S. pallida* using *D. melanogaster* TRPA1-C protein sequence as a query. Exonic structures of *S. flava* and *S. pallida TrpA1s* were manually inspected from sequence alignments using MAFFT v7.273 (Katoh and Standley 2013) and aforementioned BAM flies using IGV v2.9.4 (Thorvaldsdóttir et al. 2013) after the BLAST search against their respective reference genomes. Presence of a certain *TrpA1* isoforms was further confirmed by PCR-based subcloning using labellar and tarsal cDNA samples of the two *Scaptomyza* species.

*TrpA1* orthologues from other species were obtained from the NCBI database or detected by TBLASTN searches against reference genomes using *S. flava* TRPA1-CL as a query. Putative exons at the N-terminal region of *D. melanogaster*, *D. mojavensis*, *D. grimshawi*, *S. hsui*, *S. graminum, and S. montana TrpA1s* were also identified based on BLAST searches and manual inspection of their genomic sequences. For *D. melanogaster* and *D. mojavensis*, NCBI-deposited RNA-seq data were mapped on their genomes using STAR v2.7.1a (Dobin et al. 2013) and referred as additional evidence for the transcription of *TrpA1* splicing variants.

#### Amplicon sequencing and analyses of isoform compositions

We next aimed to quantify the proportions of expressed *TrpA1* isoforms in fly labella and tarsi to investigate whether the isoform compositions are associated with the dietary adaptation of the flies. Due to the nature of *Scaptomyza/Drosophila TrpA1* isoforms where exons defining isoform types are distantly located (i.e., ∼2 kb on mRNA, Figure 3A), common gene expression analyses such as short-read sequencing or quantitative PCR do not accurately capture the proportions of *TrpA1* splicing variants. Therefore, we employed amplicon sequencing targeting the entire *TrpA1* cDNA using a long-read sequencer MinION (Oxford Nanopore Technologies), assuming the ratio of *TrpA1* isoforms was still conserved after PCR amplification (Figure S7).

Our experimental procedures are modified from those introduced previously published (Clark et al. 2020; Hall et al. 2021). Total RNA was extracted from independent groups of fly labella or tarsi (forelegs, midlegs, and hindlegs were pooled), wherein each replicate contained roughly 40-50 individuals. 100 ng labellar total RNA or 50 ng tarsal total RNA were transcribed into cDNA using SuperScript IV (Invitrogen, USA). These cDNAs were later used as templates for the first-round PCR for *TrpA1* transcripts using Phusion High-Fidelity DNA polymerase (New England Biolab). PCR primers were designed for the first exon (i.e., Exon L; usually considered a 5’UTR in *Drosophila TrpA1*) and the last exons (i.e., Exon 18) such that the entire pool of *TrpA1*isoforms can be amplified in a single reaction (Figure 3A, Figure S7). Furthermore, the primers were concatenated to common “adapter” sequences acting as annealing sites for barcoding primers in the second-round barcoding PCR (Figure S7). Successful *TrpA1* amplicons were isolated from 1% agarose gel after electrophoresis and purified using Monarch DNA gel extraction kit (New England Biolab). Possibly because of its low expression levels, *TrpA1* amplicons from *D. melanogaster* tarsal cDNA were far less abundant and required additional rounds of PCR amplification(Figure 2F). To avoid technical inconsistency, we omitted *D. melanogaster* tarsi from the downstream analysis. 40 ng *TrpA1* amplicons were then used as templates for the second-round barcoding PCR using LongAmp Taq 2X Master Mix (New England Biolab) and PCR Barcoding Expansion (EXP-PBC001, Oxford Nanopore Technologies). Barcoded *TrpA1* fragments were cleaned with QIAquick PCR Purification Kit (Qiagen) and quantified with Qubit dsDNA BR Assay Kit (Thermo Fisher Scientific). Subsequently, multiple *TrpA1* amplicons were pooled together in a single tube for size selection using X0.6 SPRIselect magnetic beads (Beckman Coulter, USA). Selected fragments were used for library preparation using Ligation Sequencing Kit (SQK-LSK109, Oxford Nanopore Technologies). Libraries were sequenced on Flongle Flow Cell (R9.4.1; Oxford Nanopore Technologies) with MinION Mk1B sequencer (Oxford Nanopore Technologies) and MinKNOW platform v22.12.7. Basecalling was concurrently performed with sequencing using Guppy v6.4.6.

After obtaining sequencing output as FASTQ files, we performed BLASTN searches against the outputs using pre-annotated *TrpA1* isoform sequences as queries to figure out the isoform type of each single read. To take into account the high error rates intrinsic to ONT long-read sequencers, we used in-house Python scripts to filter out the reads where the lengths were outside the range between 3.8 and 4.5 kb or the sequence similarity to the best hit gene model was less than 85%. Passed reads were categorized as their best-hit *TrpA1* isoform type, and the proportions of the isoforms were calculated. The relative proportions of each *TrpA1* isoform type were fitted into a generalized linear model (GLM) assuming Gaussian distribution using R v4.2.1, and the difference between groups were tested with Tukey’s post hoc HSD test with Benjamini-Hochberg correction.

#### Structural analysis

We manually inspected the multiple sequence alignment of TRPA1 to identify herbivore-specific substitutions. We predicted structures of *S. flava* and *S. pallida* TRPA1s using the AlphaFold Server (Abramson et al. 2024) based on their amino acid sequences. Putative structural changes by amino acid substitutions were investigated by simply editing the residues on input sequences.

#### Oocyte electrophysiology

We performed two-electrode voltage clamp (TEVC) recordings using *Xenopus laevis* oocytes to characterize activity and sensitivity of detected TRPA1 isoforms. A *Xenopus* expression vector pOX(+) (Dowland et al. 2000) was first digested by restriction enzymes AflII and NotI-HF (New England Biolab). Nine *Scaptomyza*/*Drosophila TrpA1* isoform CDSs were then subcloned into pOX(+) using NEBuilder HiFi DNA Assembly Master Mix (New England Biolab). Original *TrpA1* sequences were obtained from various sources (see Key Resource Table). cDNA synthesis was conducted with 40-80 ng starting total RNA using Superscript IV (Invitrogen). pOX(+)-*TrpA1* constructs were linearized with restriction enzyme Rsrll (New England Biolab) and used as templates for cRNA (complementary RNA) synthesis using mMessage mMachine SP6 Transcription kit (Thermo Fisher Scientific). All TRPA1 cRNAs were diluted to 50 ng/µL concentration.

Oocytes were collected from adult *Xenopus laevis* and processed with Collagenase A (Roche) to remove follicular membranes. 50 nL of diluted cRNA (50 ng/µL) was injected into each single oocyte 1-2 days after egg retrieval. Two-electrode voltage clamp (TEVC) recordings were performed 2-3 days post injection using OC-725C amplifier (Warner Instruments, USA) with a 1 kHz low-pass filter and digitized at 5 kHz Digidata 1440 (Molecular Devices, USA). Oocytes were positioned in a bath-filled oocyte chamber and clamped at −60 mV during the recordings. ND96 (96 mM NaCl, 2 mM KCl, 1.8 mM CaCl_2_, 1 mM MgCl_2_, and 5 mM HEPES; pH 7.4) was used as the bath solution during the entire recordings. Bath temperature was monitored using a thermistor in an oocyte chamber and TC-344B dual channel temperature controller (Warner Instruments). AITC for chemical stimulation was first prepared in DMSO at 0.001, 0.01, 0.03, 0.1, 0.3, 1, and 3 M concentrations, each of which was further diluted 1,000-fold with bath solution prior to the recording. Heat stimulation was conducted by perfusing heated bath solution to the chamber. Taking advantage of the polymodal nature of fly TRPA1, we stimulated the channels twice in the course of a single recording. Oocytes were first heated (>40 °C) for 30 seconds, washed for 120 seconds, and then challenged with a single dosage of AITC for 90 seconds. This design allowed us to normalize a peak amplitude of an AITC-evoked raw current (*I_AITC_*) by the amplitude of heat-evoked raw current (*I_Heat_*). When the current response did not saturate within the 90-second AITC stimulation, *I_AITC_* was considered as the amplitude at the end of the 90-second window. Recordings were performed at a series of concentrations ranging from 0.001 mM to 3 mM AITC. Raw current values were retrieved using Clampfit v11.2.0.61 (Molecular Devices). Normalized current value (*I_AITC_*/*I_Heat_*) from each recording was used to estimate dose-response relationships and EC_50_ for each TRPA1 isoform using R package drc v3.0-1 (Ritz et al. 2015) under the LL.3 function. To investigate potential biases due to the difference of heat activation, *I_Heat_* from all recordings were merged for each canonical isoform type (i.e., TRPA1-CL or TRPA1-DL) and analyzed between species using Kruskal-Wallis test, followed by Conover’s post hoc test with Benjamini-Hochberg correction using R v4.2.1.

TEVC recordings with the co-expression of multiple TRPA1 isoforms were performed in the same manner, except that diluted 50 ng/µL cRNAs were mixed in a given ratio prior to the microinjection. Expression ratios of *TrpA1-CL* to *TrpA1-DL* mRNA in four tissues (i.e., *S. flava* female labella, *S. pallida* labella, *S. flava* female tarsi, *S. pallida* tarsi) were determined by taking an average of three biological replicates of amplicon read count data (Figure 3B) and approximating it to the closest integer ratio. Note that we used raw AITC-evoked currents (*I_AITC_*) for the analyses of inactive TRPA1 isoforms (i.e., SfTRPA1-CS, SfTRPA1-DS, SpTRPA1-EL) because their small heat-evoked currents (*I_Heat_*) prevented accurate normalization. Differences in labellar or tarsal mRNA mixtures between species (Figure 6C and 6D) were tested by Student’s t-test or Welch’s t-test depending on the fulfillment of parametric assumptions. Differences among the isoform mixtures within a single species (i.e., canonical/inactive isoform mixtures (Fig. 7D and 7E) and single isoforms vs. labellar/tarsal mixtures (Figure 6A and 6B)) were analyzed using one-way ANOVA followed by Tukey’s post hoc test, or Kruskal-Wallis test followed by Conover’s post-hoc test with Benjamini-Hochberg correction, also depending on the fulfillment of parametric assumptions. All statistical analyses were run on R v4.2.1.

## ACKNOWLEDGEMENTS

We thank Carolina Reisenman, Kirsten Verster, Rebecca Tarnopol, Marianthi Karageorgi, Lydia Smith, Rebecca Duncan, Susan Bernstein, Jessica Aguilar, Benjamin Goldman-Huertas, Samridhi Chaturvedi, Moe Bakhtiari, Gary Karpen, Meghan Laturney, Diana Bautista, Oscar Arenas, Juliana Valencia Lesmes, Rebecca Tarvin, Kristin Scott (University of California, Berkeley), Shoma Sato (National Institute of Physiological Sciences, Japan), Lucia Prieto-Godino, Juan Lugo Ramos (Francis Crick Institute), Gwénaëlle Bontonou, Roman Arguello (University of Lausanne), Ryoya Tanaka, Yuki Ishikawa (Nagoya University), Hiroki Takahashi (National Institute of Basic Biology, Japan), KyeongJin Kang (Korea Brain Research Institute), and Paul Garrity (Brandeis University) for their experimental support and helpful discussions. We thank Marco Gallio and Alessia Para (Northwestern University) for sharing with us the dTRPA1-D construct. We thank David Julius and John Lin-King (University of California, San Francisco) for their early support for this project and in facilitating pilot experiments. This project was primarily supported by funding from the National Institute of General Medical Sciences of the National Institutes of Health Grant (award no. R35GM119816) to N.K.W. S.R. was supported by the NIH Grant (award no. R35GM139653) to Gary H. Karpen. S.C.G. was supported by the NIH Grant (award no. R35GM151194) awarded to S.C.G.

## AUTHOR CONTRIBUTIONS

H.C.S., S.C.G., and N.K.W. conceived the project. H.C.S., S.S., and N.K.W. designed the experiments with major contributions from D.H. (on *TrpA1* amplicon sequencing), S.R. (on HCR RNA-FISH), and T.S. (on transgenic fly crossing and subsequent PER assay). H.C.S. performed all behavioral experiments, bioinformatics, and statistical analyses. H.C.S., J.N.P., T.M., and A.S.T. performed fly tissue dissection, RNA extraction, and molecular biology, and K.M.T. and A.T. supported the experiments. H.C.S. and C.T.S. performed oocyte electrophysiology and data analyses under the supervision of S.S. and M.T. S.R. performed microscopy. H.C.S. and N.K.W. drafted and revised the manuscript, and all authors provided comments and contributed to editing the manuscript.

## REFERENCES

Abramson J, Adler J, Dunger J, Evans R, Green T, Pritzel A, Ronneberger O, Willmore L, Ballard AJ, Bambrick J, et al. 2024. Accurate structure prediction of biomolecular interactions with AlphaFold 3. Nature 630:493–500.

Aguilar JM, Gloss AD, Suzuki HC, Verster KI, Singhal M, Hoff J, Grebenok R, Nabity PD, Behmer ST, Whiteman NK. 2024. Insights into the evolution of herbivory from a leaf-mining fly. Ecosphere 15:e4764.

Akashi HD, Saito S, Cádiz Díaz A, Makino T, Tominaga M, Kawata M. 2018. Comparisons of behavioural and TRPA1 heat sensitivities in three sympatric Cuban Anolis lizards. Mol. Ecol. 27:2234–2242.

Al-Anzi B, Tracey WD, Benzer S. 2006. Response of Drosophila to Wasabi Is Mediated by painless, the Fly Homolog of Mammalian TRPA1/ANKTM1. Curr. Biol. 16:1034–1040.

Álvarez-Ocaña R, Shahandeh MP, Ray V, Auer TO, Gompel N, Benton R. 2023. Odor-regulated oviposition behavior in an ecological specialist. Nat. Commun. 14:3041.

Anders S, Pyl PT, Huber W. 2015. HTSeq--a Python framework to work with high-throughput sequencing data. Bioinformatics 31:166–169.

Arenas OM, Zaharieva EE, Para A, Vásquez-Doorman C, Petersen CP, Gallio M. 2017. Activation of planarian TRPA1 by reactive oxygen species reveals a conserved mechanism for animal nociception. Nat. Neurosci. 20:1686–1693.

Auer TO, Khallaf MA, Silbering AF, Zappia G, Ellis K, Álvarez-Ocaña R, Arguello JR, Hansson BS, Jefferis GSXE, Caron SJC, et al. 2020. Olfactory receptor and circuit evolution promote host specialization. Nature 579:402–408.

Bendesky A, Bargmann CI. 2011. Genetic contributions to behavioural diversity at the gene–environment interface. Nat. Rev. Genet. 12:809–820.

Bischof J, Maeda RK, Hediger M, Karch F, Basler K. 2007. An optimized transgenesis system for Drosophila using germ-line-specific phiC31 integrases. Proc. Natl. Acad. Sci. U. S. A. 104:3312–3317.

Bontonou G, Saint-Leandre B, Kafle T, Baticle T, Hassan A, Sánchez-Alcañiz JA, Arguello JR. 2024. Evolution of chemosensory tissues and cells across ecologically diverse Drosophilids. Nat. Commun. 15:1047.

Brand P, Hinojosa-Díaz IA, Ayala R, Daigle M, Yurrita Obiols CL, Eltz T, Ramírez SR. 2020. The evolution of sexual signaling is linked to odorant receptor tuning in perfume-collecting orchid bees. Nat. Commun. 11:244.

Capek M, Arenas OM, Alpert MH, Zaharieva EE, Méndez-González ID, Simões JM, Gil H, Acosta A, Su Y, Para A, et al. 2025. Evolution of temperature preference in flies of the genus Drosophila. Nature [Internet]. Available from: 10.1038/s41586-025-08682-z

Chen S, Zhou Y, Chen Y, Gu J. 2018. fastp: an ultra-fast all-in-one FASTQ preprocessor. Bioinformatics 34:i884–i890.

Chhangawala S, Rudy G, Mason CE, Rosenfeld JA. 2015. The impact of read length on quantification of differentially expressed genes and splice junction detection. Genome Biol. 16:1–10.

Choi HMT, Schwarzkopf M, Fornace ME, Acharya A, Artavanis G, Stegmaier J, Cunha A, Pierce NA. 2018. Third-generation in situ hybridization chain reaction: multiplexed, quantitative, sensitive, versatile, robust. Development 145:dev165753.

Clark MB, Wrzesinski T, Garcia AB, Hall NAL, Kleinman JE, Hyde T, Weinberger DR, Harrison PJ, Haerty W, Tunbridge EM. 2020. Long-read sequencing reveals the complex splicing profile of the psychiatric risk gene CACNA1C in human brain. Mol. Psychiatry 25:37–47.

Cordero-Morales JF, Gracheva EO, Julius D. 2011. Cytoplasmic ankyrin repeats of transient receptor potential A1 (TRPA1) dictate sensitivity to thermal and chemical stimuli. Proceedings of the National Academy of Sciences 108:E1184–E1191.

Diaz F, Allan CW, Matzkin LM. 2018. Positive selection at sites of chemosensory genes is associated with the recent divergence and local ecological adaptation in cactophilic Drosophila. BMC Evol. Biol. 18:144.

Dobin A, Davis CA, Schlesinger F, Drenkow J, Zaleski C, Jha S, Batut P, Chaisson M, Gingeras TR. 2013. STAR: ultrafast universal RNA-seq aligner. Bioinformatics 29:15–21.

Dowland LK, Luyckx VA, Enck AH, Leclercq B, Yu AS. 2000. Molecular cloning and characterization of an intracellular chloride channel in the proximal tubule cell line, LLC-PK1. J. Biol. Chem. 275:37765–37773.

Duckworth RA. 2009. The role of behavior in evolution: a search for mechanism. Evol. Ecol. 23:513–531.

Du EJ, Lee M, Kim SY, Park SH, Ohk H-J, Kang K. 2024. Linkage of alternative exon assembly in Drosophila TrpA1 transcripts. Mol. Cells 47:100110.

Dweck HK, Talross GJ, Wang W, Carlson JR. 2021. Evolutionary shifts in taste coding in the fruit pest Drosophila suzukii. Elife [Internet] 10. Available from: 10.7554/eLife.64317

Eigenbrod O, Debus KY, Reznick J, Bennet NC, Sánchez-Carranza O, Omerbasic D, Hart DW, Barker AJ, Zhong W, Lutermann H, et al. 2019. Rapid molecular evolution of pain insensitivity in multiple African rodents. Science 859:852–859.

Finch S. 1978. VOLATILE PLANT CHEMICALS AND THEIR EFFECT ON HOST PLANT FINDING BY THE CABBAGE ROOT FLY (DELIA BRASSICAE). Entomol. Exp. Appl. 24:150–159.

Gloss AD, Dittrich ACN, Lapoint RT, Huertas BG, Verster KI, Pelaez JL, Nelson ADL, Aguilar J, Armstrong E, Charboneau JLM, et al. 2019. Evolution of herbivory remodels a Drosophila genome. bioRxiv:767160.

Gloss AD, Vassão DG, Hailey AL, Nelson Dittrich AC, Schramm K, Reichelt M, Rast TJ, Weichsel A, Cravens MG, Gershenzon J, et al. 2014. Evolution in an ancient detoxification pathway is coupled with a transition to herbivory in the drosophilidae. Mol. Biol. Evol. 31:2441–2456.

Goldman-Huertas B, Mitchell RF, Lapoint RT, Faucher CP, Hildebrand JG, Whiteman NK. 2015. Evolution of herbivory in Drosophilidae linked to loss of behaviors, antennal responses, odorant receptors, and ancestral diet. Proceedings of the National Academy of Sciences 112:3026–3031.

Grabherr MG, Haas BJ, Yassour M, Levin JZ, Thompson DA, Amit I, Adiconis X, Fan L, Raychowdhury R, Zeng Q, et al. 2011. Trinity: reconstructing a full-length transcriptome without a genome from RNA-Seq data. Nat. Biotechnol. 29:644–652.

Gracheva EO, Cordero-Morales JF, González-Carcacía JA, Ingolia NT, Manno C, Aranguren CI, Weissman JS, Julius D. 2011. Ganglion-specific splicing of TRPV1 underlies infrared sensation in vampire bats. Nature 476:88–92.

Gracheva EO, Ingolia NT, Kelly YM, Cordero-Morales JF, Hollopeter G, Chesler AT, Sánchez EE, Perez JC, Weissman JS, Julius D. 2010. Molecular basis of infrared detection by snakes. Nature 464:1006–1011.

Guo L, Qiao X, Haji D, Zhou T, Liu Z, Whiteman NK, Huang J. 2023. Convergent resistance to GABA receptor neurotoxins through plant–insect coevolution. Nature Ecology & Evolution:1–13.

Gu P, Gong J, Shang Y, Wang F, Ruppell KT, Ma Z, Sheehan AE, Freeman MR, Xiang Y. 2019. Polymodal Nociception in Drosophila Requires Alternative Splicing of TrpA1. Curr. Biol. 29:3961–3973.e6.

Gu P, Wang F, Shang Y, Liu J, Gong J, Xie W, Han J, Xiang Y. 2022. Nociception and hypersensitivity involve distinct neurons and molecular transducers in Drosophila. Proc. Natl. Acad. Sci. U. S. A. 119:e2113645119.

Hall NAL, Husain SM, Lee H, Tunbridge EM. 2021. Long read transcript profiling of ion channel splice isoforms. Methods Enzymol. 654:345–364.

Himmel NJ, Cox DN. 2020. Transient receptor potential channels: current perspectives on evolution, structure, function and nomenclature. Proc. Biol. Sci. 287:20201309.

Hinman A, Chuang H-H, Bautista DM, Julius D. 2006. TRP channel activation by reversible covalent modification. Proc. Natl. Acad. Sci. U. S. A. 103:19564–19568.

Huang X, Renwick JAA, Sachdev-Gupta K. 1993a. A chemical basis for deifferential acceptance of Erysimum cheiranthoides by two Pieris species. J. Chem. Ecol. 19:195–210.

Huang X, Renwick JAA, Sachdev-Gupta K. 1993b. Oviposition stimulants and deterrents regulating differential acceptance of Iberis amara by Pieris rapae and P. napi oleracea. J. Chem. Ecol. 19:1645–1663.

Inagaki HK, Panse KM, Anderson DJ. 2014. Independent, reciprocal neuromodulatory control of sweet and bitter taste sensitivity during starvation in Drosophila. Neuron 84:806–820.

Jenner RA. 2006. Unburdening evo-devo: ancestral attractions, model organisms, and basal baloney. Dev. Genes Evol. 216:385–394.

Jordt SE, Bautista DM, Chuang HH, McKemy DD, Zygmunt PM, Högestätt ED, Meng ID, Julius D. 2004. Mustard oils and cannabinoids excite sensory nerve fibres through the TRP channel ANKTM1. Nature 427:260–265.

Kang K, Panzano VC, Chang EC, Ni L, Dainis AM, Jenkins AM, Regna K, Muskavitch MAT, Garrity PA. 2012. Modulation of TRPA1 thermal sensitivity enables sensory discrimination in Drosophila. Nature 481:76–81.

Kang K, Pulver SR, Panzano VC, Chang EC, Griffith LC, Theobald DL, Garrity PA. 2010. Analysis of Drosophila TRPA1 reveals an ancient origin for human chemical nociception. Nature 464:597–600.

Karageorgi M, Bräcker LB, Lebreton S, Minervino C, Cavey M, Siju KP, Grunwald Kadow IC, Gompel N, Prud’homme B. 2017. Evolution of Multiple Sensory Systems Drives Novel Egg-Laying Behavior in the Fruit Pest Drosophila suzukii. Curr. Biol. 27:847–853.

Kassambara A, Kosinski M, Biecek P, Fabian S. 2017. Package “survminer.” Drawing Survival Curves using “ggplot2*”(R package version 03 1)* 3.

Katoh K, Standley DM. 2013. MAFFT multiple sequence alignment software version 7: Improvements in performance and usability. Mol. Biol. Evol. 30:772–780.

Khallaf MA, Auer TO, Grabe V, Depetris-Chauvin A, Ammagarahalli B, Zhang D-D, Lavista-Llanos S, Kaftan F, Weißflog J, Matzkin LM, et al. 2020. Mate discrimination among subspecies through a conserved olfactory pathway. Science Advances 6:eaba5279.

Kim BY, Wang JR, Miller DE, Barmina O, Delaney E, Thompson A, Comeault AA, Peede D, D’Agostino ER, Pelaez J, et al. 2021. Highly contiguous assemblies of 101 drosophilid genomes. Elife [Internet] 10. Available from: 10.7554/eLife.66405

Lapoint RT, O’Grady PM, Whiteman NK. 2013. Diversification and dispersal of the Hawaiian Drosophilidae: The evolution of Scaptomyza. Mol. Phylogenet. Evol. 69:95–108.

Lee Y, Moon SJ, Montell C. 2009. Multiple gustatory receptors required for the caffeine response in Drosophila. Proc. Natl. Acad. Sci. U. S. A. 106:4495–4500.

Leung NY, Thakur DP, Gurav AS, Kim SH, Di Pizio A, Niv MY, Montell C. 2020. Functions of Opsins in Drosophila Taste. Curr. Biol. 30:1367–1379.e6.

Lichtenstein EP, Morgan DG, Mueller CH. 1964. Insecticides in Nature:Naturally Occurring Insecticides in Cruciferous Crops. J. Agric. Food Chem. 12:158–161.

Li Y, Hoffmann J, Li Y, Stephano F, Bruchhaus I, Fink C, Roeder T. 2016. Octopamine controls starvation resistance, life span and metabolic traits in Drosophila. Sci. Rep. 6:35359.

Love MI, Huber W, Anders S. 2014. Moderated estimation of fold change and dispersion for RNA-seq data with DESeq2. Genome Biol. 15:550.

Macpherson LJ, Dubin AE, Evans MJ, Marr F, Schultz PG, Cravatt BF, Patapoutian A. 2007. Noxious compounds activate TRPA1 ion channels through covalent modification of cysteines. Nature 445:541–545.

Matsunaga, T., C. E. Reisenman, B. Goldman-Huertas, S. Rajshekar, H. C. Suzuki, D. Tadres, J. Wong, M. Louis, S. R. Ramírez, N. K. Whiteman. 2025. Odorant receptors tuned to isothiocyanates in Drosophila melanogaster are co-opted and expanded in herbivorous relatives. bioRxiv.

Matsunaga T, Reisenman CE, Goldman-Huertas B, Brand P, Miao K, Suzuki HC, Verster KI, Ramírez SR, Whiteman NK. 2022. Evolution of Olfactory Receptors Tuned to Mustard Oils in Herbivorous Drosophilidae. Mol. Biol. Evol. [Internet] 39. Available from: 10.1093/molbev/msab362

McBride CS. 2007. Rapid evolution of smell and taste receptor genes during host specialization in Drosophila sechellia. Proceedings of the National Academy of Sciences 104:4996–5001.

McBride CS, Arguello JR. 2007. Five Drosophila genomes reveal nonneutral evolution and the signature of host specialization in the chemoreceptor superfamily. Genetics 177:1395–1416.

McBride CS, Baier F, Omondi AB, Spitzer SA, Lutomiah J, Sang R, Ignell R, Vosshall LB. 2014. Evolution of mosquito preference for humans linked to an odorant receptor. Nature 515:222–227.

Montell C. 2021. Drosophila sensory receptors—a set of molecular Swiss Army Knives. Genetics [Internet] 217. Available from: https://academic.oup.com/genetics/article-abstract/217/1/iyaa011/6128801

Nishida R, Ohsugi T, Kokubo S, Fukami H. 1987. Oviposition stimulants of a Citrus-feeding swallowtail butterfly, Papilio xuthus L. Experientia 43:342–344.

Oh SM, Jeong K, Seo JT, Moon SJ. 2021. Multisensory interactions regulate feeding behavior in Drosophila. Proc. Natl. Acad. Sci. U. S. A. [Internet] 118. Available from: 10.1073/pnas.2004523118

Okamura Y, Dort H, Reichelt M, Tunström K, Wheat CW, Vogel H. 2022. Testing hypotheses of a coevolutionary key innovation reveals a complex suite of traits involved in defusing the mustard oil bomb. Proc. Natl. Acad. Sci. U. S. A. 119:e2208447119.

Ozaki K, Ryuda M, Yamada A, Utoguchi A, Ishimoto H, Calas D, Marion-Poll F, Tanimura T, Yoshikawa H. 2011. A gustatory receptor involved in host plant recognition for oviposition of a swallowtail butterfly. Nat. Commun. 2:542.

Peláez JN, Gloss AD, Goldman-Huertas B, Kim B, Lapoint RT, Pimentel-Solorio G, Verster KI, Aguilar JM, Nelson Dittrich AC, Singhal M, et al. 2023. Evolution of chemosensory and detoxification gene families across herbivorous Drosophilidae. G3 [Internet] 13. Available from: 10.1093/g3journal/jkad133

Peláez JN, Gloss AD, Ray JF, Chaturvedi S, Haji D, Charboneau JLM, Verster KI, Whiteman NK. 2022. Evolution and genomic basis of the plant-penetrating ovipositor: a key morphological trait in herbivorous Drosophilidae. Proc. Biol. Sci. 289:20221938.

Prieto-Godino LL, Rytz R, Cruchet S, Bargeton B, Abuin L, Silbering AF, Ruta V, Dal Peraro M, Benton R. 2017. Evolution of Acid-Sensing Olfactory Circuits in Drosophilids. Neuron 93:661–676.e6.

Ritz C, Baty F, Streibig JC, Gerhard D. 2015. Dose-Response Analysis Using R. PLoS One 10:e0146021.

Ryuda M, Calas-List D, Yamada A, Marion-Poll F, Yoshikawa H, Tanimura T, Ozaki K. 2013. Gustatory Sensing Mechanism Coding for Multiple Oviposition Stimulants in the Swallowtail Butterfly, Papilio Xuthus. Journal of Neuroscience 33:914–924.

Saito S, Saito CT, Igawa T, Takeda N, Komaki S, Ohta T, Tominaga M. 2022. Evolutionary Tuning of Transient Receptor Potential Ankyrin 1 Underlies the Variation in Heat Avoidance Behaviors among Frog Species Inhabiting Diverse Thermal Niches. Mol. Biol. Evol. 39:msac180.

Sato S, Magaji AM, Tominaga M, Sokabe T. 2023. Avoidance of thiazoline compound depends on multiple sensory pathways mediated by TrpA1 and ORs in Drosophila. Front. Mol. Neurosci. 16:1249715.

Scott K. 2018. Gustatory Processing in *Drosophila melanogaster*. Annu. Rev. Entomol. 63:15–30.

Shiraiwa T. 2008. Multimodal chemosensory integration through the maxillary palp in Drosophila. PLoS One 3:e2191.

Shiraiwa T, Carlson JR. 2007. Proboscis extension response (PER) assay in Drosophila. J. Vis. Exp.:193.

Stephens M. 2017. False discovery rates: a new deal. Biostatistics 18:275–294.

Talavera K, Startek JB, Alvarez-Collazo J, Boonen B, Alpizar YA, Sanchez A, Naert R, Nilius B. 2020. Mammalian Transient Receptor Potential TRPA1 Channels: From Structure to Disease. Physiol. Rev. 100:725–803.

Thorvaldsdóttir H, Robinson JT, Mesirov JP. 2013. Integrative Genomics Viewer (IGV): high-performance genomics data visualization and exploration. Brief. Bioinform. 14:178–192.

Tracey WD, Wilson RI, Laurent G, Benzer S. 2003. painless, a Drosophila gene essential for nociception. Cell 113:261–273.

Valencia-Montoya WA, Pierce NE, Bellono NW. 2024. Evolution of sensory receptors. Annu. Rev. Cell Dev. Biol. 40:353–379.

Wallbank BE, Wheatley GA. 1979. Some responses of cabbage root fly (Delia brassicae) to allyl isothiocyanate and other volatile constituents of crucifers. Ann. Appl. Biol. 91:1–12.

Wang W, Dweck HKM, Talross GJS, Zaidi A, Gendron JM, Carlson JR. 2022. Sugar sensation and mechanosensation in the egg-laying preference shift of Drosophila suzukii. Elife 11:e81703.

Weiss LA, Dahanukar A, Kwon JY, Banerjee D, Carlson JR. 2011. The molecular and cellular basis of bitter taste in Drosophila. Neuron 69:258–272.

Wheat CW, Vogel H, Wittstock U, Braby MF, Underwood D, Mitchell-Olds T. 2007. The genetic basis of a plant insect coevolutionary key innovation. Proceedings of the National Academy of Sciences 104:20427–20431.

Whiteman NK, Gloss AD, Sackton TB, Groen SC, Humphrey PT, Lapoint RT, Sønderby IE, Halkier BA, Kocks C, Ausubel FM, et al. 2012. Genes involved in the evolution of herbivory by a leaf-mining, drosophilid fly. Genome Biol. Evol. 4:900–916.

Whiteman NK, Groen SC, Chevasco D, Bear A, Beckwith N, Gregory TR, Denoux C, Mammarella N, Ausubel FM, Pierce NE. 2011. Mining the plant-herbivore interface with a leafmining Drosophila of Arabidopsis. Mol. Ecol. 20:995–1014.

Wittstock U, Agerbirk N, Stauber EJ, Olsen CE, Hippler M, Mitchell-Olds T, Gershenzon J, Vogel H. 2004. Successful herbivore attack due to metabolic diversion of a plant chemical defense. Proc. Natl. Acad. Sci. U. S. A. 101:4859–4864.

Yang J, Guo H, Jiang N-J, Tang R, Li G-C, Huang L-Q, van Loon JJA, Wang C-Z. 2021. Identification of a gustatory receptor tuned to sinigrin in the cabbage butterfly Pieris rapae. PLoS Genet. 17:e1009527.

Yapici N, Cohn R, Schusterreiter C, Ruta V, Vosshall LB, Yapici N, Cohn R, Schusterreiter C, Ruta V, Vosshall LB. 2016. Taste Circuit that Regulates Ingestion by Integrating Food and Hunger Signals Article A Taste Circuit that Regulates Ingestion by Integrating Food and Hunger Signals. Cell 165:715–729.

Zhao J, Lin King JV, Paulsen CE, Cheng Y, Julius D. 2020. Irritant-evoked activation and calcium modulation of the TRPA1 receptor. Nature 585:141–145.

Zhong L, Bellemer A, Yan H, Honjo K, Robertson J, Hwang RY, Pitt GS, Tracey WD. 2012. Thermosensory and Nonthermosensory Isoforms of Drosophila melanogaster TRPA1 Reveal Heat-Sensor Domains of a ThermoTRP Channel. Cell Rep. 1:43–55.

